# Cerebellar α1_D_-adrenergic receptors mediate stress-induced dystonia in tottering^tg/tg^ mice

**DOI:** 10.1101/2023.11.12.566757

**Authors:** Pauline Bohne, Mareike Josten, Lina Rambuschek, Xinran Zhu, Max O. Rybarski, Melanie D. Mark

**Author notes:** Correspondence: Dr. Melanie D. Mark.

## Abstract

Episodic ataxia type 2 (EA2) is an inherited neurological disorder, where patients suffer from chronic ataxia and severe episodes of motor dysfunction exhibited as dystonia. Despite other factors, physical and emotional stress triggers those episodes reliably in both human and mice. We used the well-established EA2 mouse model tottering to explore the cerebellar adrenergic receptor (AR) involvement in stress-induced dystonic attacks. We found that α1-ARs, but not α2-ARs, on cerebellar Purkinje cells (PCs) are activated by norepinephrine (NE) from the locus coeruleus (LC), differentially expressed and required for initiation of dystonia. Moreover, pharmacological blockade and shRNA-induced knock down of cerebellar α1_D_-ARs was sufficient to effectively prevent stress-induced dystonia in homozygous tottering^tg/tg^ mice but had no impact on ataxia amelioration. *In vivo* recordings and live calcium (Ca^2+^) imaging of PCs demonstrated that α1_D_-AR blockade successfully protects PCs from NE-mediated erratic firing patterns through decreased release of calcium from intracellular stores, thus preventing stress-induced dystonia. Furthermore, chemogenetic inhibition of the LC-NE pathway alleviated the frequency and symptoms of stress-induced dystonia. Together, our data show the modulatory effects of NE on dystonia severity and suggest a predominant role of cerebellar α1_D_-ARs in the formation of stress-induced dystonia in tottering^tg/tg^ mice and, thereby providing a potential new therapeutic target to treat stress-induced dystonia in EA2.

## Introduction

Episodic Ataxia Type 2 (EA2) is a rare, autosomal inherited episodic neurological disorder belonging to a broad family of channelopathies premised on mutations in ion channels. EA2 is caused by loss-of-function mutations in the CACNA1A gene, encoding the pore-forming α1 subunit of the P/Q-type voltage-gated calcium channel (Ca_V_2.1) (*1*), which is predominantly expressed on cerebellar Purkinje cells (PCs) and granule cells (GCs) and required for neurotransmitter release (*2*, *3*). EA2 patients suffer from permanent ataxia, nausea, nystagmus, cognitive deficits and episodes of increasing ataxia, instability and dystonia due to involuntary antagonistic muscle contraction (*4*). Episodic attacks occur spontaneously or can be triggered by chemical stressors such as caffeine and ethanol, as well as by physical and psychological stressors (*5*). Since these episodic attacks can last hours to days, patients are advised to remain relaxed by controlling their environment (*6*), however, this is accompanied with lifetime limitations.

Similar to EA2 patients, tottering^tgtg^ mice carry a spontaneous loss-of-function mutation (P601L) in the CACNA1A gene, resulting in a ~ 40% reduction of Ca^2+^ currents through the P/Q-type channel (*7*), thereby resembling symptoms of EA2 patients including ataxia and episodes of severe paroxysmal nonkinesigenic dyskinesia. Stress, ethanol and caffeine reliably elicit attacks of motor dysfunction in tottering^tg/tg^ (*8–12*), making them an ideal animal model to study the neuronal basis of EA2 (*13*). During stressful conditions, norepinephrine (NE) is released from the LC, innervating the whole brain. In the cerebellum, PCs are the main target of LC synapses (*14*), receiving predominantly inhibitory input though α- and β-adrenergic receptors (ARs) (*15–17*), resulting in long-lasting inhibition of PCs (*18*).

Tottering^tg/tg^ mice are NE hyperinnervated (*19*), and pharmacological blockade of cerebellar α1-ARs was effective in preventing stress-induced dystonia (*20*), though the exact mechanism was not understood until recently. Snell and colleagues showed that NE-mediated stress-induced attacks of motor dysfunction are caused by avid burst firing of cerebellar PCs via activation of α1-ARs and consequent reduction of SK channel activity (*21*). However, systemic administration of general α1-ARs antagonists targets ARs in the entire body, resulting in unwanted side effects. Here, we utilized the well-characterized EA2 mouse model tottering^tg/tg^ to identify a specific α1-AR subtype predominantly involved in formation of stress-mediated paroxysmal dyskinesia. We found that the α1_D_-AR specific antagonist BMY-7378 (BMY) successfully alleviated stress-induced dystonia and recovered PC simple spike (SS) firing after NE-mediated inhibition. Cerebellar α1_D_-ARs were also upregulated compared to tottering^−/−^ control mice and chronic infusion of BMY and shRNA-induced knock-down of α1_D_-ARs in the cerebellar vermis of tottering^tg/tg^ mice reduced or prevented dystonic attacks and severity, verifying the cerebellar α1_D_-AR involvement in the formation of stress-induced paroxysmal dyskinesia. *In vivo* Ca^2+^ imaging of PCs using miniscopes showed an increase in intracellular calcium release at the onset of dystonic attacks, which was abolished after application of BMY-7378, hinting to impaired calcium homeostasis as the cause for dystonic episodes. We further explored the circuitry responsible for NE-mediated stress-induced dystonia by chemogenetically silencing LC neurons using inhibitory DREADDS (designed receptors exclusively activated by designer drugs) (*22*). Chemogenetic silencing of LC neurons, which mainly synapse on PCs, alleviated stress-induced dystonia, while silencing of cerebellar interneurons failed to do so, strengthening the selective involvement of PCs in dystonia (*8*, *23*).

Our findings provide further insight into the noradrenergic modulation of the cerebellar circuitry in stress-induced paroxysmal dystonia in tottering^tg/tg^ mice. More importantly, we identified cerebellar α1_D_-ARs as the accountable adrenergic receptor for the formation of dystonia, providing a new therapeutic target to treat E2 patients.

## Results

### Upregulated α1_D_ adrenergic receptors located on cerebellar PCs mediate stress-induced paroxysmal dystonia in *tottering^tg/tg^* mice

Noradrenergic blockade through α1-ARs was shown to prevent stress-induced attacks in the EA2 mouse model tottering^tg/tg^ (*20*, *21*), however it is not clear, whether and to what extend the specific α1-AR subtypes contribute to the formation of stress-induced dystonia. To investigate this, we initially verified the reported predominant contribution of α1-ARs, but not α2-ARs in stress-induced motor dysfunction (*20*) (Figure 1A). Indeed, injection of the α1-AR antagonist prazosin hydrochloride (Praz) 30 min prior to cage change stress significantly alleviated frequency of stress-induced motor dysfunction in tottering^tg/tg^ mice but did not affect the onset or duration of dystonia in animals that still underwent and dystonic episode. Strikingly, permanent ataxia reported in both EA2 patients (*5*, *24*) and tottering^tg/tg^ mice (*10*) was not improved in presence of Praz (Table S1). Furthermore, injection of the α2-AR antagonist yohimbine hydrochloride (Yoh) did not affect the frequency or duration of stress-induced attacks, it did show a trend in delaying attack onset (Figure 1A), thereby confirming the postulated predominant role of α1-ARs in stress-induced dystonia.

**Figure 1:**
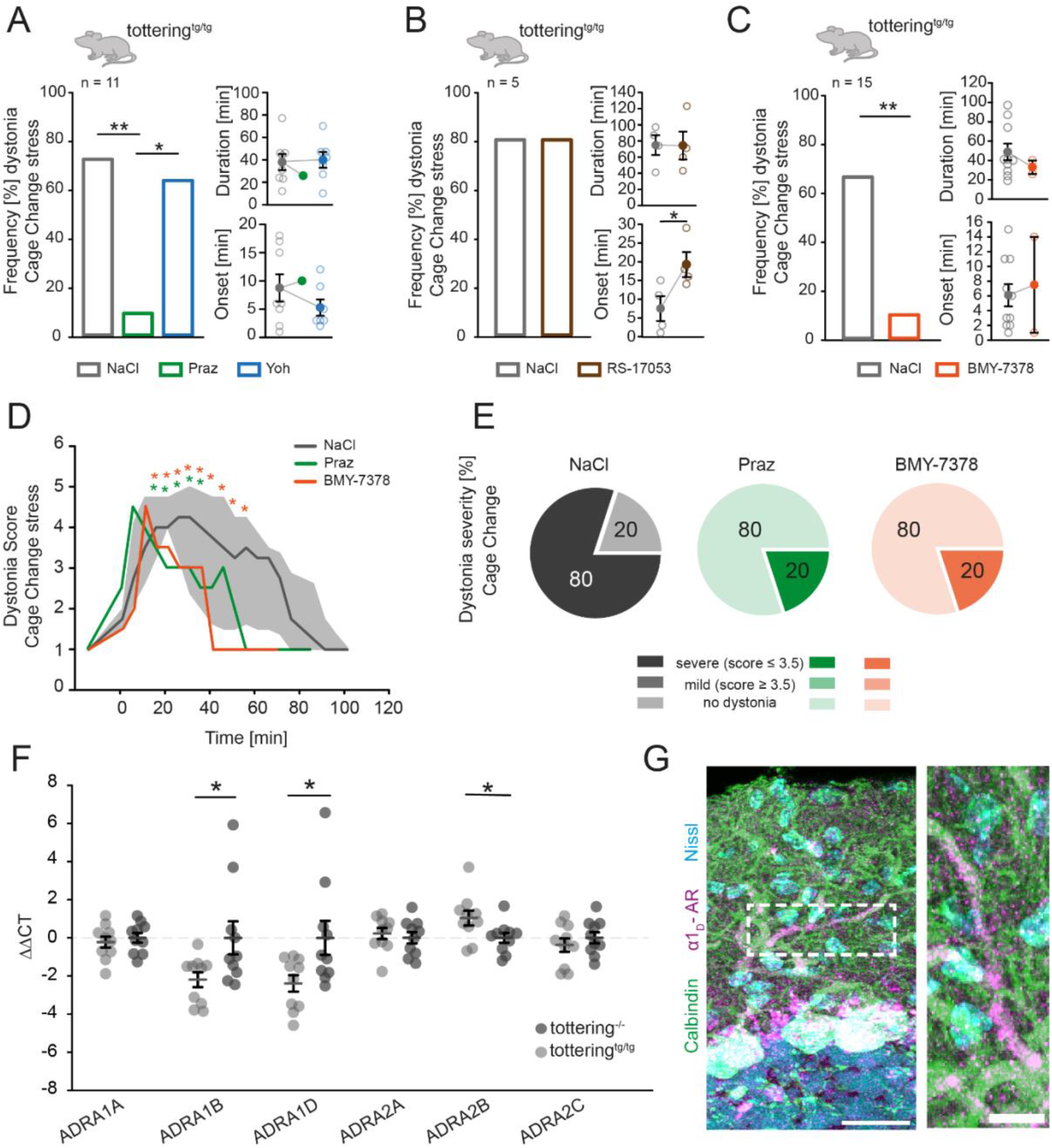
α1 adrenergic receptor antagonists prevent stress-induced dystonia in tottering^tg/tg^ mice. Efficiency of adrenoreceptor antagonists in blocking stress-induced dystonic attacks in homozygous tottering mice were evaluated using the cage change stress paradigm. **(A)** 72% of vehicle (NaCl) injected tottering^tg/tg^ mice (n=11) respond with paroxysmal dystonia to the stress-test starting after 8.7±2.4 min and 37.8±7.1 min duration. Injection of 10 mg/kg prazosin hydrochloride (Praz), a general α1 adrenergic receptor antagonist 30 min prior to the test, reduced the frequency of mice displaying dystonia significantly to 9% (p=0.003). Only one attack was observed after 10 min with a duration of 27min. Injections of 20 mg/kg yohimbine hydrochloride (Yoh), an α2 adrenoreceptor antagonist, did neither reduce frequency of stress-induced dystonia (64%, p=0.684), nor influenced onset (5.2±1.4min, p=0.256) or duration (39.9±7min, p=0.847) of attacks. (**B)** The α1_A_-AR antagonist RS-17053 did neither alleviate stress-induced dystonia (80% vehicle, 80% RS, p=1), nor influenced dystonia duration (74.75±12.195 NaCl, 74.25±17.24 RS, p=0.982), but delayed onset of attacks from 7.5±3.304 min to 19.25±3.351min (p=0.047) in tottering^tg/tg^ mice. **(C)** Injections of 10 mg/kg BMY-7378 dihydrochloride, a subtype specific α1_D_-AR antagonist significantly reduced the frequency of dystonia to 13% compared to vehicle injected tottering^tg/tg^ mice (66%, p=0.004), while duration (48.8±8.6min NaCl, 33±7min BMY, p=0.451) and onset (6.1±1.5min NaCl, 7.5±6.5min BMY, p=0.745) were not affected. **(D)** Both Praz and BMY-7378 alleviated severity of stress-induced dystonic attacks when they occurred compared to NaCl injected mice starting 15 min after exposure to stress. **(E)** 80% of mice displayed severe motor symptoms undergoing stress-induced dystonia (score≤3.5) after NaCl application and cage change stress, while 20% did not display an attack (score≥3.5). Both Prazosin and BMY increased the number of mice showing no dystonia to 80%, while 20% still displayed severe motor impairments. **(F)** Analysis of the AR subtypes mRNA levels in the cerebella of both control and tottering^tg/tg^ mice via qPCR reveals that α1_B_- and α1_D_-AR mRNA are significantly upregulated in tottering^tg/tg^ compared to control mice (p=0.038; p=0.045, respectively). α1_A_-ARs (p=0.564), α2_A_-ARs (p=0.592) and α2_C_-ARs (p=0.410) are not differentially expressed in tottering^tg/tg^ compared to tottering^−/−^ control mice. However, α2_B_-AR mRNA is downregulated (p=0.04). **(G)** IHC staining against the α1_D_-AR (magenta) shows expression on the PC (green) dendritic tree. Scale bars: 25 µm (left) and 15 µm (right). See table S6 for data and p-values. Data are presented as mean±SEM or median±75%/25% quartiles (D). Statistical significance was evaluated with student’s t-test or Mann-Whitney Rank Sum test or Two Way Repeated Measure ANOVA. (*p≤0.05, **p≤0.01, ***p≤0.001).

Next, we injected tottering^tg/tg^ mice with AR specific antagonist to further dissect the involvement of the three α1-AR subtypes in stress-induced paroxysmal dystonia. Mice were injected with either RS-17053, an α1_A_-AR specific antagonist (Figure 1B), or BMY-7378, an α1_D_-AR specific antagonist (Figure 1C) (*25*) 30 min prior to a cage change. Block of α1_A_-Ars had no impact on the probability of stress-induced dystonia or duration of attacks, but significantly delayed the onset of attacks in response to the stress test (Figure 1B). By contrast, block of α1_D_-ARs effectively prevented stress-induced dystonic episodes in tottering^tg/tg^ mice and showed tendencies to reduce attack duration (Figure 1C). In tottering^tg/tg^ mice, episodes of dystonic attacks follow a characteristic uniform progression starting with extension of their hind limbs, followed by an overarched back, twitching forelimbs and a stiff neck before reaching the head, which results in prolonged immobility of the mouse (*9*, *10*). This uniformity allowed us to score the severity of dystonic attacks using a modified scale (*26*) (Figure 1D, E). Those few dystonic attacks that still proceeded in presence of Praz or BMY-7378, reached identical severities to attacks caused by NaCl injection in the first 20 min, but showed a significant milder course and attack duration (Figure 1D, E). These data strongly suggest a predominant role for α1_D_-ARs in the formation of stress-induced dystonia.

To further strengthen this hypothesis, we analyzed cerebellar AR expression. A study performed in the late 80s found slight overall α_1_-AR upregulation in the cerebellum of tottering^tg/tg^ mice via receptor audiography with [^3^H]prazosin (*27*). However, the individual AR subtypes were not further characterized. Additionally, AR localisation on different cell types within the cerebellum yielded different results, depending on the method used and species analysed (*28–30*). For α1_D_-ARs, a study in humans found expression restricted to PCs and DCN (*30*). However, adrenergic receptor expression may be different in EA2 patients and tottering^tg/tg^ mice, which could partially explain their increased susceptibility to stressors. To identify the different AR subtypes in the cerebellum, we performed qPCR of cerebellar ARs (Figure 1F) and conducted supporting IHC localization staining (Figure S1). mRNA analysis revealed altered AR expression in tottering^tg/tg^ compared to control mice. Strikingly, cerebellar α1_D_-AR mRNA was significantly upregulated in tottering^tg/tg^ mice, as well as α1_B_-AR mRNA, which is needed for cell surface expression of α1_D_-ARs *in vitro* (*31*). α2_B_-ARs displayed a reduced expression in tottering^tgtg^ compared to control littermates (see Table S6 for data and p-values). Staining against the α1_D_-AR subtype displays localization on the PC dendritic tree (Figure 1G), possibly postsynaptic, which was not observed for the other subtypes (Figure S1). Taken together, our data suggest that α1_D_-AR may primarily be located at the PC dendritic tree and may play a role in mediating stress-induced dystonia in tottering^tg/tg^ mice.

### *In vivo* microinjection of the α1_D_-adrenoreceptor antagonist BMY-7378 rescues norepinephrine-induced inhibition of cerebellar Purkinje cells

Release of NE was previously shown to inhibit PC firing (*32*) and induce erratic PC firing in tottering^tg/tg^ mice (*21*). The loss of timely precise PC pacemaking is suggested to be the pivotal factor in cerebellar dysfunction (*33*). We therefore explored the direct effects of α1_D_-AR inhibition on PC activity by electrophysiological *in vivo* recordings (Figure 2). Initial extracellular *in vivo* recordings of PCs in both control tottering^−/−^ and tottering^tg/tg^ mice confirmed that PC firing rates are not decreased in tottering^tg/tg^ mice, but display increased irregularity in SS firing compared to control mice (*34*) (Figure S3; Table S7). The frequency of complex spikes (CS) was significantly reduced compared to control tottering^−/−^ mice, suggesting altered climbing fiber (CF) synaptic innervation in tottering^tg/tg^ mice (Figure S3C, Table S7). Since CF inputs trigger synaptic plasticity at both the PC dendrites and independently generate a strong distal output signal in the PC axon (*35*), this decrease may hint to PC coding deficits in tottering^tg/tg^ mice. When we hindered PC firing of tottering^tg/tg^ mice by pressure injection of 20 µM NE, a dose previously reported to activate α1-ARs in PCs (*36*) (Figure S4). We observed a 53% decrease of PC SS firing (Figure S4C), increased irregularity of PC SS firing (Figure S4E) and a 53% reduction of CS which did not recover during recordings (Figure S4F, J). Therefore, our *in vivo* NE application resembles PC irregularity observed during and identified as the cause for caffeine (*37*) and stress-.induced motor attacks (*21*).

**Figure 2:**
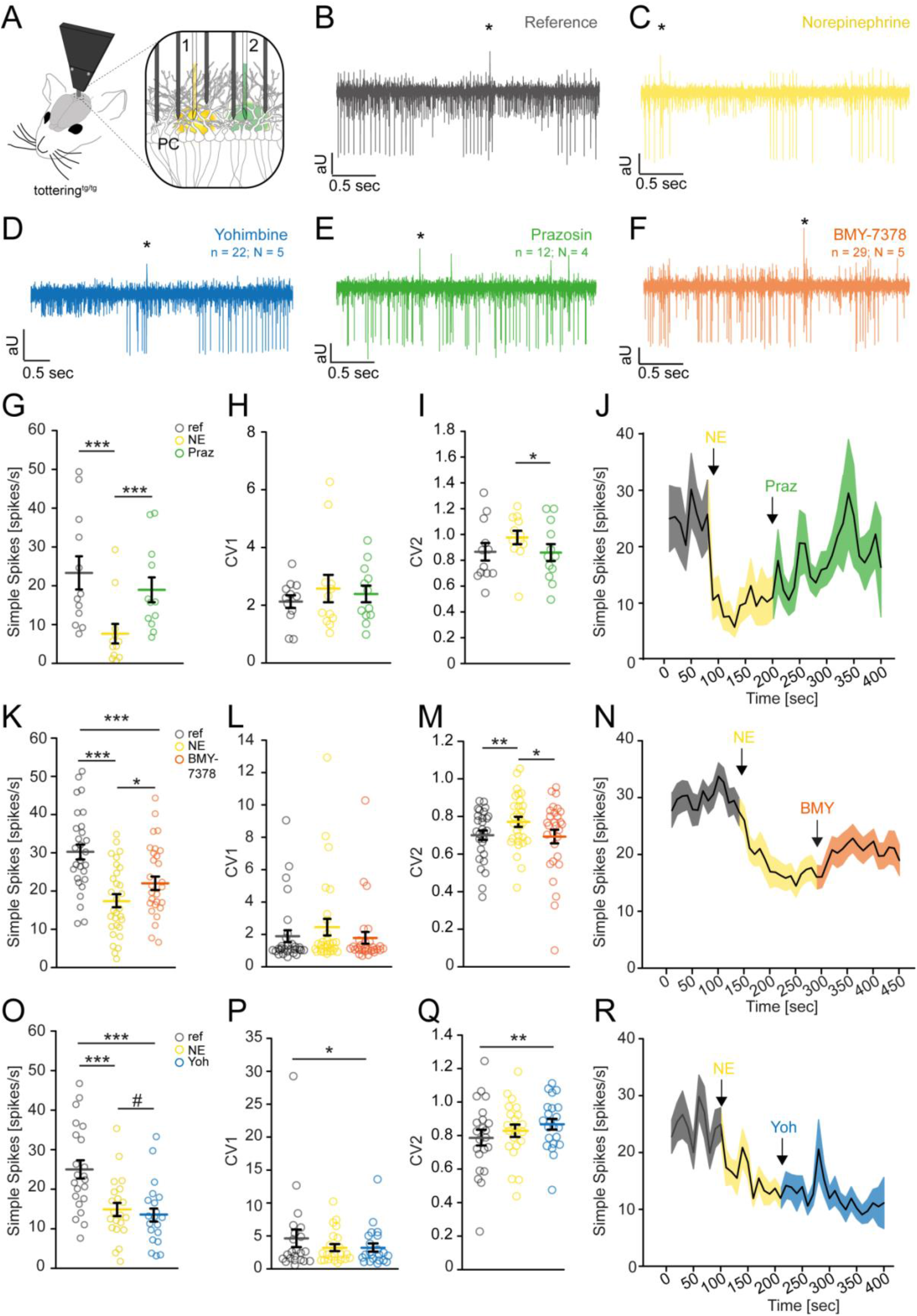
The general α1AR antagonist Prazosin hydrochloride and subtype-specific α1_D_ AR antagonist BMY-7378 restore Purkinje cell firing *in vivo* in tottering^tg/tg^ mice. **(A)** Schematic of the *in vivo* recording setup. NE (1) and AR blockers (2) were pressure injected via microinjection pipettes and PC activity was simultaneously recorded via 5 electrodes. Example traces of spontaneous PC activity **(B)**, and after application of NE **(C**), YOH (**D**), Praz (**E**) and BMY-7378 (**F). (G-J)** Praz application recovers PC SS firing after NE-mediated inhibition (7.65±2.51 vs 18.942±3.209, p≤0.001) and restores CV2 (0.977±0.0524 vs 0.860±0.0657, p=0.034) **(I),** while the CV1 remains unaltered (2.574±0.476 vs 2.388±0.288, p=0.741) **(H)**. **(K-N)** The α1_D_-AR subtype specific blocker BMY alleviates NE-mediated inhibition of PC SS firing (17.486±1.711 vs 22.031±1.77, p=0.013). **(L)** NE (2.449±0.512) and BMY (1.786±0.363) did not influence PC firing regularity, though NE significantly increased PC irregularity (0.772±0.0268, p=0.002), which was recovered after BMY application (0.693±0.0362, p=0.015). **(M).** The α2-AR blocker YOH did not recover PC SS firing after NE-mediated inhibition (14.882±1.657 vs 13.464±1.642, p=0.055) **(O),** nor did application of the drug impact CV1 (3.229±2.516 vs 3.241±0.622, p=0.194) **(P)** but increased CV2 compared to reference recordings (0.788±0.0468 vs 0.868±0.0322, p=0.006) **(Q). (R)** Average traces of PC SS after NE and YOH application. See table S6 for data and p-values. Data are presented as mean±SEM. Statistical significance was evaluated by paired t-test. (*p≤0.05, **p≤0.01, ***p≤0.001).

We next dissected the effects of α1-AR blockers on NE-mediated inhibition of PCs (Figure 2A-F) (*17*, *32*, *38*). The general α1-AR blocker Praz reliably recovered PC SS firing to 81% of reference levels after NE-mediated inhibition (Figure 2E, G-J) and successfully restored PC regularity (Figure 2I). Since the α1_D_-AR specific antagonist BMY-7378 reduced the frequency and severity of dystonia in tottering^tg/tg^ mice, we investigated the putative protective effects of BMY-7378 on PC pacemaking (Figure 2F, K-N). The dose of BMY was established beforehand in control animals (Table S3). Although less effective than Praz, BMY-7378 injection after NE-mediated inhibition still recovered PC SS firing by 72% (Figure 2K, N) and also successfully recovered PC regularity to reference levels (Figure 2M). These data indicate that NE decreases PC regularity predominantly through α1_D_-ARs. CS were not altered by adrenergic receptor blockers (Figure S5).

To minimize the chance of measuring false positive effects from the pressure injection, we performed control recordings using the α2-AR antagonist Yoh (Figure 2O-R). Following NE-induced PC SS inhibition, subsequent Yoh injection neither recovered, nor further decreased PC SS firing in tottering^tg/tg^ mice (Figure 2O) but did increase PC irregularity (Figure 2Q). These results are in accordance with our behavior data (Figure 1A).

Taken together, our recording data suggests that NE alters both the pacemaking frequencies and regularity of PCs in tottering^tg/tg^ mice via α1_D_-ARs. Major influences by β-ARs on PC firing have been ruled out (*21*). Since NE has been shown to have bimodulatory effects on PC firing (*36*) through activation of β-ARs on molecular layer interneurons (MLIs) resulting in GABAergic potentiation (*16*, *32*), we cannot rule out secondary effects of BMY-7378 on other cerebellar neurons that help to recover PC activity.

### Chronic delivery of BMY-7378 in the cerebellum of tottering^tg/tg^ mice abolishes stress-induced dystonia

Based on our data, the α1_D_-AR expressed in the cerebellum may be a promising therapeutic target to alleviate stress-induced dystonia. However, our data does not specifically suggest that the cerebellum and cerebellar ARs are involved in generation of dystonia. Several studies have shown a correlative link of episodic motor dysfunction to cerebellar dysfunction or disruptions of PC firing (*23*, *37*, *39–41*). Since the cerebella of tottering^tg/tg^ mice are LC-NE hyperinnervated (*19*) and optogenetic activation of the LC in tottering^tg/tg^ mice is sufficient to provoke episodes of motor dysfunction (*21*), blocking specifically cerebellar α1_D_-ARs should prevent or at least decrease the probability of stress-induced dystonia. To test this hypothesis, we chronically infused BMY-7378 or vehicle alone directly into the cerebellum of tottering^tg/tg^ mice using osmotic pumps (*33*, *37*) (Figure 3). Five days after surgery, BMY-7378 infused mice manifested fewer stress-induced dystonic attacks coupled with a delayed onset and decreased duration compared to vehicle infused mice (Figure 3B, D). Strikingly, dystonic attacks were completely abolished after 7 days of drug infusion lasting another 7 days until depletion of the pumps, while vehicle infused mice permanently displayed stress-induced dystonia (Figure 3B). After depletion of the pump the frequency, onset and duration of dystonic episodes returned to pre-surgery levels (Figure 3B-D), thereby eliminating cerebellar damage as a contributing factor to preventing episodic dystonic attacks (*42*). Vehicle infused mice displayed an accelerated responsiveness to the cage change stress as indicated by reduced onset of attacks (Figure 3C), while their dystonia duration remained unaltered (Figure 3D). Although BMY-7378 infused mice still displayed dystonia 5 days post infusion, scoring of dystonia severity revealed significantly milder attacks (Figure 3E, I), while severity of episodic dystonic attacks of vehicle infused tottering^tg/tg^ mice were not diminished (Figure 3E-J). Dystonia severity increased again after pump depletion in BMY-7378 infused tottering^tg/tg^ mice (Figure 3H). Collectively, these data verify that cerebellar α1_D_-ARs are the predominant AR subtype involved in the formation of stress-induced dystonia in tottering^tg/tg^ mice, providing compelling evidence that antagonizing α1_D_-ARs decrease the frequency and severity of dystonic episodes.

**Figure 3:**
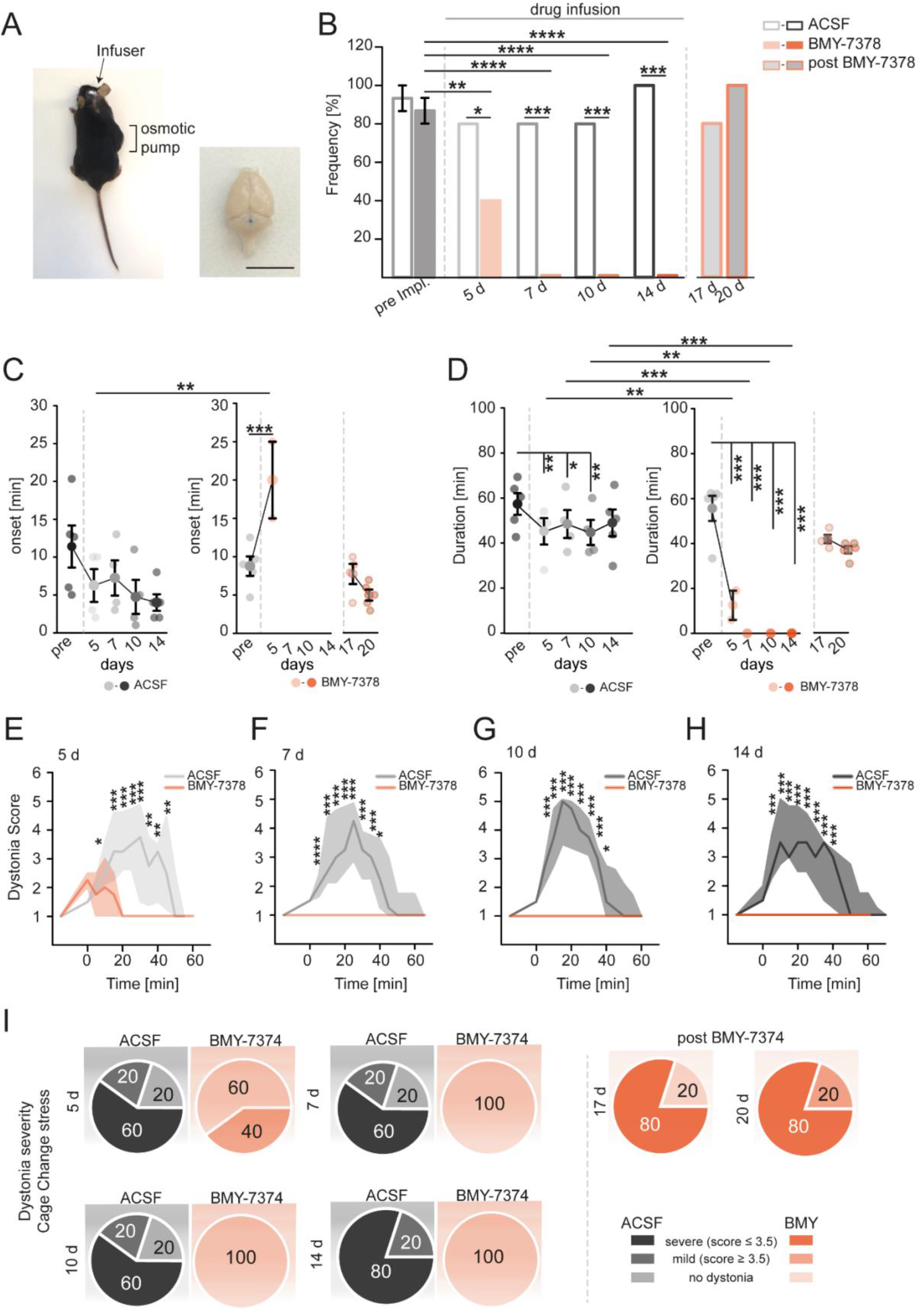
Intracerebellar chronic infusion of BMY-7378 alleviates stress-induced paroxysmal dystonia in tottering^tg/tg^ mice. **(A)** The osmotic pump was implanted subcutaneously at the animals’ right side, a tubing connected the pump to the infuser, allowing chronic intracerebellar infusion of drugs. Canula placement was verified by post-mortem injection of Evans blue stain. **(B)** Intracerebellar chronic delivery of the α1_D_-AR antagonist BMY-7374 effectively prevented stress-induced dystonic attacks starting 5 d after surgery by 50% compared to ACSF infused mice and were completely abolished after 7 d (each n=5). Dystonic attacks returned after 17 d, when infusion of the drug stopped, eliminating cerebellar damage as contributing factor to preventing dystonia. **(C)** The onset of dystonia in ACSF infused tottering^tg/tg^ mice showed a trend in reduction starting 11.4±2.775 min pre-implantation and 4.0±1.095 min after 14 d of infusion (p=0.435), while for BMY infused mice onset was delayed from 8.766±1.268 min to 20.0±5.0 min after 5 d (p≤0.001) and recovered to pre-implantation levels after infusion of the drug stopped (7.75±1.315 min, 17 d). **(D)** Dystonia duration was significantly reduced from 55.632±5.593 min to 12.5±6.5 min after 5 d (p≤0.001) or completely abolished in α1_D_-AR blocker infused mice (p≤0.001), though dystonia duration recovered to pre-implantation levels after delivery of BMY (42.0±1.871 min, 17 d). ACSF infused mice displayed reduced dystonia duration at some time points tested (57.365±4.809 min pre, 45.25±5.851 min 5d (p=0.003), 48.5±6.198 min 7 d (p=0.012), 44.75±5.558 min 10d (p=0.003)), though there was no overall reducing trend observed. **(E-H)** Comparison of dystonia severity revealed an alleviation of severity already after 5 d of BMY infusion, while ACSF infusion did not alleviate severity at any given time. **(I)** During chronic infusion of ACSF, 60-80% of tottering^tg/tg^ mice displayed severe dystonic attacks, while none of BMY infused animals did. Severe paroxysmal dystonic attacks were observed after BMY infusion stopped (80%). See table S6 for data and p-values. Data are presented as mean±SEM or median±75%/25% quartiles (E-H). Statistical significance was evaluated by Two-Way repeated measure ANOVA (*p≤0.05, **p≤0.01, ***p≤0.001).

### Cerebellar shRNA-mediated knock-down of α1_D_ adrenergic receptors prevents stress-induced dystonia

To specifically implicate the role of α1_D_-ARs in dystonia, we conditionally knocked-down these receptors in the cerebellum using small hairpin RNAs (shRNAs). Tottering^tg/tg^ mice were stereotaxically injected with either a mixture of AAV2-CMV-eGFP-U6-mAdra1d[shRNA#1-3] (sh-[ADRA1D]) targeting three different α1_D_ mRNA sequences or non-targeting AAV2-CMV-eGFP-U6-scramble (sh-[scramble]), resulting in robust eGFP expression in cerebellar PCs for both constructs (Figure 4B). Starting three weeks post injection, both sh-[ADRA1D] and sh-[scramble] injected mice were tested in the cage change stress paradigm to determine their susceptibility to stress-induced dystonia (Figure 4C). sh-[ADRA1D] injected mice showed a significant reduction of stress-induced dystonia after three weeks of expression compared to sh-[scramble] injected control tottering^tg/tg^ mice, with a reduction in dystonia duration (Figure 4E). After four weeks, stress-induced motor attacks were prevented completely and mice did not display motor impairments (Movie S1), while sh-[scramble] injected tottering^tg/tg^ mice continued to display stress-induced dystonia without alterations in onset or duration of motor attacks (Figure 4C-E) (Movie S2). Although 20% of sh-[ADRA1D] injected mice experienced dystonia in the third week of expression, severity of attacks was significantly reduced compared to control mice (Figure 4F). Quantitative PCR analysis of sh-[ADRA1D] injected cerebella revealed a significant reduction of α1_D_-AR mRNA levels compared to sh-[scramble] injected tissue (Figure 4G), confirming specificity of targeting α1_D_-AR mRNA. Additional western blot analysis showed decreased levels of α1_D_-AR expression, verifying KO of the receptor in only sh-[ADRA1D] but not sh-[scramble] injected mice (Figure 4H-I).

**Figure 4:**
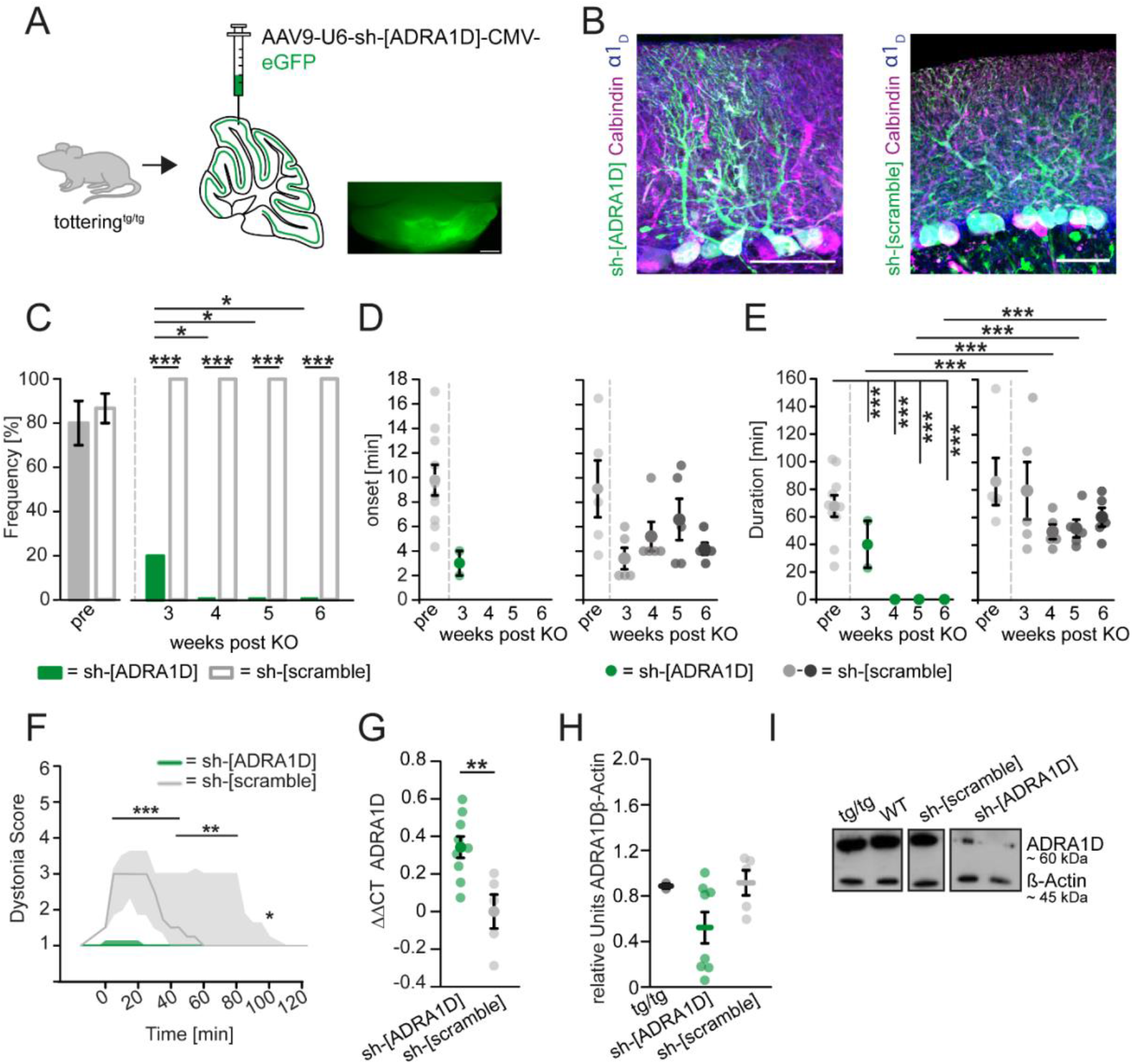
shRNA-mediated knock-down of cerebellar α1_D_ adrenergic receptors prevents stress-induced dystonia. **(A)** Targeting AAV9-U6-sh-[ADRA1D]-CMV-eGFP or non-targeting AAV9-U6-sh-[scramble]-CMV-eGFP were injected into the cerebella of tottering^tg/tg^ mice (n = 10; n = 5, respectively). **(B)** Robust expression of both sh-[ADRA1D] and sh-[scramble] virus (green) in PCs (magenta) after 6 weeks of expression. Scale bars 50 µm. **(C)** After 3 weeks of expression, sh-[ADRA1D] (green) injected mice significantly displayed fewer stress-induced motor attacks than sh-[scramble] control mice (grey 20% vs 100%, p≤0.001). After 4 weeks, stress-induced dystonia was abolished completely, while sh-[scramble] continued to experience attacks (100%). **(D)** Onset of stress-induced dystonia was not statistically analyzed with the Two Way RM ANOVA since sh-[ADRA1D] injected mice did not suffer from dystonia. Onset of attacks was not altered in control mice (p=0.347, One-Way RM ANOVA). **(E)** Compared to pre-injection dystonia duration (67.816±7.782 min), mice with α1_D_-AR knock-down showed significant decrease of attack duration to 40±17 min. No attacks were present after 4 weeks, while attack duration in sh-[scramble] injected mice was not altered. **(F)** sh-[ADRA1D] expressing mice experiencing attacks at 3 weeks post injection had significantly milder dystonia compared to sh-[scramble] injected mice. **(G)** Reduced expression of α1_D_-AR mRNA was found in sh-[ADRA1D] injected mice (green) compared to control mice (grey) (0.343±0.0572 vs 0±0.0903; p=0.006) by qPCR. **(H)** Western blot analysis shows knock-down of the α1_D_-AR (~60kDa) in sh-[ADRA1D] injected mice, while there was distinct protein expression in tottering^tg/tg^ and sh-[scramble] animals. **(I)** Example western blots depicting protein levels of ADRA1D and β-actin in tottering^tg/tg^, WT, sh-[scramble] and in sh-[ADRA1D] injected mice. See table S6 for data and p-values. Data are presented as mean±SEM or median±75%/25% quartiles (F). Statistical significance was evaluated by Two-Way repeated measure ANOVA or Student’s-t test (G, H) (*p≤0.05, **p≤0.01, ***p≤0.001).

Together with our drug studies both electrophysiological and behavioral in mice and shRNA results, these data suggest the involvement of cerebellar α1_D_-AR in stress-induced dystonia and as an effective new therapeutic target to EA2 patients.

### *In vivo* calcium imaging shows a rise of intracellular calcium during early stages of stress-induced dystonia

To explore the effects of BMY-7378 injection on PC firing during stressful events, we performed *in vivo* calcium imaging in tottering^tg/tg^ mice (Figure 5). Mice were injected with a fast GCaMP8m Ca^2+^ sensor into the vermis, followed by immediate implantation of the Inscopix ProView^TM^ Prism lens (*43*, *44*) (Figure 5A). We found that tottering^tg/tg^ still displayed stress-induced dystonic attacks during recordings (Figure 5A) (Movie S3), since damage of the cerebellum was shown to reduce the frequency of dystonia (*42*) with a robust expression of virus only in PCs (Figure 5B). Recordings were performed in both WT and tottering^tg/tg^ mice with both saline and BMY-7378 injection during a cage change test. Control mice displayed regular dendritic ROI activity, which did not change during the duration of recordings independent from the injected agent (Figure 5C-D), while tottering^tg/tg^ mice showed a more irregular activity as indicated by the ROI activity as a measure for frequency (Figure 5C-E) (Movie S4). BMY-7378 injection, however, significantly reduced the ROI activity in tottering^tg/tg^ mice (Figure 5E) (Movie S4), approximating close to the activity observed in control mice. Analysis of the integral (∫) as a measure for released calcium per recorded activity showed an overall decrease in calcium levels with ongoing recordings in tottering^tg/tg^ mice independent from the injected agent (Figure 5G). We exclude decreasing sensitivity of the sensor as a reason for this, since control mice displayed constant levels (Figure 5F), but rather a physiological circumstance based on the general decreased Ca ^2+^ influx through P/Q-type channels in tottering^tg/tg^ mice (*7*). In addition to this, we found a rise of intracellular calcium released at the beginning of a dystonic episode, as indicated by the light grey arrows (Figure 5F-G). With ongoing of the attack, the PCs do not seem to recover well from the erratic burst firing observed during attacks (*21*, *33*), as indicated by the decreasing integral as well as ΔF/F (Figure 5HI).

**Figure 5:**
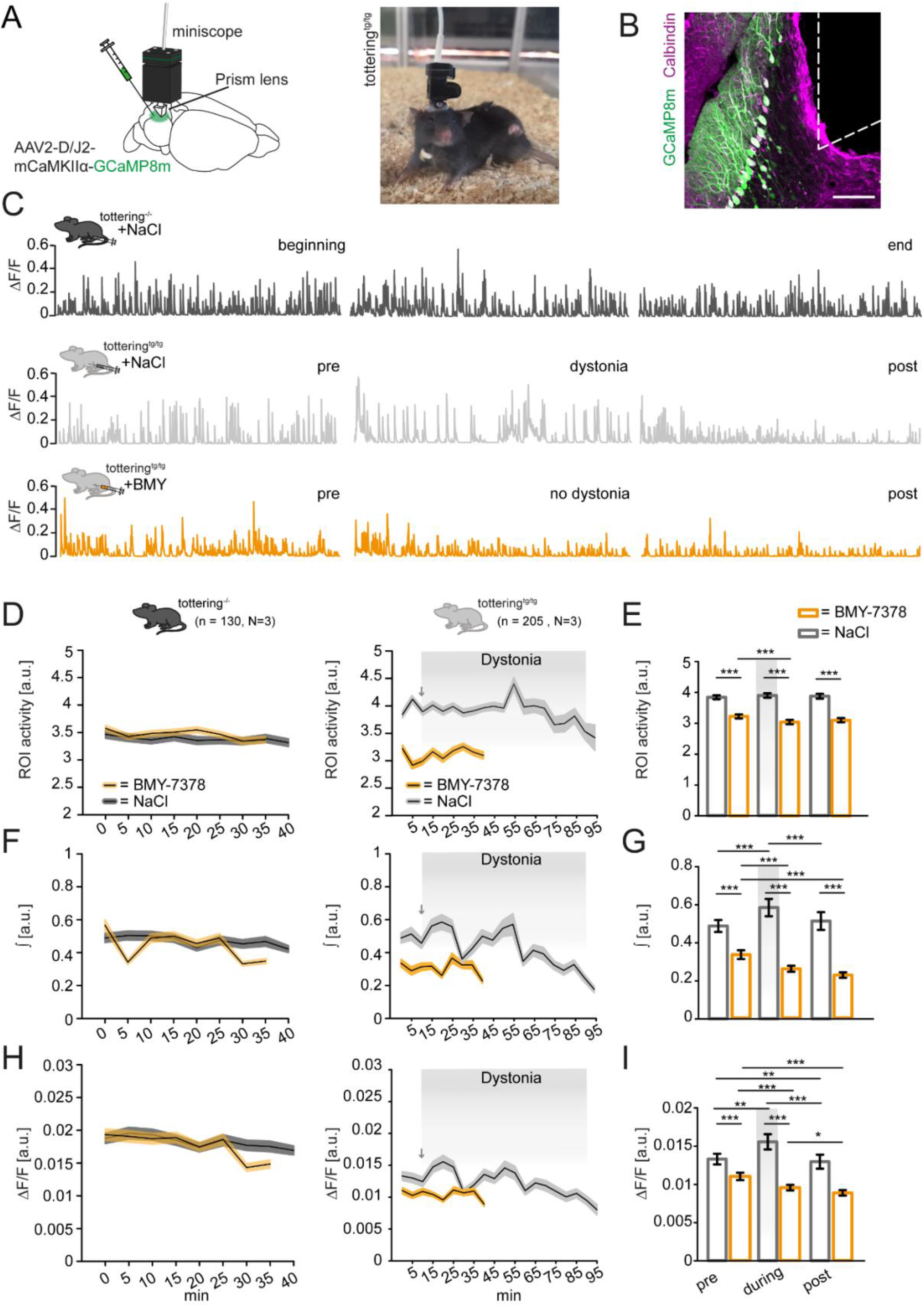
In-vivo calcium imaging of Purkinje cells verifies the involvement of intracellular calcium during stress-induced dystonia in tottering^tg/tg^ mice. **(A)** Scheme of the surgery and image of miniscope (Inscopix) mounted to a tottering^tg/tg^ mouse. **(B)** Example confocal image of AAV2-D/J2-mCaMKIIα-GCaMP8m (green) expression in Purkinje cells (magenta). Dashed lines indicate placement of the prism. Scale bar: 100 µm **(C)** Example traces from ROIs on a Purkinje cell dendrite after ip injection of NaCl in a control mouse (top) and a tottering^tg/tg^ mouse experiencing dystonia (middle). Bottom traces show a ROI recording from a tottering^tg/tg^ mouse after ip injection of BMY-7378 without dystonia. **(D)** Analysis of the ROI activity in tottering^−/−^ control mice display no effects of BMY-7378 ip injection on PC firing frequency. Tottering^tg/tg^ mice display a higher, more irregular ROI activity, which was reduced after ip administration of BMY-7378. **(E)** Comparison of the mean ROI activity in the first recording (0-1 min), forth recording (15-16 min) and last recording (after dystonia ended or in case of no dystonia after 40 min) in NaCl and BMY-7378 injected tottering^tg/tg^ mice. BMY-7378 significantly decreases the mean ROI activity compared to NaCl. **(F)** Analysis of the integral (∫) as a measure of total calcium release during recordings shows that the calcium release is constant in control mice after both NaCl and BMY-7378 ip injection. Tottering^tg/tg^ mice display decreasing calcium levels, independent from the drug injected. **(G)** Comparison of the mean ∫ from the first recording (0-1 min), forth recording (15-16 min) and last recording (after dystonia ended or in case of no dystonia after 40 min) in tottering^tg/tg^ mice shows significantly increased amounts of calcium released at the beginning of a dystonic episode, which normalize once attacks stop. BMY-7378 injection lowers the total amount of released calcium in all recordings. **(H)** Courses of the mean fluorescence intensity (ΔF/F) in control and tottering^tg/tg^ mice shows the overall relative constant fluorescence signals measured in the PC dendritic tree. In tottering^tg/tg^ mice, there is a leap of fluorescence during the early stages of dystonia, indicated by the grey arrow. **(I)** Comparison of the mean ΔF/F signals of the first recording (0-1 min), forth recording (15-16 min) and last recording (after dystonia ended or in case of no dystonia after 40 min) in tottering^tg/tg^ mice shows a significant increase in fluorescence as an indicator for rising calcium levels in the presence of dystonia. Signals decreased under baseline after dystonia is over and mice recovered from the attack. In the presence of BMY-7378 the fluorescence signals decreased, indicating lowering calcium levels. Light grey boxes in the back of graphs indicate presence of dystonia throughout NaCl recordings. See table S6 for data and p-values. Data are displayed as mean±SEM. Statistical significance was evaluated by Student’s-t test or Mann-Whitney Rank Sum test (*p≤0.05, **p≤0.01, ***p≤0.001).

Based on the lower calcium transients measured in tottering^tg/tg^ mice, we assume that their calcium homeostasis is impaired. Calcium mobilization from internal stores is mediated through ryanodine receptors (RyR) and inositol 1,4,5-triphosphate receptors (IP3R) (*45*). The role of intracellular calcium homeostasis of PCs in dystonia has been implicated as RyR only on PCs are required to induce attacks in tottering^tg/tg^ mice (*11*) and interestingly, RyR1 levels are decreased in tottering^tg/tg^ mice (*46*). Our recordings of calcium activity on PC dendritic trees show rises in calcium levels at the beginning of dystonic events. Our findings strengthen the idea of impaired calcium homeostasis during stress-induced dystonia through the Gq-coupled GPCR α1_D_-AR (*47*).

### Chemogenetic silencing of noradrenergic neurons in the locus coeruleus alleviates stress-induced paroxysmal dystonia

Various studies showed that PC SS firing in EA2 mice is impaired and correlates with ataxia and dystonia through interruptions of PC pacemaking (*21*, *33*, *37*, *39*, *48*) NE plays a key role in generation of stress-induced dystonia in tottering^tg/tg^ mice (*9*, *11*, *20*, *21*), and the sole source of NE in the cerebellum is the LC (*49*). LC neurons synapse almost exclusively on PCs (*14*), where binding of NE has a predominant inhibitory effect (*15*, *16*). Its neurons display increased firing rates when mice are confronted with noxious or stressful stimuli (*50*), resulting in a significant release of NE from the LC (*51*) and long-lasting inhibition of PCs (*18*). Consequently, optogenetic stimulation of the LC was shown only to trigger paroxysmal dystonia in tottering^tg/tg^ mice (*21*) and not prevent dystonia by silencing the LC. Therefore, we exploited a chemogenetic approach to prevent NE release from LC synapses during stressful events, likely by maintaining PC pacemaking (Figure 6).

**Figure 6:**
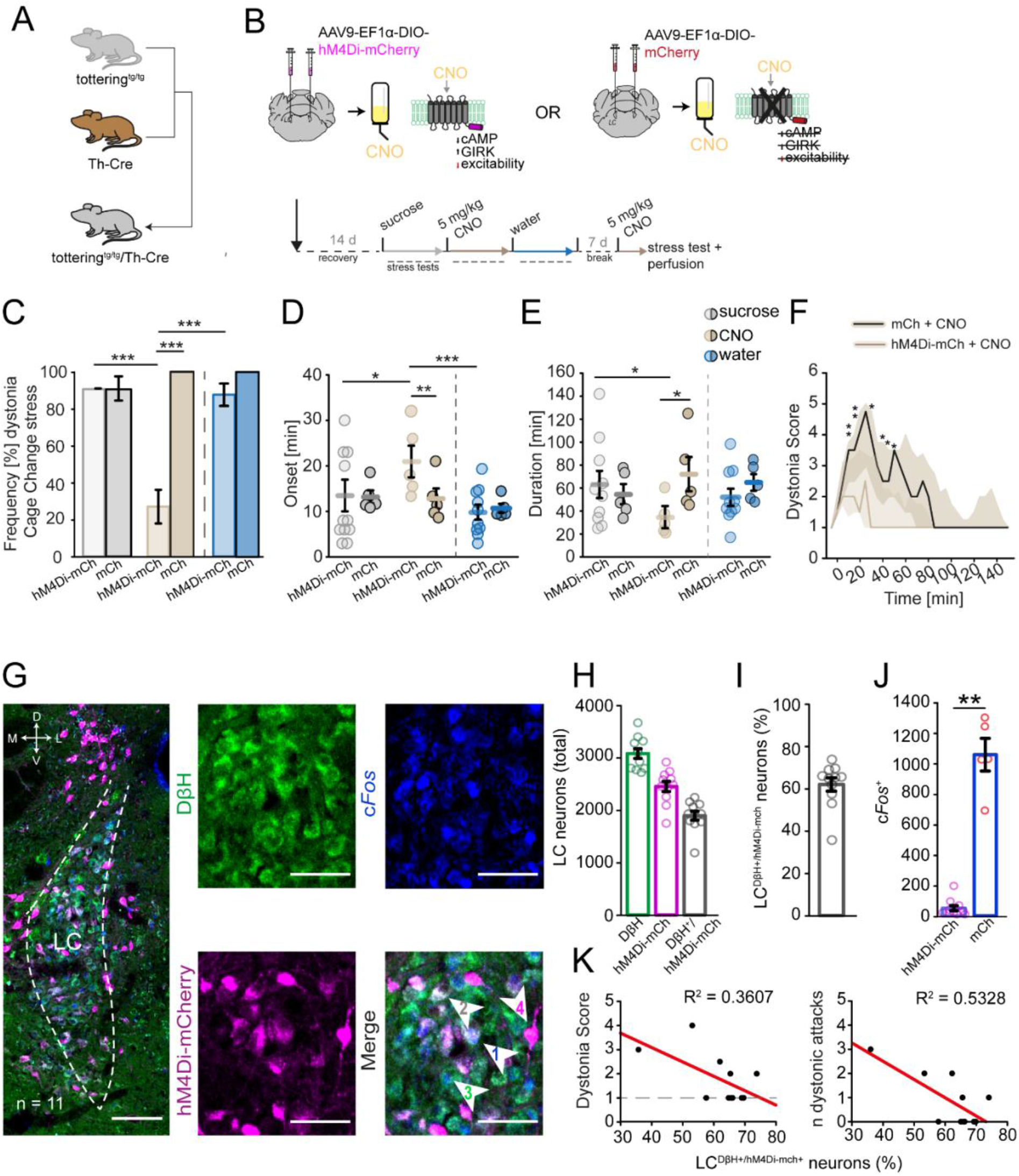
Chemogenetic oral silencing of LC-NE^+^ neurons reduces the frequency, duration and severity of stress-induced dystonic attacks through decreased LC neuronal activity. **(A)** Tottering^tg/tg^/Th-Cre mice were generated by crossing tottering^tg/tg^ mice with homozygous Th-Cre mice. **(B)** Mice were bilaterally injected in the LC with either Cre-dependent AAV9-EF1α-DIO-hM4Di-mCherry or control AAV9-EF1α-DIO-mCherry virus. Both groups received CNO oral administration in their drinking water (2% sucrose solution). Mice were allowed to recover for 14 days prior to testing. Initially, 2% sucrose solution was given *ad libitum,* followed by 5 mg/kg CNO in sucrose solution and water. **(C)** DIO-hM4Di-mCh injected mice showed a significant reduction in stress-induced dystonic episodes when given CNO (27.24±9.12%, p≤0.001) compared to sucrose (90.9%) or water (89.873±6.063%) consumption. DIO-mCh injected mice showed no significant differences in dystonia frequency. **(D)** CNO administration significantly delayed onset of attacks (23.962±3.01 min) compared to sucrose (14.03±1.941 min, p=0.023), water (9.029±1.941, p=0.001) and control mice drinking CNO (6.733±2.626 min, p=0.002) in DIO-hM4Di-mCh injected mice. **(E)** Dystonia duration was significantly reduced in DIO-hM4Di-mCh injected mice when consuming CNO (27.436±10.658 min) compared to control mice (72.133±9.298 min, p=0.025) or drinking sucrose (65.525±6.873 min, p=0.021). **(F)** Comparison of dystonia scores displays shorter attack duration and milder attack progression, if present, in DIO-hM4Di-mCh injected mice consuming CNO compared to DIO-mCh injected tottering^tg/tg^ mice. **(G)** Example image of an AAV9-Ef1α-DIO-hM4Di-mCherry (magenta) injected mice displaying DβH^+^ noradrenergic LC neurons (green) stained for *cFos* (blue). Arrows in the merged image point to either triple positive (DβH^+^/hM4Di-mCherry^+^/*cFos*^+^) LC neurons used for analysis (1), or marginal DβH^+^/hM4Di-mCherry^+^ (2), DβH^+^/*cFos*^+^ (3) or hM4Di-mCherry only (4) expressing LC neurons. Scale bar LC overview: 100 µm. Scale bars in close-ups: 50 µm. **(H)** Cell counts revealed a total of 3084.81±308.44 DβH^+^ (green) neurons in the LC. 2456.909±97.617 cells were found expressing the virus (magenta), of which 1899.9±286.48 were also DβH^+^ (grey), yielding a percentage of 62.07%±10.4% infected noradrenergic neurons **(I). (J)** c*Fos* analysis in triple positive neurons showed a significant reduction in neuronal activity in hM4Di-mCherry injected and CNO treated mice (53.09±16.47, n=11) compared to the control group injected with mCherry virus (1059.8±239.73, n=5) (p=0.002). **(K)** There is a mild negative correlation between the percentage of hM4Di-mCherry infected neurons and the dystonia score when attacks occurred in the presence of CNO (Spearman correlation: −0.532). The more lc neurons were infected with hM4Di-mCherry, the fewer dystonic attacks occurred in the three cage changes performed under CNO administration (Pearson correlation: −0.73). See table S6 for data and p-values. Data are presented as mean±SEM or median±75%/25% quartiles. Statistical significance was evaluated with Mann-Whitney U Rank Sum test, student’s t-test or Two Way Repeated Measure ANOVA, post-hoc Holm-Sidak all pairwise multiple comparison procedure (*p≤0.05, **p≤0.01, ***p≤0.001). DβH = dopamine-β-hydroxylase; LC = Locus coeruleus;

We bred tottering^tg/tg^ with tyrosine hydroxylase (Th)-Cre mice to selectively express a Cre-dependent inhibitory DREADD (AAV9-EF1α-DIO-hM4Di-mCherry) or control virus (AAV9-EF1α-DIO-mCherry) in Th^+^-LC neurons (Figure 6A, B). DREADDs allow a non-invasive manipulation of the neuronal network via administration of an exogenous drug, clozapine-N-oxide (CNO), which can be given either through injection or drinking water (*52*, *53*). We tested both oral administration of CNO (Figure 6) and CNO injection (Figure S7) regarding their effectiveness in preventing stress-induced dystonia. We found that oral CNO consumption is more favorable than intraperitoneal (i.p.) injection (Figure S7), since it was more effective at alleviating stress-induced dystonia in the cage change stress (Figure 6C-F), cage transport (Figure S6A-D) and restraint stress (Figure S6F-I) of hM4Di-mCherry injected, but not mCherry injected tottering^tg/tg^ mice. To eliminate the possibility that chronic DREADD activation through long-term consumption of CNO or sucrose consumption alone decreased dystonia, we administrated 2% sucrose and water alone as controls.

Inhibitory DREADD injected mice displayed a 70% reduction in frequency of stress-induced dystonia with CNO compared to sucrose or water consumption, while control mice showed no changes in dystonia frequency (Figure 6C). Additionally, we observed a delay in onset of stress-induced dystonia when hM4Di-mCherry tottering^tg/tg^ mice were given CNO compared to water and sucrose consumption or compared to mCherry injected mice (Figure 6D). Dystonic episodes of hM4Di-mCherry injected animals still undergoing attacks were considerably shorter (Figure 6E) and milder compared to mCherry control mice (Figure 6F). The more stressful cage transport and restraint tests of hM4Di-mCherry injected mice yielded identical reductions in dystonia frequency, delayed onset and shorter as well as milder dystonia once attacks occurred (Figure S6A-H). To rule out side effects of CNO on locomotion (*54*, *55*), mice underwent an additional open field test (Table S2). It is noteworthy that a relative high dose of 5 mg/kg CNO was needed to reduce dystonia, however CNO is 20x less potent at inhibitory compared to excitatory DREADDs (*55*). Interestingly, the same study reported that α1_A_-ARs and α2_A_-ARs are completely or partially blocked by CNO, respectively. However, our control mCherry injected tottering^tg/tg^ mice did not show any reduction in frequency and dystonia duration or delayed onset of stress-induced paroxysmal dystonia when consuming CNO, thereby eliminating potential contributing off-target effects of CNO in stress-induced dystonia.

A predominant role of the LC in formation of dystonia has been postulated since the 1980’s, however no study investigated its neuronal activity during dystonic episodes. We analyzed our hM4Di-mCherry or mCherry injected tottering^tg/tg^ mice for expression of the immediate early gene marker c*Fos* in the LC as an indirect measurement for neuronal activity, which was shown to increase when mice were exposed to stress (*56–58*) (Figure 6G). The number of LC neurons identified by dopamine-β-hydroxylase (DβH) staining in our injected mice were similar to previously reported values in tottering^tg/tg^ and control tottering^−/−^ mice (*19*), verifying that virus expression and repetitive inhibition with CNO did not cause cell death (Figure 6G-H, Figure S6I). A total of 3085±93 neurons was found in the LC, where 62±3% were infected with DIO-hM4Di-mCherry (Figure 6I) and analyzed for *cFos* expression (Figure 6J). Remarkedly less *cFos* expressing LC neurons were counted in hM4Di-mch injected compared to mCherry injected controls after CNO consumption (Figure 6J), indicating that chemogenetic silencing decreases LC activity, likely resulting in decreased NE release and thus reduction of stress-induced dystonia. Furthermore, we found a negative correlation between the percentage of infected LC neurons and dystonia score and frequency of attacks (Figure 6K), meaning the more LC neurons that were silenced, the fewer or at least milder attacks occurred.

The chemogenetic silencing of LC neurons most likely prevent NE release from presynaptic terminals targeting the cerebellar cortex. Past studies also showed that NE enhances presynaptic GABA release onto PCs by binding to postsynaptic α1-ARs on MLIs (*59*) and MLI electrical stimulations mimic NE-like inhibitory effects on PCs (*60*), suggesting a MLI contribution to dystonia formation. To explore the role of MLIs in stress-induced dystonia, we silenced cerebellar MLIs with DIO-hM4Di-mCherry in tottering^tgtg^/Gad2-Cre mice. Following oral administration of CNO for five days, no alleviation of stress-induced dystonia were observed during the cage change test (Figure S8), strengthening the predominant role of PCs in dystonia (*33*).

Our findings are the first to present an inevitable role of the LC in formation of stress-induced dystonia. Chemogenetic inhibition of the LC alleviates stress-induced dystonia with significant reduction in dystonia severity, suggesting the LC as a new therapeutic target to alleviate dystonia in EA2 patients.

## Discussion

EA2 is an episodic neurological disorder caused by mutations in the Ca _V_2.1 channel (*41*, *61*, *62*), where patients suffer from attacks of severe motor dysfunction elicited through emotional or physical stressors. The ataxic EA2 mouse model tottering^tg/tg^ displays ataxia and episodes of severe paroxysmal dystonia (*10*), which are reliably provoked by stress, the most common trigger in episodic neurological disorders (*8*, *9*). Stress is mediated through release of NE from the LC (*51*), the sole noradrenergic source for the cerebellum, where LC noradrenergic terminals almost exclusively synapse on PCs (*14*). Tottering^tg/tg^ mice are LC - NE hyperinnervated (*19*) and noradrenergic blockade of α1-AR was shown to prevent stress-induced motor attacks (*20*, *21*).

In accordance with the general hypothesis that stress triggers the episodes of motor dysfunction in these mice, we report here that NE release onto PCs is indeed the cause for dystonic events through activation of specifically α1_D_-ARs in EA2 tottering^tg/tg^ mice. Chemogenetic silencing of noradrenergic neurons in the LC decreased neuronal activity and directly correlates to dystonia frequency and severity in tottering^tg/tg^ mice (Figure 6K), thereby verifying that LC activity drives stress-induced dystonia. We show that NE activation of α1_D_-ARs on PCs are the predominant adrenergic subtype driving stress-induced episodic paroxysmal dystonia through long-lasting inhibition of PC SS firing (Figure 2, Figure S4) (*15–18*). This inhibition of PC activity is caused by the bimodulatory effects of NE through direct excitation or indirect inhibition (*36*). Activation of presynaptic β_2_-ARs on MLIs causes excitation, resulting in increased MLI firing and high potentiation of GABAergic transmission at the basket neuron to PC synapse (*32*, *63–66*). However, neither β-AR blockade nor chemogenetic silencing of MLIs prevented stress-induced dystonia (Figure S8) (*20*, *21*). Synaptic transmission in tottering^tg/tg^ mice is mediated by slower N-type Ca^2+^ channels as opposed to the WT P/Q-type channel, which display increased sensitivity to GPCR mediated inhibition and thereby producing a 3-5 fold higher NE-driven synaptic inhibition through GABA_B_-R and α2-AR (*67*). Additionally, NE binding to presynaptic α1-ARs on MLIs increases their firing (*32*, *68*), and simultaneous binding to postsynaptic α1-ARs enhances MLI presynaptic inhibitory transmitter release onto PCs (*59*), supporting the theory that postsynaptic α1-ARs, specifically α1_D_-ARs on PCs, are the main driver of stress-induced dystonia. In agreement with this theory, we show that pharmacological blockade of α1_D_-ARs in the cerebellum by BMY-7378 (*25*) diminished, but not abolished the NE-induced SS depression of PCs, thereby suggesting that NE enhances the GABA-mediated inhibition of PC SS through postsynaptic α1_D_-ARs. In addition to altered NE innervation (*69*), we show that cerebellar α1_D_-ARs mRNA levels are increased in tottering^tg/tg^ (Figure 1F). Moreover, immunohistochemical staining indicates expression mainly on PCs, likely at postsynapses (*21*, *70*) (Figure 1 G). In accordance with our findings, a study in humans also found α1_D_-AR mRNA exclusively in PCs (*30*). Our findings contribute significantly to the understanding of why PCs of tottering^tg/tg^ mice display increased susceptibility to NE-mediated inhibition (Figure 2). Consequently, pharmacological blockade of cerebellar α1_D_-AR through BMY-7378 rescued PC SS firing (Figure 2K-N) and prevented stress-induced dystonia (Figure 1C, Figure 3–4). However, cerebellar *in vivo* pressure injection of BMY did not fully recover SS firing. This may be due to NE activation of α1_A_-, and α1_B_-ARs on GABAergic presynaptic terminals, leading to increased spontaneous inhibition of PC postsynapses (*59*). BMY-7378 can also bind to α2_C_-ARs (*71*). However, we precluded α2-AR involvement in dystonia formation, since Yoh application did not alleviate stress-induced dystonia (Figure 1A), or rescued PC firing (Figure 2O-R) (*20*). More importantly, knockdown of specifically cerebellar α1_D_-ARs using shRNA completely abolished stress-induced dystonia without visible motoric side effects as seen in α1_D_ knock out mice (Figure 4) (*72*).

The mechanism, how blockade of α1-ARs prevents stress-induced attacks of motor dysfunction was recently postulated (*73*, *74*). Snell et al showed that NE binding to α1-ARs results in a reduced open probability of small conductance K^+^ channels (SK) through phosphorylation of SK-associated calmodulin (CaM) through SK2-associated casein kinase 2 (CK2) (*21*). CaM functions as a Ca^2+^ sensor at the intracellular carboxy terminus of the SK channel, consequently CaM phosphorylation lowers its affinity to Ca^2+^, thereby reducing the open probability of SK2 channels (*75*, *76*). SK channels are also required for the intrinsic pacemaking activity in PCs (*33*, *77*) and reduction of their activity through NE inhibition leads to further disruption of the already impaired PC firing in tottering^tg/tg^ mice (Figure S3) (*33*, *78*, *79*), which strongly correlates to dystonia severity (*37*). Interestingly, Snell et al noted that CaM dephosphorylation is required for the recovery of PC firing regularity. This dysfunction in PC firing recovery is hypothesized to be much slower than the rise and fall of NE levels in the cerebellum and thereby dictating the duration of motor attacks (*21*). Strikingly, we found that tottering^tg/tg^ mice displaying dystonia in the presence of BMY-7378 (Figure 1D, E; Figure 3E, I), had significantly milder and shorter motor attacks, suggesting that BMY-7378 may have partially rescued dysfunctional PC firing. Our Ca^2+^ imaging data verified that BMY-7378 regulates the PC firing frequency of ROI dendritic signaling close to those of tottering^−/−^ levels (Figure 5). We confirmed that a rise of intracellular Ca^2+^ is responsible for formation of stress-induced dystonia in tottering^tg/tg^ mice. This was reflected in the rise of intracellular Ca^2+^ recorded and increased integral at the early stages of dystonia formation after NaCl injection (Figure 5G, I), while ROI activity was not altered (Figure 5E). Interestingly, we found that after the dystonic attack is over both integral and ΔF/F return close to pre-dystonia levels (Figure 5E, G, I). We assume that BMY-7378 injection decreases the ROI activity and prevents the release of intracellular Ca^2+^, as it impedes the activation of the Gq-PKC-IP3 pathway stimulated by α1-ARs (*47*), resulting in release of Ca^2^ from intracellular stores through IP_3_R and RyR. Studies showed that RyR1 is expressed in cerebellar PCs, while RyR3 is expressed on granular cells (*45*, *80*, *81*). Interestingly, RyR1 levels are decreased in tottering^tg/tg^ mice (*46*) and RyR blockers effectively alleviated both stress-induced and caffeine induced attacks of paroxysmal dyskinesia predominantly through PCs in tottering^tg/tg^ mice, strongly indicating a shared mechanisms of these triggers to provoke attacks (*11*). Here we verified that the release of intracellular Ca^2+^ increases during the onset of stress-induced dystonic episodes in tottering^tg/tg^ mice, contributing to a novel valuable finding in understanding the NE effects on PC firing.

In accordance with previously published studies, our data strengthen the current understanding of NE and the role α1-AR in stress-induced dystonic episodes. In the present study, we demonstrated that cerebellar α1_D_-ARs are an essential contributore to the formation to stress-induced dystonia in tottering^tg/tg^ mice by increasing PC susceptibility to NE. Purkinje cell firing rate and regularity are thought to encode motor-coordination related information (*82*) and are especially sensitive to changes in currents through P/Q-type channels (*33*, *83*). Since PCs from tottering^tg/tg^ mice display ~40% reduction of Ca^2+^ influx through P/Q-type calcium channels (*7*) and signs of axonal damage (*34*), protecting the already altered firing properties by blocking α1_D_-ARs helps to maintain the firing encoded information and prevent stress-induced paroxysmal dystonia caused by aberrant PC firing (*21*, *33*, *37*).

The use of α1-AR antagonist as therapeutical treatment remains to be determined. Prazosin or silodosin, another FDA-approved α1-AR antagonist, are suggested as therapeutics to treat EA2 patients (*21*). Especially prazosin was shown to cause headache, dizziness, palpitation and tachycardia (*84*, *85*). Development and further exploration of α1_D_-AR specific drugs may not only be beneficial for EA2 patients, but for all patients suffering from episodic neurological disorders where stress is a common trigger (*74*). The α1_D_-AR may provide a more specific therapeutic target in treating episodic neurological disorders.

## Material and Methods

### Animals and genotyping

Genomic *Cacna1a* mutant mice tottering^tg/tg^ (JAX stock #000544; B6.D2-Cacna1a^tg/J^; gift from CI de Zeeuw) were bred with Gad2-Cre mice (JAX stock #010802; Gad2^tm2(cre)Zjh^/J,(*86*) and Th-1-Cre mice (JAX stock #008601; B6.Cg-7630403G23Rik^Tg(Th-cre)1Tmd^/J, (*87*) to obtain tottering^tg/tg^/Gad2-Cre and tottering^tg/tg^/Th-1-Cre mice, respectively. The genomic background of the mice was determined by PCR of genomic tail biopsy. The tottering allele was amplified using the following primer: *tott* forward 5’ TTCTGGGTACCAGATACAGG 3’, *tott* reverse 5’ AAGTGTCGAAGTTGTGCGC 3’. For Cre recombinase identification, the primers were: cre fw 5’ ATTCTCCCACCACCGTCAGTACG 3’, cre rev 5’ AAAATTTGCCTG CATTACCG 3’.

Adult mice (3-8 months) were single housed in the laboratory on a 12 h light/dark cycle with *ad libitum* access to food and water. Experiments were performed during the wake cycle of the animals. The present study was carried out in accordance with the European Communities Council Directive of 2010 (2010/63/EU) for care of laboratory animals and approved by a local ethics committee (Bezirksamt Arnsberg) and the animal care committee of North Rhine-Westphalia, Germany, based at the LANUV (Landesamt für Umweltschutz, Naturschutz und Verbraucherschutz, Nordrhein-Westfalen, D-45659 Recklinghausen, Germany). The study was supervised by the animal welfare commission of the Ruhr-University Bochum. All efforts were made to minimize the number of mice used for this study.

### Drug administration

Stock solutions of adrenergic receptor antagonist prazosin hydrochloride (Sigma Aldrich, P7791), yohimbine hydrochloride (Sigma Aldrich, Y3125) and BMY-7378 dihydrochloride (Tocris, #1006) were prepared in distilled water (d_2_H_2_O) and stored as recommended by the manufacturer. Working solutions were prepared daily using sterile NaCl (vehicle) in doses of 5 mg/kg, 20 mg/kg and 10 mg/kg, respectively. Mice were intraperitoneally (i.p.) injected 30 min prior to the tests, with each mouse receiving each drug at least once. Test days were followed by at least one recovery day. The DREADD ligand CNO (clozapine-N-oxide, HelloBio, #HB6149) was prepared in d_2_H_2_O and diluted in sterile NaCl in working concentrations of 1 mg/kg, 3 mg/kg and 10 mg/kg. Stock and working dilutions were stored at −20°C for 14 days. CNO was administrated either orally or via i.p. injection.

### Stress-induced dystonia

Stress-induced dystonic attacks were triggered by either changing the cage (cage change stress), transport of mice for 10 min (cage transport stress) or restraining the animals for 10 min (short restraint) and releasing them to a novel cage, according to previous methods (*39*, *40*) which were shown to elicit attacks in tottering^tg/tg^ mice (*9*, *20*, *21*). Mice were observed for the presence or absence of dystonic attacks for 40 min and onset and duration of episodic dystonia were noted. Since dystonia follows a characteristic progression (*9*, *10*), attacks can be classified and compared, thus severity of dystonia was scored as described below. After expiration of this time or when dystonia was over, mice were returned to their home cages and given at least one day to recover.

### Dystonia characterization

To quantify presence of dystonia, the overall motor behavior of animals was observed. Dystonia severity was scored every 5 min until end of attack following a modified scale as previously published (*26*). Briefly, 0 = normal motor behavior; 1 = slightly slowed and/or abnormal movements; 2 = mild impairments, limited ambulation; 3 = moderate impairments, limited ambulation when disturbed; 4 = severe motor impairments, almost no ambulation, sustained abnormal postures; 5 = prolonged immobility in abnormal postures, no ambulation. Dystonic episodes were scored severe, when a median score ≤ 3.5 or higher was reached, otherwise it was scored mild (score ≥ 3). The median of dystonia severity was first calculated per mouse/time, then for all mice within a group. Corresponding graphs display the median and the 75% and 25% quartiles per time.

### Behavior experiments

To evaluate ataxia, motor-coordination and balance skills, 6 month old tottering^tg/tg^, and control tottering^−/−^ mice underwent a beam walk test, gait analysis, hangwire test, rotarod and vertical pole test according to previous methods (*39*, *40*). Mice received either a saline or 5 or 10 mg/kg prazosin hydrochloride (in NaCl) i.p. injection 30 min prior to testing. Mice were given one day to recover between tests and drugs.

#### Beam walk

To access balance skills and motor coordination, mice were trained to cross a 70 cm long, 1 cm wide beam in 60 cm height. The start platform was illluminated and mice trained to cross the beam to reach the dark shelter (20 cm^3^) on the other site. During training, mice were shown the shelter, then placed on the beam with increasing distances until the beam was crossed completely. Mice were given a rest day and tested the following day. Mice were placed on the illuminated platform and the latency to step on the beam (Idle), time to cross the beam and slips of paws was recorded. A maximum period of 120 sec was given to perform the test. Falls were noted and scored as 120 sec. Each individual was tested 3 × and the average was calculated.

#### Gait analysis

To evaluate ataxia, mice underwent the gait analysis. Individuals were placed at the end of a 10×70×10cm (width × length × height) plastic chamber and trained to walk to the other end, where a hole connects the box with their home cage. To obtain footprints, forepaws were painted red and hindpaws painted blue with non-toxic, water-soluble childrens paint (Pelikan) and the box laid out with white paper. The length and width between steps were measured by hand. Each mouse underwent one trial per condition.

#### Hangwire

As an indicator for grip and muscle strength, the hangwire test was performed. Mice were placed on a wire screen (12mm × 12mm grid) and hung upside-down above a 15.5×29×26.3cm (width × length × height) box for a maximum of 60 sec. The latency to fall was recorded. Mice that did not fall within the trial period were removed and given a 60 sec score. Each mouse was tested twice and the average taken.

#### Rotarod

To screen for motor coordination and balance skills, mice underwent the rotarod test (Columbus Instruments). Therefore, mice were placed on a rod (3 cm) rotating at 4 rpm for 1 min (acclimatization phase), followed by a speed increase of 0.1 rpm/s to a max of 40 rpm. The trial was over when all mice fell, latency to fall and speed where recorded. Mice that fell during the acclimatization phase were placed back on the rod 3x, after that they received a score of 4 rpm and 0 sec time.

#### Vertical pole test

Motor coordination and balance skills were accessed using the vertical pole test. A 52 cm long metal pole wrapped in tape for grip was secured to a platform. The pole was placed in an empty cage and the platform and floor covered in towels. Mice were positioned on top of the pole face upwards. Their latency to turn around and time to climb down the pole were recorded. A max of 120 sec was given to complete the test. Mice falling were given the maximum time. Individuals were tested 3 times and the average was taken.

#### Open Field

Mice were placed in the center of a 50 cm × 50 cm opaque plastic box and allowed to freely explore it for 5 min. Their exploring path was tracked using the EthoVisionXT 8.5 video tracking software (Noldus). The total distance moved (cm) and time spent in the center or border region of the arena were analyzed. The apparatus was cleaned between subjects with 70% ethanol. Each mouse underwent 1 trial. Statistical significance was analysed using SigmaPlot software (Systat Software Inc.).

### QPCR

#### Quantification of adrenoreceptor expression

tottering^tg/tg^ (n = 10) and tottering^−/−^ (n = 10) 3 – 4 months old mice were deeply anaesthetized and quickly decapitated. Cerebella were immediately removed and frozen in liquid nitrogen. RNA isolation was conducted using the ReliaPrep^TM^ RNA Tissue Miniprep System (Promega) and concentration (ratio A260/A280) and quality (ratio A260/A230) of RNA were estimated using a NanoDrop 2000c Spectrophotometer (Thermo Fisher Scientific). cDNA synthesis was performed using the RevertAid First Strand cDNA synthesis kit (Thermo Fisher Scientific). For qPCR, probes were amplified in triplicates using the SYBR green based GoTaq® qPCR Mastermix (Promega) in a Rotor-Gene Q real-time PCR cycler (Qiagen) in a twostep program (40 cycles: 15 sec 95°C, 90 sec 60°C). Obtained ΔCT values were analyzed with the corresponding Rotor-Gene Q Series Software. Normalization was performed through geometric averaging of multiple reference genes (*88*), previously selected for cerebellar expression stability (*89*) and confirmed with the NormFinder software (*90*). Primer specificity was verified via melting curve analysis and gel electrophoresis. All primers were obtained from Invitrogen (Germany) and are listed in table S5.

#### Analysis of sh-RNA induced α1_D_-KO

Both tottering^tg/tg^ injected with either sh-[ADRA1D] (n=9) or sh-[scramble] (n=5) were sacrificed as mentioned above. One hemisphere was used for qPCR analysis, the other half was used for WB analysis.

### Histology

Mice were deeply anesthetized with ketamine and xylazine (100 mg/kg and 10 mg/kg, respectively) and perfused transcardially with 1x PBS followed by ice-cold 4% PFA (paraformaldehyde, Sigma-Aldrich) in PBS (pH 7.4). Brains were dissected and post-fixed for 1 h in 4% PFA, then cryoprotected in 30 % sucrose in PBS overnight before slicing using a cryostat (Leica CM3050 S).

#### cFos and dopamine-β-hydroxylase staining

For cFos and *dopamine-β-hydroxylase* staining after chemogenetic inhibition, 25 µm thick coronal slices of the locus coeruleus were prepared. Sections were caught and rinsed in 1x PBS (3 × 15 min) prior to incubation in blocking solution (5% NDS (Merck) in 0.3% PBS-Triton-X-100) for 1 h at RT, followed by incubation of primary antibodies rabbit-α-*dopamine-β-hydroxylase* (1:700, #AB1585, Sigma Aldrich) and mouse-α-cFos (1:500, C-10, sc-271243, Santa Cruz) in blocking solution overnight at 4°C. Slices were washed with 1x PBS before 2 h incubation of corresponding secondary antibody solutions containing either donkey-α-rabbit DyLight 488 (1:700, #SA5-10038, Thermo Fischer) and/or donkey-α-mouse DyLight 650 (1:1000, #SA5-10169, Thermo Fisher) in blocking solution. Slices were rinsed again, then transferred to Superfrost Plus microscope slides and mounted with Mowiol containing Dabco.

#### Adrenoreceptor staining

35 µm sagital slices were co-stained with mouse-α-calbindin (1:300, Sigma, C9848). Sections for staining with rabbit-α-α1_D_ (1:200, Alomone labs, AAR-019), rabbit-α-α2_A_ (1:100, Alomone labs, AAR-020), and rabbit-α-α2_B_ (1:100, Alomone labs, AAR-021), were caught and washed in 1 × PBS, followed by blocking in 10% NDS in 0.3% PBST and incubation in primary antibodies at 4°C overnight. Sections for rabbit-α-α1_A_ (1:250, Invitrogen^TM^, PA1-047), rabbit-α-α1_B_ (1:100, Alomone labs, AAR-018), and rabbit-α-α2_C_ (1:100, Alomone labs, AAR-022) were caught and washed in 1 × TBS, blocked in 10% NDS in 0.25% TBS-T before overnight incubation of primary antibodies in blocking solution at RT. The next day, slices were washed and incubated 3 h at RT in secondary antibodies donkey-α-rabbit DyLight 650 (1:500, SA5-10169, Thermo Fisher) and donkey-α-mouse Alexa 568 (1:500, Invitrogen^TM^, A10037) in blocking solution, respectively, before mounting.

Fluorescent Nissl staining (NeuroTrace^TM^ 435/455, Invitrogen^TM^ N21479) was conducted before blocking.

#### Calbindin staining

For Purkinje cell staining of shRNA and GCaMP8m injected cerebella, chicken-α-*calbindin (1:500,* antibodies.com, *A85359)* was conducted. 25 µm thick sections were washed 3x in 1x PBS, followed by blocking in 10% NDS in 0.5% PBST and incubation in primary antibody at 4°C overnight. The next day, slices were rinsed in 1x PBS, then incubated in donkey-α-rabbit DyLight 650 (1:500, SA5-10169, Thermo Fisher) and donkey-α-chicken Cy3 (1:500, AP194C, Merck) in blocking solution for 2 h at RT.

### Imaging

All images were obtained using an inverted Leica TCS SP5 confocal laser scanning microscope (Leica DMI6000 B, Wetzlar Germany) interfaced to a computer running the Leica Application Suite Advanced Fluorescence software (LAS AF 2.6). Sequential scans were performed according to the required wavelengths using 20x or 40x objectives. Sequential z-stacks of 10 to 15 images were made for each section and crosstalk of fluorophores was eliminated automatically by the LAS AF 2.6 software. Acquired images were analyzed using ImageJ (NIH).

### Electrophysiology

#### Recordings

Electrophysiological recordings were conducted as previously described (*91*). Briefly, mice were deeply anesthetized with 2% isoflurane via a precision vaporizer (E-Z Anathesia, Euthanes Corp, Palmer, PA, USA) and placed in a stereotaxic frame (SR-6M, Narishige, Tokyo, Japan). Mice received 2 mg/kg rimadyl s.c. and 2% lidocaine hydrochloride locally before surgery. A craniotomy (2×2 mm) was performed above the recording site, approximately – 6.5 mm from bregma using a dental drill. The dura mater was removed and extracellular recordings of PCs were performed using a multielectrode system (Eckhorn system, Thomas Recording, Giessen, Germany) with 5 electrodes (impedance, 2 – 3 MΩ, at 1 kHz, Thomas Recording) simultaneously. Signals were amplified and filtered (band-pass, 0.1 – 8 kHz) with a multichannel signal conditioner (CyerAmp380, Axon Instruments, Union City, CA, USA) and sampled with 32 kHz via a A/D converter (NI PCI-6259 multifunction data acquisition board, National Instruments, Austin, TX, USA), controlled via a custom-made software using MatLab (MathWorks, Natick, MA, USA) as previously presented (*91*). For microinjection of adrenoreceptor modulators diluted in ACSF, quartz glass microinjection pipettes (outer diameter: 115 µm, inner diameter: 85 µm; Thomas Recording, Giessen, Germany) were positioned next to the recorded cell. Via pressure injection, norepinephrine (20 mM, Sigma Aldrich, A7257) was applied after 150 sec of reference recording and its effects recorded for 150 sec before application of the antagonists was conducted (1 mM yohimbine hydrochloride, α2-AR antagonist, Sigma Aldrich, Y3125; 1 mM prazosin hydrochloride, α1-AR antagonist, Sigma Aldrich, P7791; 1 mM BMY-7378 dihydrochloride, α1D-AR antagonist, Tocris, #1006). Recordings were conducted for 6 min and saved for offline analysis.

#### Data analysis

Single unit action potentials were detected with custom-made software implemented in Matlab (*91*). Purkinje cells were identified by their unique firing properties. Simple spikes occur spontaneously at a frequency of 20 – 150 Hz, while CS were observed with < 1 spikes/sec, characterized by a strong depolarization spike and multiple wavelets. Cells were analyzed, when a CS was followed by a SS pause of ~ 25 ms, proving spikes were generated by one single PC. The mean firing rate of both SS and CS within the 150 sec recording intervals was compared. In addition coefficients of variation (CV) of interspike intervals (ISIs) *[CV = stdev (ISI)/mean(ISI)]* and the coefficients of variation for adjacent intervals CV2 of *ISIs [CV2= 2 | ISIn+1–ISIn | /(ISIn+ISIn+1)]* were calculated. An average of CV2 over n estimates the intrinsic variability of a spike train, nearly independent of slow variations in average rate. Error bars denote SEM.

### Chronic cerebellar drug infusion

Chronic perfusion of the cerebelli of tottering^tg/tg^ mice was performed using the *ALZET®* osmotic pumps (model 1007D, Durect), connected to the ‘Brain Infusion Kit 3’ (#0004760, Durect). Pumps were prepared sterilely according to the manual. Briefly, the vinyl catheter tubing was cut to ~3 cm length, connected to the infuser and 2 spacer discs were glued to the infusion cannula to allow tissue penetration of ~ 1.5 mm depth. Pumps, catheter and the infuser were filled with either 1 mM BMY in ACSF or ACSF alone before connecting all parts. 0.01% methylene blue was added to solutions to allow for postmortem identification of the perfusion site. For osmotic equilibration, assembled pumps were incubated overnight at 37°C in 0.9% NaCl. The next day, mice were prepared for surgery as already described and a craniotomy was drilled at AP: −6.5 mm from bregma. The osmotic pump was inserted into a lateral subcutaneous pocket of the animal, while the infusion canula was inserted into the craniotomy and fixed to the skull using the light-cured Gradia® DIRECT Flo composite (GC corporation). Mice were placed in their home cages and monitored closely for recovery. Carprofen (100µl/100ml) was orally administrated for 5 days. The cage change stress test paradigm was performed 5-, 7-, 10- and 14 days post implantation to monitor frequency and severity of stress-induced dyskinesia.

### Intracerebellar virus injections

Mice received subcutaneous injection of carprofen (2 mg/kg) for analgesia 30 min prior to surgery, then deeply anesthetized with 1.5%–2.0% isoflurane and placed into a stereotactic frame (Narishige, Japan). OmniVision® gel was applied to the eyes to prevent dehydration. 2% lidocaine hydrochloride was subcutaneously injected at the scalp for local anesthetic. The skin was disinfected and opened with a sagittal incision along the midline. Craniotomies were performed above the LC or CB vermis according to the methods of Bohne et al (*92*). Briefly, the virus was sucked into a customized glass pipette attached to a 5 ml syringe. Via pressure injection, the virus was dispensed in 50-100 µm steps with a time interval. After injection, the skin was sutured (Surgicryl Monofilament, Belgium) and mice were removed to their home cages. Analgesia was applied for 5 subsequent days or longer when required and animals were given 14 - 21 days to recover from surgery.

### shRNA mediated KO of α1_D_-ARs

shRNA sequences against *Mus musculus* Adra1d were obtained from VectorBuilder in AAV2-eGFP-U6-shRNA packed adeno-associated viruses. AAVs used included the following: AAV2-CMV-eGFP-U6-mAdra1d[shRNA#1] at a titer of 3.62×10^13^ GC/ml; AAV2-CMV-eGFP-U6-mAdra1d[shRNA#2] at a titer of 4.04×10^13^ GC/ml; AAV2-CMV-eGFP-U6-mAdra1d[shRNA#3] at a titer of 4.92×10^13^ GC/ml and AAV2-CMV-eGFP-U6-Scramble at a titer of 1.20×10^13^ GC/ml. 3 – 4 months old tottering^tg/tg^ and tottering^−/−^ mice of both sex were injected at the following injection sites: AP: −6 mm, ML: 0 mm, DV: −1.2 – −0.9 mm; AP: −6 mm, ML: ±1. 8 mm, DV: −2 – −1.7 mm; AP: −6.25 mm, ML: 0 mm, DV: −1.2 − −0.9 mm; AP: −6.25 mm, ML: ± 1 mm, DV: −1.7 − −1.4 mm; AP: −6.4 mm, ML: 0 mm, DV: −1.2 – −0.9 mm; AP: −6.4 mm, ML: ± 2.7 mm, DV: −2 − −1.7 mm; AP: −6.6 mm, ML: 0 mm, DV: −1.2 – −0.9 mm; AP: −6.6 mm, ML: ± 1.5 mm, DV: −1.7 − −1.4 mm; AP: −6.96 mm, ML: 0 mm, DV: −1.2 – −0.9 mm; AP: −6.96 mm, ML: ± 1 mm, DV: −1.7 − −1.4 mm. Starting 3 weeks post injection, animals underwent a weekly cage change for 3 weeks. After a total of 6 weeks of expression, mice were killed through cervical dislocation. Their cerebellar were removed and bisected. Hemispheres were individually used for qPCR or WB analysis. One shRNA injected mouse was perfused for verification of virus expression. 100 µm thick vermal sections were taken of one scramble injected moue using a vibratome (Leica) prior to analysis.

### Western Blots

Cerebellar hemispheres were weighted then lysed and homogenized in TDL buffer (1M HEPES buffer (pH 7.4), 5M NaCl, 0.5M EDTA (pH 8), 10% NP40, 10% natrium deoxycholate, 10% SDS, 25x protease inhibitor). Lysates were incubated on ice for 30 min, then mixed and centrifuged at 12.000 – 14.000*g for 10 min at 4°C. Protein concentration was determined through photometric OD_600_ using a Bradford protein assay 5 min after lysis. Equal amounts of Laemmli sample buffer were added to the supernatants and cooked for 5 min at 99°C. Samples were aliquoted and stored at −20°C. Samples were loaded onto 12% T Tris-HCl separation gels covered by 4% T collecting gels and run for 55 – 77 min, 20 mA. Protein transfer onto a PVDF membrane was conducted for 1.5 h at 200mA in running buffer.

#### Antibody incubation

PVDF membranes were incubated in blocking buffer (3% milk powder, 2% BSA in 0.1% TBS-T) for 1 h, at RT on a shaker prior to primary antibody incubation (see table S4) over night at 4°C. Membranes were washed 3 × 5 min in 0.1% TBS-T and incubated in secondary antibodies (see table S4) for 1.5 h at RT. Membranes were washed again prior to detection.

#### Detection

Detection was performed using the ECL^TM^ Select Western Blotting Detection Reagent (Amersham^TM^). Equal amounts of the two detection reagents were mixed and added to the PVDF membranes. Incubation was performed for 5 min at RT before transferring the PVDF membranes to a clear plastic cover and immediately placed into a Hypercassette^TM^ (Amersham Pharmacia Biotech, Inc.). Under red-light conditions, a medical X-Ray film (Fujifilm Super RX-N) was placed on the membranes and exposed for 30 s to 2 min depending on the intensity of the bands. X-Ray films were developed in developer solution (AGFA Dentus D-1000, Kulzer) for 2 min, washed in water, and fixed in fixer solution (AGFA Dentus F-1000, Kulzer) for 2 min. After that, the films were rinsed in water and dried hanging in a ventilated cabinet at RT. X-Ray films were scanned and band intensity analysed using ImageJ (Fiji).

#### Stripping

Protein loaded PVDF membranes were used two times. Therefore, the old primary antibody was removed by initially washing in 0.1% TBS-T for 15, followed by incubation in ROTI® Free Stripping buffer 2.0 (Carl Roth) for 30 min at RT. Membranes were washed twice in 0.1% TBS-T for 20 min, then blocking was repeated.

### *In vivo* Ca^2+^ imaging of Purkinje cells

#### Implantation

Mice were injected with AAV-DJ/2-mCaMKIIα-jGCaMP8m-WPRE-bGHp(A) (#p630, VVF, Zurich) in the cerebellar vermis at AP: −6.25 mm, ML: ± 0 mm, DV: −1.2 & 0.9 mm as already described and given 14 – 21 days for virus expression and recovery. Then, mice were implanted with the ProView^TM^ Prism Probe (1.0 mm diameter, ~4.3 mm length, #1050-004606, Inscopix, USA). Briefly, mice were anaesthetized and fixed to the stereotaxic frame. The skull was primed using OptiBond^TM^ Universal adhesive (#36519, Kerr, Germany) and 1/3 of a screw (2 mm length) was drilled into the skull above the motor cortex for better grip of the implant. A craniotomy (~1.8 mm diameter) was drilled above the injection site using a trephine (FST, #18004-18), the exposed cerebellum was moisturized with ACSF and slowly cut ~200 µm deeper than the virus was injected using a micro knife (FST, #10315-12) either from anterior to posterior for ML imaging experiments, or from medial to lateral for AP imaging experiments. At the cutting edge, the prism was slowly lowered to the desired depth (~200µm deeper than virus was injected) and attached to the skull using light-cured Gradia® DIRECT Flo composite (GC corporation). Mice recovered for 2 weeks before weekly assessment of virus expression and Ca^2+^ signaling before attachment of baseplates (#1050-004638, Inscopix, USA) above the lens using Gradia® DIRECT Flo. Attachment of baseplates was conducted under visual control where the miniscope was slowly lowered towards the lens using the micromanipulator to allow for optimal positioning and distance of the miniscope to the lens for maximal Ca^2+^ signals. Afterwards, the protective cap was placed on the baseplate to protect the lens from scratches, dust or animal bedding that could potentially contaminant the lens. Animals were given at least 2 days to recover from anesthesia before habituated to handling and mounting the miniscope.

#### Imaging

Mice were habituated to handling and mounting of the nVista 2.0 Miniscope Sytem for 2 days. On recording days, mice were injected i.p. with saline or 5 mg/kg BMY-7374 30 min prior to recordings to explore alterations in Ca^2+^ signals of PCs before, during and after stress-induced dystonia. The miniscope was secured to the baseplate and the cage change stress paradigm was conducted. Recordings of calcium transients were acquired using the nVista Inscopix Data Acquisition Software (IDAS, Inscopix, USA) at 30 Hz, LED power 1 – 1.3, Gain 3 – 4). Videos were acquired for 1 min per video in 5 min intervals for a total of 40 min or until end of the dystonic attack. The behavior of mice during the calcium recording acquisition was simultaneously recorded using a USB video camera connected to the laptop to allow for synchronization of behavior and Ca^2+^ signals of mice. Sessions were individually saved and stored on an external hard drive for analysis. Each mouse was recorded several times. Only one recording was performed per day/mouse with 2 day intervals between recordings.

#### Preprocessing

Analysis of Ca^2+^ recordings was performed using the Inscopix Data Processing Software (IDPS, Inscopix, USA). Videos were cropped, temporally down sampled and motion corrected, before ΔF/F was acquired to normalize each pixel value in the recording to a baseline level. Since Ca^2+^ signals varied between recorded mice, individual regions of interest (ROIs), putative PC dendrites were manually encircled and analyzed for their signals. Signals traces were deconvolved and exported for offline analysis.

#### Data analysis

Exported traces were further analyzed and plotted using Excel. Spikes per second (frequency), integral of Ca^2+^ peaks and Ca^2+^ peak height were analyzed. Fluorescence signals indicating calcium transients of individual ROIs were compared within the recorded videos of one session per mouse.

### Virus production

Adeno-associated viruses were either purchased or produced as previously described (*92*). Briefly, low passage 293T cells were co-transfected with pAAV-EF1α-DIO-mcherry or pAAV-EF1α-DIO-hM4Di_(Gi/o)_-mcherry (original plasmid pAAV-hSyn-hM4D(Gi)-mCherry was a gift from Bryan Roth (Addgene plasmid #50475; http://n2t.net/addgene:50475; RRID:Addgene_50475)), pAAV-RC9, and pHelper using the polyethylenimine (PEI) based protocol. Three days after transfection, cells were harvested, pelleted (3,700 g, 20 min, 4°C), resuspended in 10 ml lysis buffer (150 mN NaCl, 50 mM Tris-HCl, pH 8.5) and lysed via seven freeze (15 min)/thaw (10 min) cycles in a dry ice/ethanol and 37°C water bath. The cell suspension was treated with DNase I (Roche) for 30 min at 37°C to degrade free DNA. The cell debris was spun down at 3,700 g for 20 min at 4°C. The supernatant was collected in a syringe and filtered into a 15 ml falcon tube through a 0.2 µm filter to obtain the crude lysate. Then the supernatant was resuspended in a polyethylene glycol (PEG) solution overnight at 4°C and pelleted at 3,700 g for 20 min at 4°C. The pellet was resuspended in PBS, 0.001% pluronic and aliquots were stored at −80°C until further use.

### *In vivo* chemogenetic manipulation

For chemogenetic silencing of the LC-NE pathway tottering^tg/tg^/Th-1-Cre mice were bilaterally injected in the LC with either AAV9-EF1α-DIO-mCherry (n = 5) or AAV9-EF1α-DIO-hM4Di_Gi/o_-mCherry (n = 11 for oral administration of CNO, n = 10 for i.p. injection of CNO) at: AP: −5.8 mm, ML: ± 0.8 mm, DV: −3.25 – 2.7 mm as already described. tottering^tg/tg^/Gad2-Cre mice were injected with AAV9-EF1α-DIO-mCherry (n = 4) at AP: −6.25 mm, ML: ± 0 mm, DV: −1.5 – 0.7 mm. For chemogenetic silencing of MLIs, tottering^tg/tg^/Gad2-Cre mice (n = 4) were injected with AAV9-EF1α-DIO-hM4Di_Gi/o_-mCherry at: AP: −6.3 mm, ML: ±0 mm, DV: −1.7 − −0.9 mm. Animals recovered for 2 weeks to allow for sufficient virus expression.

For direct, short-term silencing of the LC NE projections, mice were i.p. injected with 10 mg/kg CNO or saline 30 min prior to the stress-induced dystonia tests to allow for sufficient clozapine levels in ACSF and penetration of brain tissue (*55*, *93*).

For chronic silencing of the LC or MLIs, 5 mg/kg CNO in 2% sucrose (to mask bitterness) was orally administrated *ad libitum* for at least 5 days prior to testing as previously described (*52*, *94*). Bottles were weighted daily to control sufficient intake of CNO. Solutions were replenished as needed or every third day. Mice underwent the stress-induced dystonia tests after consumption of 2% sucrose, 5 mg/kg CNO or water.

One test per day/mouse was performed and stress tests were conducted in random order, though each mouse underwent each test under each condition three times. To evaluate the level of LC silencing and virus expression, animals underwent a last cage change stress test and were sacrificed 90 min later and stained for cFos.

### cFos counts

For cFos counts in the LC z-stacks were acquired and individually screened for AAV9-EF1α-DIO-hM4Di_Gio_-mCherry/AAV9-EF1α-DIO-mCherry expressing, DβH positive or cFos positive neurons and counted manually using ImageJ. Channels were merged in one z-stack to allow identification of triple positive cells used for analysis.

### Statistical Analysis

Test procedure, statistical significances and number of animals (*n*) for each experiment can be found in table S6 - S7. Statistical analysis was conducted with SigmaPlot (Systat Software), the level of significance was set to p < 0.05. Error bars display mean±SEM, if not stated otherwise Statistical significance is reported as n.s (not significant); * p<0.05; ** p<0.01; *** p≤0.001.

## Supporting information

ADRA Supplementary Material BioRxiv.pdf

## Acknowledgements

We would like to thank Dr. Norbert Hogrefe, Stephanie Krämer, Margareta Möllmann, Petra Knipschild, Winfried Junke, Gina Hillgruber, Manuela Schmidt and Katja Schmidtke for their excellent technical assistance.

## Author contribution

Conceptualization: PB, MDM

Methodology: PB, MOR, XZ, MDM

Investigation: PB, MOR, LR, MJ

Supervision: MDM

Writing—original draft: PB

Writing—review & editing: PB, XZ, MDM

## Data and Materials availability

Data needed to evaluate the conclusions in the paper are present in the paper and/or the Supplementary Materials. For additional information, please contact the corresponding author at

## List of Material contained in the Supplementary Material

- Figures S1-S8
- Tables S1-S7
- Movie S1-S5

