## Supplementary material for "Cerebellar α1_D_-adrenergic receptors mediate stress-induced dystonia in tottering^tg/tg^ mice": ADRA Supplementary Material BioRxiv.pdf

**This PDF file includes:**

Figs. S1 to S8
Tables S1 to S7
Movies S1 to S5

**Other Supplementary Materials for this manuscript include the following:**

Movies S1 to S4

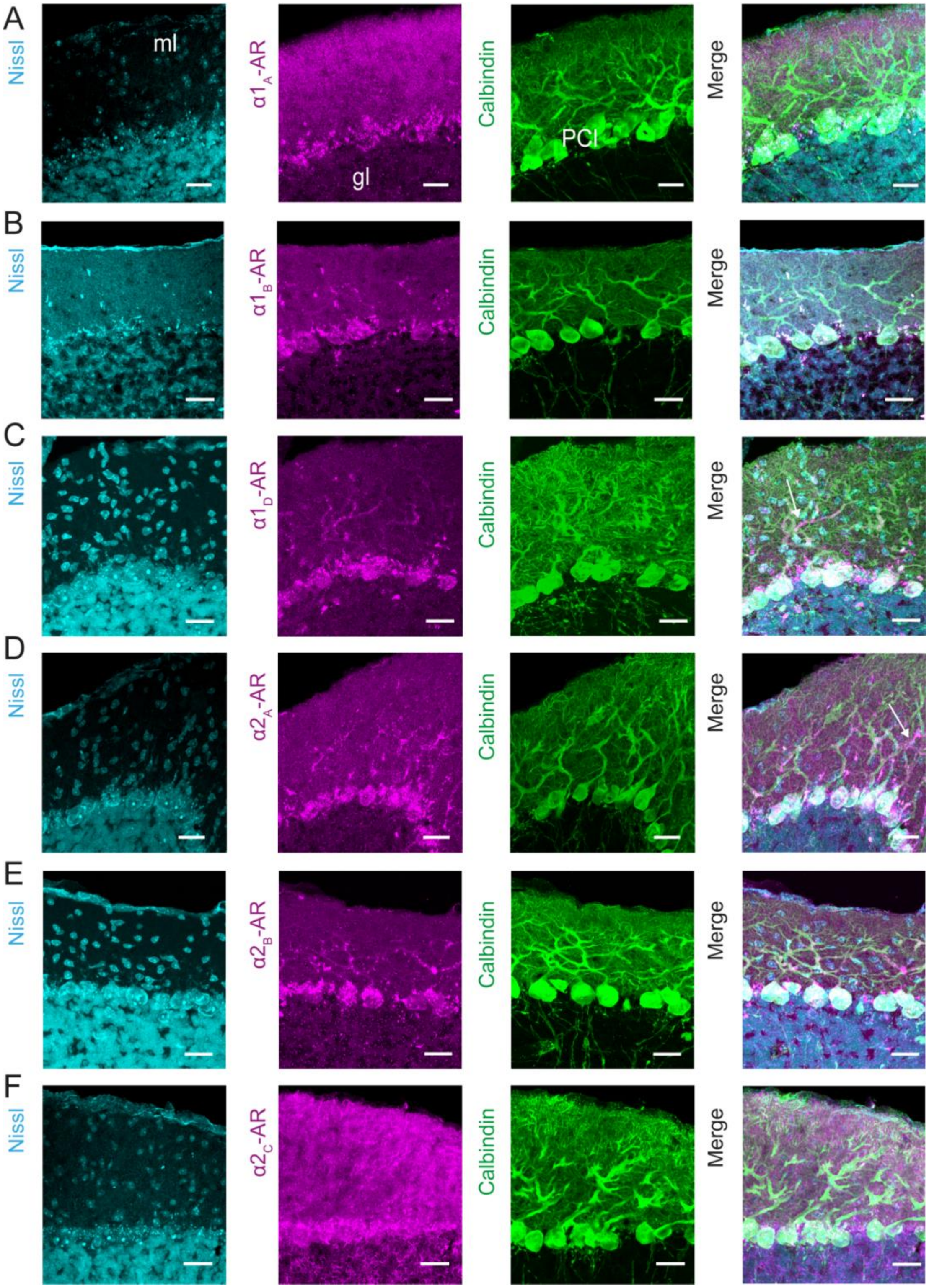


**Fig. S1. Adrenergic receptor localization in the cerebellar cortex of tottering$^{tg/tg}$ mice.**

**(A)** IHC staining against α$1_A$-AR (magenta) shows a punctual expression in the molecular layer and PC soma in the cerebellum of tottering$^{tg/tg}$ mice. Both the α$1_B$- **(B)** and α$1_D$-AR **(C)** are predominantly located on PC soma. **(D)** IHC staining against the $2_A$-AR shows strong labeling of PC soma and dendrites (magenta), as indicated by overlay with calbindin staining (green). **(E)** Overlay of calbinidin and $\alpha 2_B$-AR staining reveals that the $\alpha 2_B$-AR is located on cerebellar PCs. **(F)** IHC staining against the $\alpha 2_C$-AR shows expression in the ml, PCl and gl. Scale bars 25 µm. AR = adrenergic receptor, gl = granular layer, ml = molecular layer, PCl = Purkinje cell layer

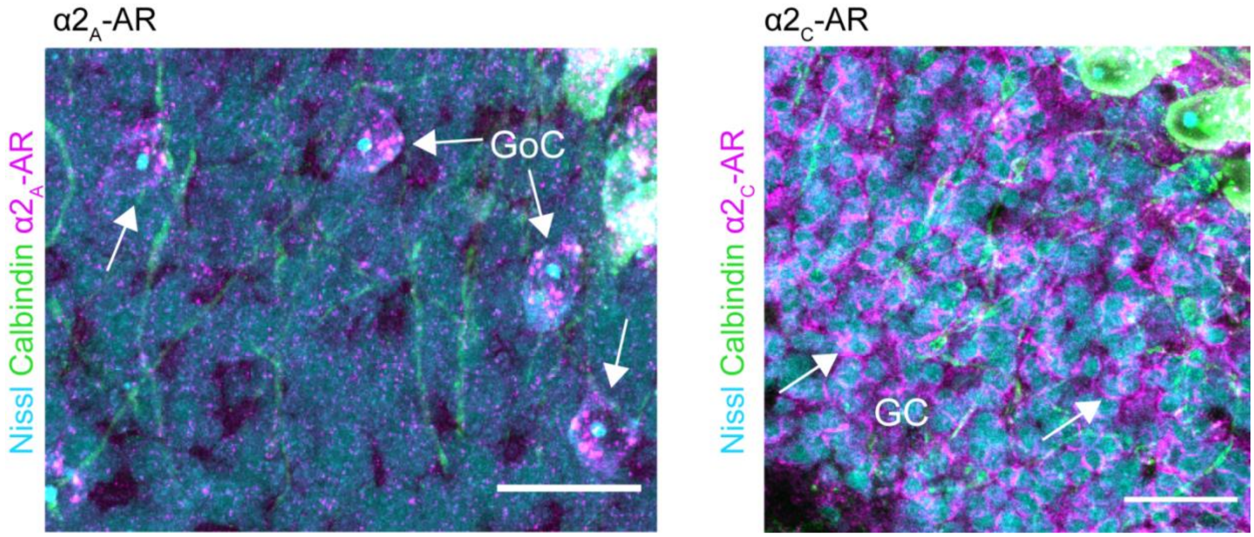


**Fig. S2. $\alpha 2_A$ and $\alpha 2_C$ adrenergic receptors are located on cerebellar Golgi cells (GoC) and Granule cells (GC).**

Antibody staining against the adrenergic $\alpha 2_A$ receptor subtype indicates localization on interneurons in the granular layer (gl), potentially Golgi cells (GoC). Staining against the $\alpha 2_C$-AR shows membrane-like staining around cerebellar granule cells (GC), potentially showing presynapses. Scale bars 25 µm. AR = adrenergic receptor

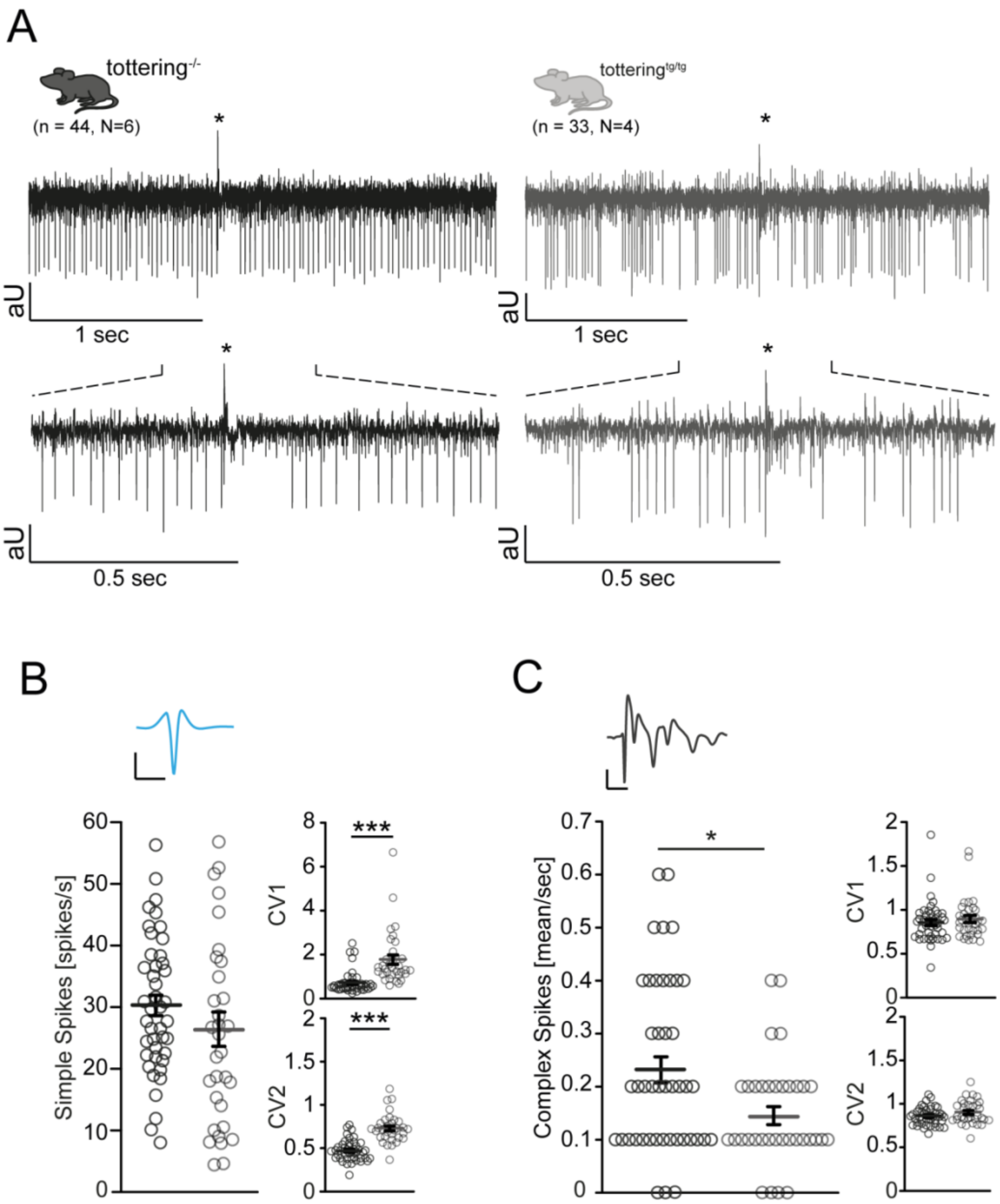


**Fig. S3. Purkinje cell firing is irregular in tottering$^{tg/tg}$ compared to control tottering-/- mice.**

**(A)** Example traces from extracellular recorded PCs in tottering-/- and totteringt$^{g/tg}$, mice displaying the irregularity in PC firing of totteringt$^{g/tg}$, indicating complex spikes (*). **(B)** Mean simple spike firing is not altered in totteringt$^{g/tg}$, mice, but are highly irregular and variable. **(C)** The mean number of complex spikes is decreased in totteringt$^{g/tg}$, compared to control mice, while there is no difference in regularity and variability indicated by CV1 and CV2 values. Data are presented as mean±SEM. Statistical significance was evaluated by students t-test. ($^{*}p \leq 0.05$, $^{**}p \leq 0.01$, $^{***}p \leq 0.001$).

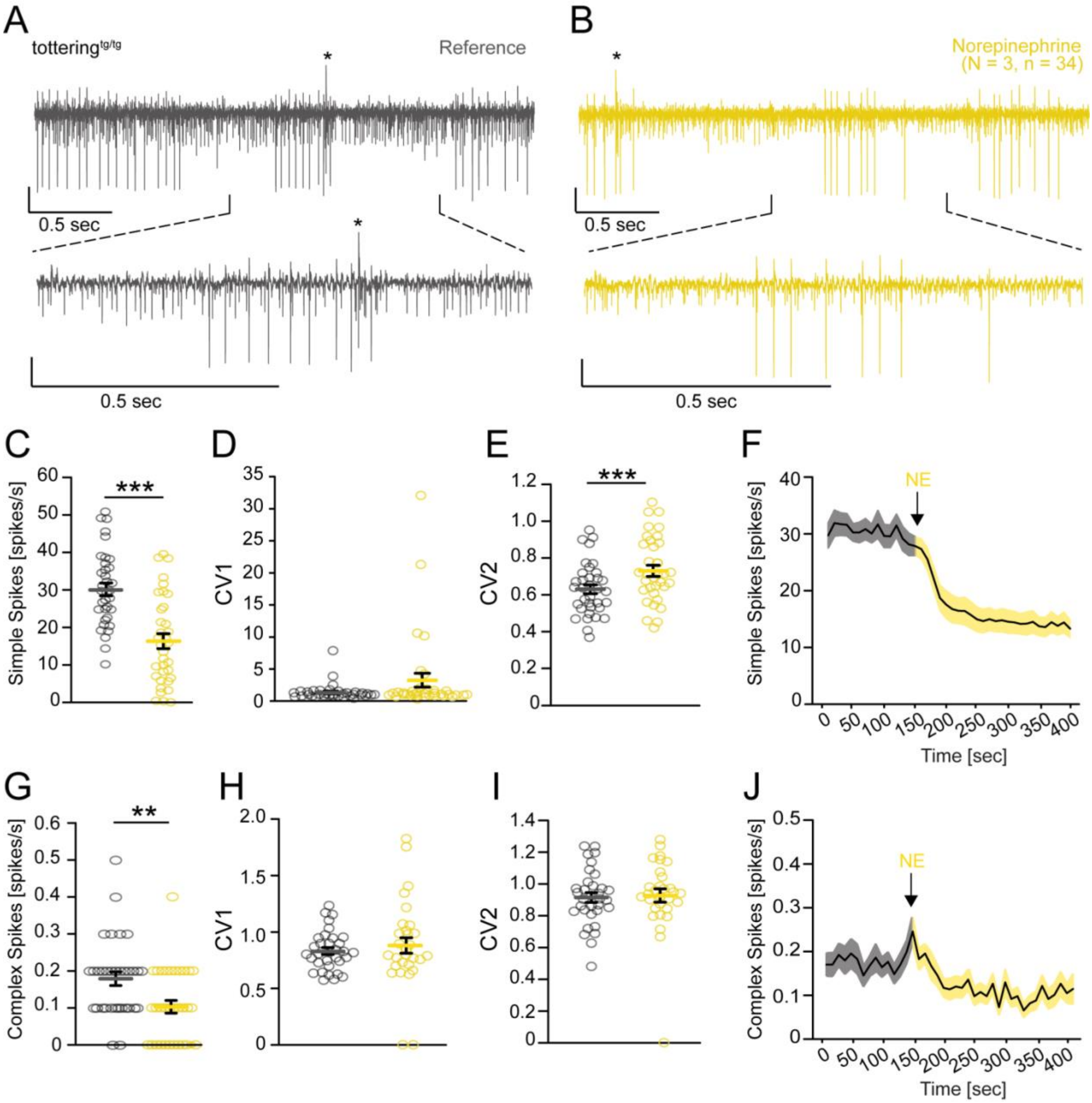


**Fig. S4. Extracellular *in vivo* recordings verify inhibitory effects of NE on tottering$^{tg/tg}$ Purkinje cells**.

Example traces of an extracellular recorded PC of a tottering$^{tg/tg}$ mouse before **(A)** and after exogenously applied NE **(B)**. **(C)** Simple spikes are significantly decreased after NE application. **(D)** CV1 shows a trend to be increased, while CV2 is significantly increased **(E)**. **(F)** Course of the mean simple spike firing of all PCs recorded shows that the firing frequency does not recover after NE application during the recording period of 400 sec. **(G)** Complex spike firing is significantly decreased in tottering$^{tg/tg}$ mice after NE application. **(H)** The CV1 of complex spikes tends to be increased, while the CV2 is not altered **(I)**. **(J)** Course of the mean complex spike firing of all PCs recorded shows that the firing frequency does not recover after NE application during the recording period of 400 sec. Data are presented as mean±SEM. Statistical significance was evaluated by paired t-test. (*$p \leq 0.05$, **$p \leq 0.01$, ***$p \leq 0.001$).

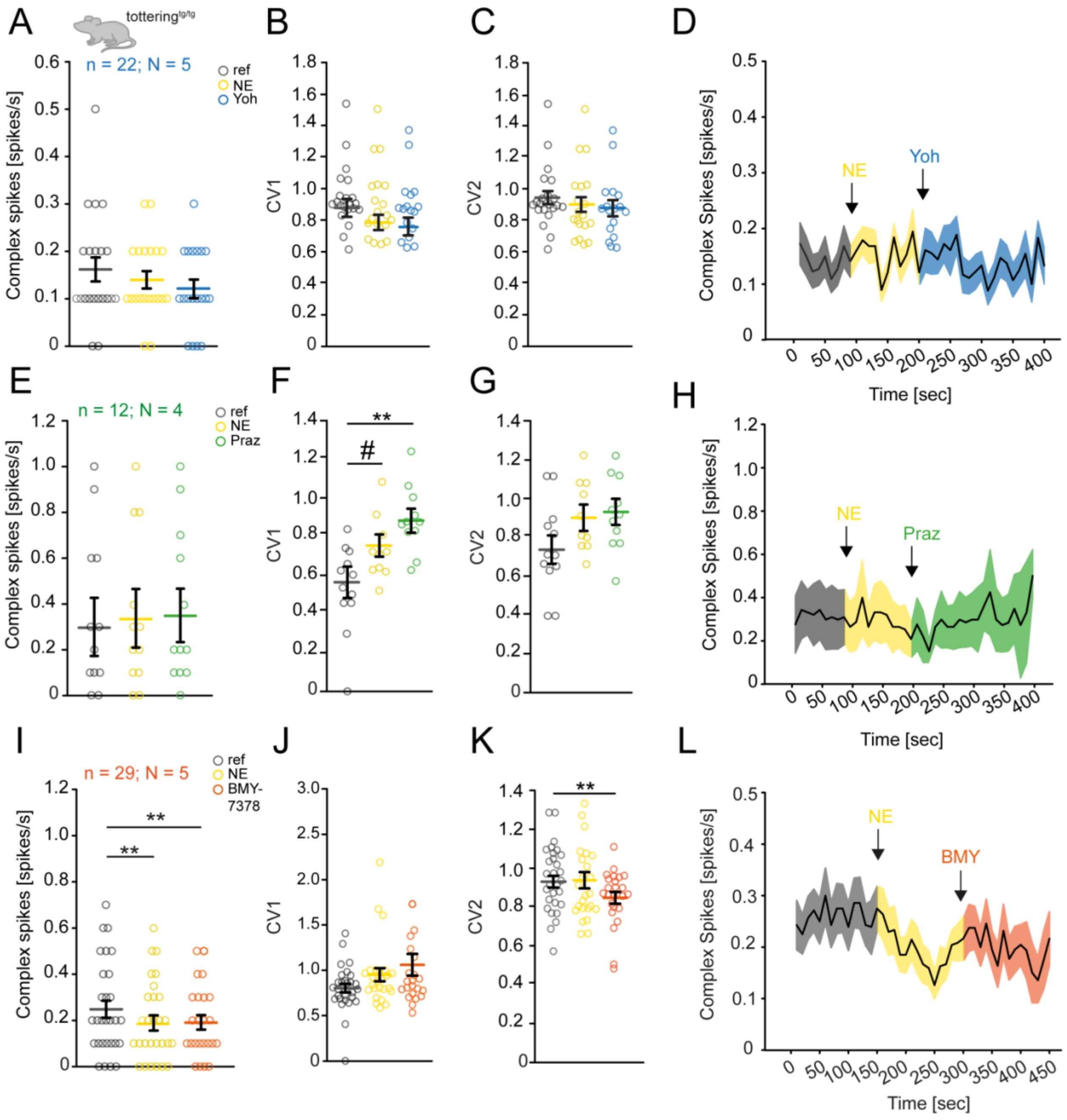


**Fig. S5. Effects of α1 and α2-AR antagonists on Purkinje cell complex spikes in tottering$^{tg/tg}$ mice *in vivo*.**

**(A)** Complex spike firing is neither further decreased, nor recovered after application of the α2-AR antagonist Yoh. **(B)** CV1 and CV2 **(C)** were not further altered after Yoh application following NE injection. **(D)** Course of the mean complex spike firing of all PCs recorded shows that the firing frequency is not altered after application of NE and Yoh. **(E)** Praz does not impact complex spike firing after NE application. **(F)** NE increased the CV1 of PCs, which was further increased by Praz application; however, the CV2 was not altered **(G)**. **(H)** Course of the mean complex spike firing of all PCs recorded shows a slight recovery after Praz application. **(I)** BMY-7378 does not rescue complex spike firing after NE mediated inhibition. **(J)** The CV1 of recorded PC complex spikes is not altered during drug application. **(K)** Application of BMY-7378 improved intrinsic complex spike variability. **(L)** Course of the mean Complex spike firing of all PCs recorded after NE and BMY-7378 application. Data are presented as mean±SEM. Statistical significance was evaluated by paired t-test. (*$p \leq 0.05$, **$p \leq 0.01$, ***$p \leq 0.001$).

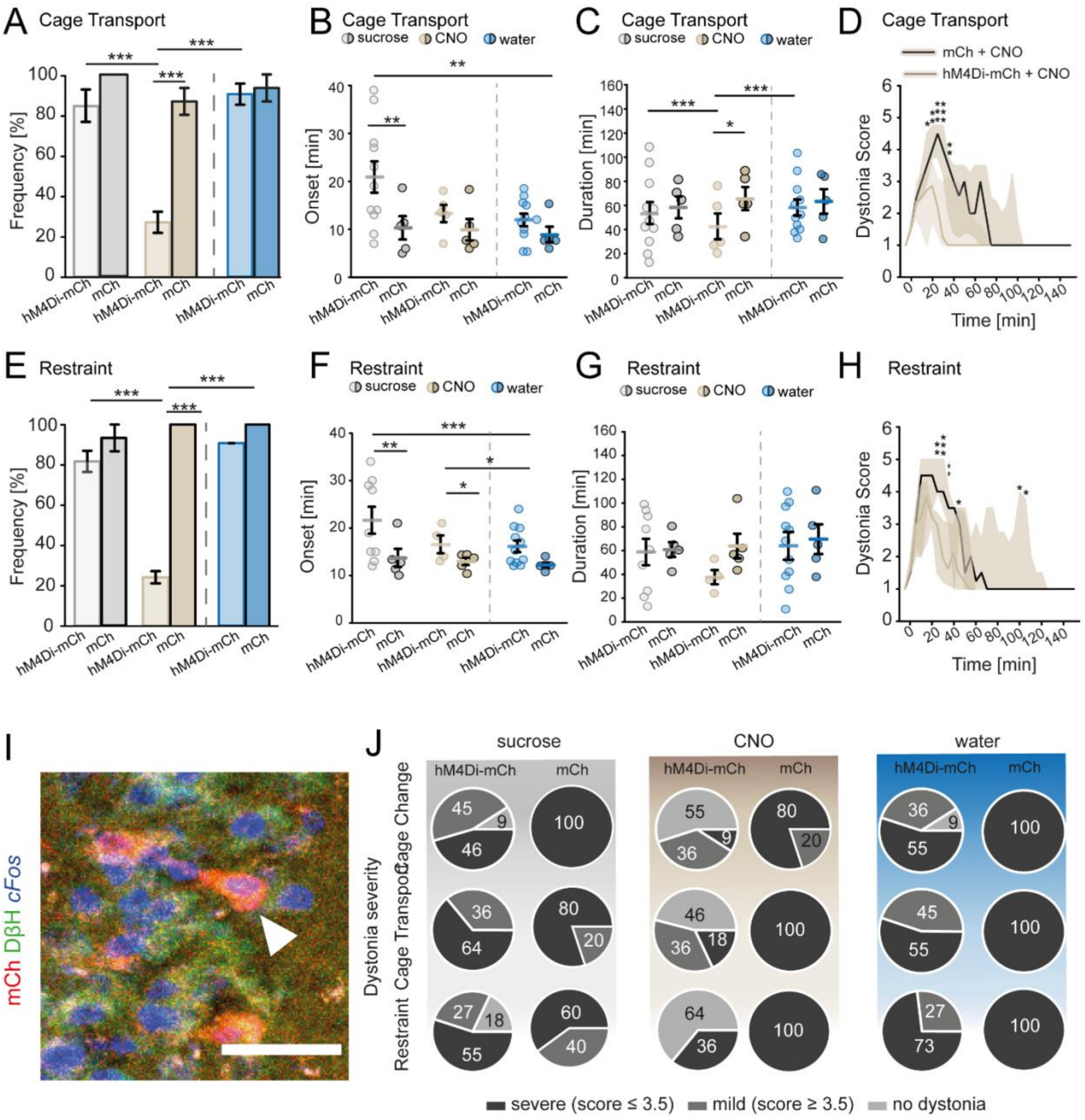


**Fig. S6. Oral administration of CNO reduces the frequency and severity of stress-induced dystonic attacks in the cage transport and restraint stress in DIO-hM4Di-mCherry, but not DIO-mCherry injected tottering$^{tg/tg}$/Th-Cre mice.**

**(A)** In the cage transport, DIO-hM4Di-mCh injected mice frequently displayed stress-induced dystonia when consuming sucrose (84.84±8.02%) and water (90.903±5.251%) compared to CNO treatment (27.27±5.248%, p≤0.001). **(B)** Attack onset was not different between groups, except for sucrose consuming DIO-hM4Di-mCh mice, showing delayed attacks (20.909±1.605

min). **(C)** Dystonia duration was reduced in DIO-hM4Di-mCh injected mice consuming CNO (21.05±4.993 min) compared to sucrose (53.455±2.984 min, $p \leq 0.001$) and water consumption (58.014±2.984, $p \leq 0.001$) or compared to control DIO-mCh injected mice consuming CNO (65.6±4.426, $p=0.016$). **(D)** Comparison of dystonia scores displays shorter attack duration and milder attack progression, if present, in DIO-hM4Di-mCh injected mice consuming CNO compared to DIO-mCh injected tottering$^{tg/tg}$ mice after a cage transport. **(E)** After a restraint, DIO-hM4Di-mCh injected tottering$^{tg/tg}$ mice displayed less dystonic attacks when consuming CNO (24.24±3.03%) compared to sucrose (81.547±5.019, $p \leq 0.001$) and water consumption (90.9±0, $p \leq 0.001$). Frequencies were not altered in DIO-mCh injected mice. **(F)** Onset of attacks was delayed in DREADD expressing mice compared to DIO-mCh injected mice (21.557±1.757 min vs 12.926±1.340 min, $p=0.034$). **(G)** Dystonia duration was reduced in DIO-hM4Di-mCh injected tottering$^{tg/tg}$ mice consuming CNO compared to control mice. **(H)** Dystonia severity is reduced in DIO-hM4Di-mCh injected tottering$^{tg/tg}$ compared to control mice. (**I)** Example image of mCh (red) infected LC neurons identified by DβH staining (green) and stained for c*Fos* (blue). Scale bars 25 µm. **(J)** mCh control mice display severe dystonic attacks in all stress tests and independent from the drinking agent. hM4Di-mCh injected mice show reduced severity and less attacks when drinking CNO, while severity is not altered when drinking sucrose alone or water. Data are presented as mean±SEM or median±75%/25% quartiles. Statistical significance was evaluated with Two Way Repeated Measure ANOVA, post-hoc Holm-Sidak all pairwise multiple comparison procedure (*$p \leq 0.05$, **$p \leq 0.01$, ***$p \leq 0.001$).

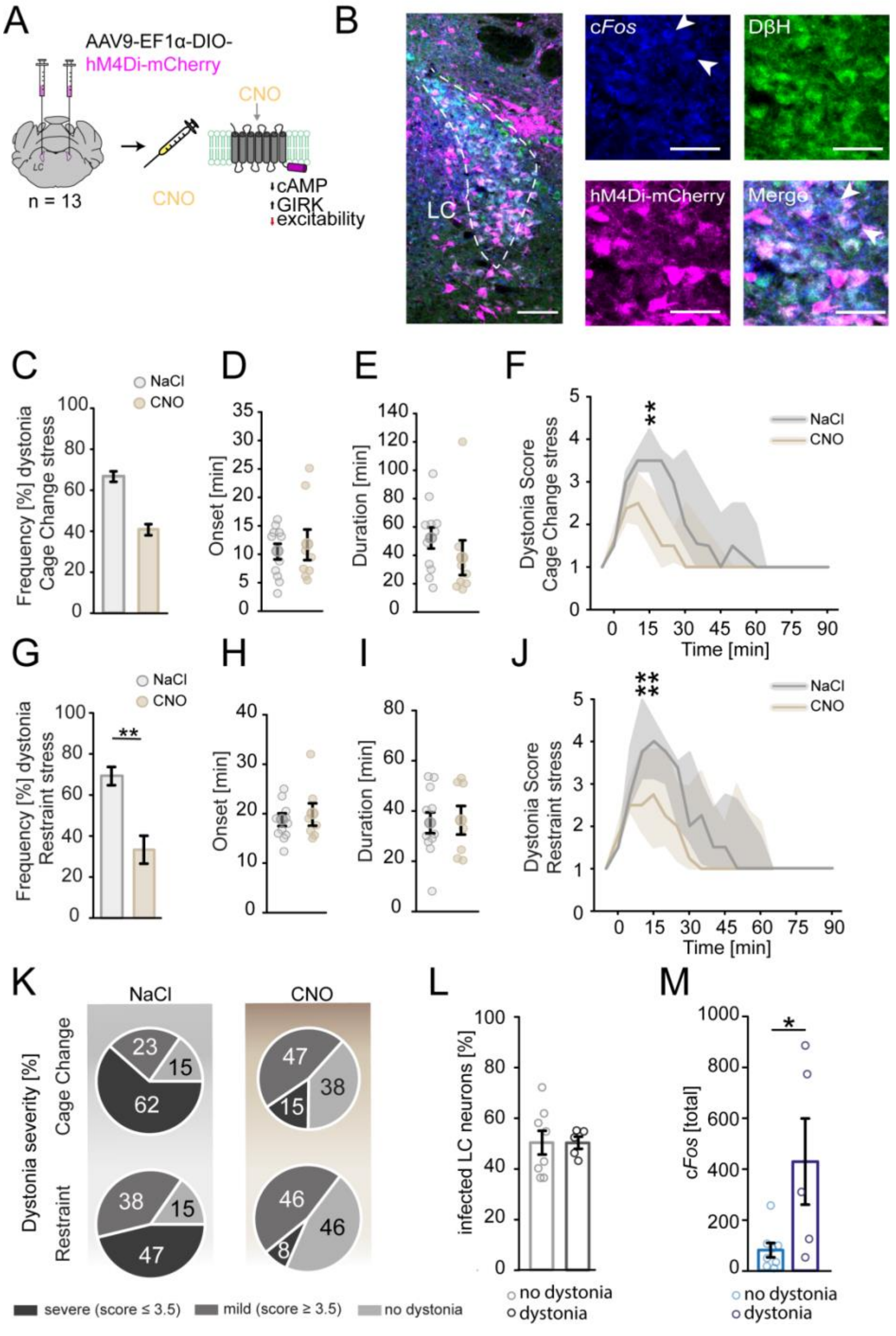


**Fig. S7. Single intraperitoneal injections of CNO are not as efficient in alleviating stress-induced paroxysmal dystonia compared to oral CNO delivery.**

**(A)** Tottering$^{tg/tg}$ mice were bilaterally injected with AAV9-EF1α-DIO-hM4Di-mCherry in the LC and ip injected with 10 mg/kg CNO to reduce neuronal excitability. **(B)** Example images of LC neurons infected with DIO-hM4Di-mCherry (magenta) identified via DβH staining (green), expressing cFos (blue). **(C)** Injection of CNO resulted in a trend of reducing the frequency of stress-induced dystonia compared to saline injected mice, while dystonia onset **(D)** and duration **(E)** were not affected in the cage change stress paradigm. **(F)** Comparison of the dystonia score showed trends in a reduction of dystonia severity for CNO injected mice. **(G)** CNO injected mice showed a significant alleviation of stress-induced dystonia in the restraint stress paradigm. **(H)** Onset of dystonia and duration **(I)** were not affected. **(J)** Dystonia scores were not significantly different between NaCl or CNO injected mice in the restraint test. **(K)** Comparison of dystonia severity showed that CNO injection leads to less severe attacks in mice undergoing dystonia and to fewer mice having an attack for both stress tests. **(L)** Some injected DIO-hM4Di-mCherry mice displayed stress-induced dystonia, but there was no difference in infected LC neurons compared to those mice which had no dystonic attack. **(M)** cFos counts of these dystonic mice revealed significantly more cFos expressing LC neurons compared to those not undergoing an attack. Data are presented as mean±SEM, except (F) and (J) are presented as median+25%/75% percentiles. Statistical significance was evaluated with student's t-test or Two Way repeated measure ANOVA. (*$p \leq 0.05$, **$p \leq 0.01$, ***$p \leq 0.001$).

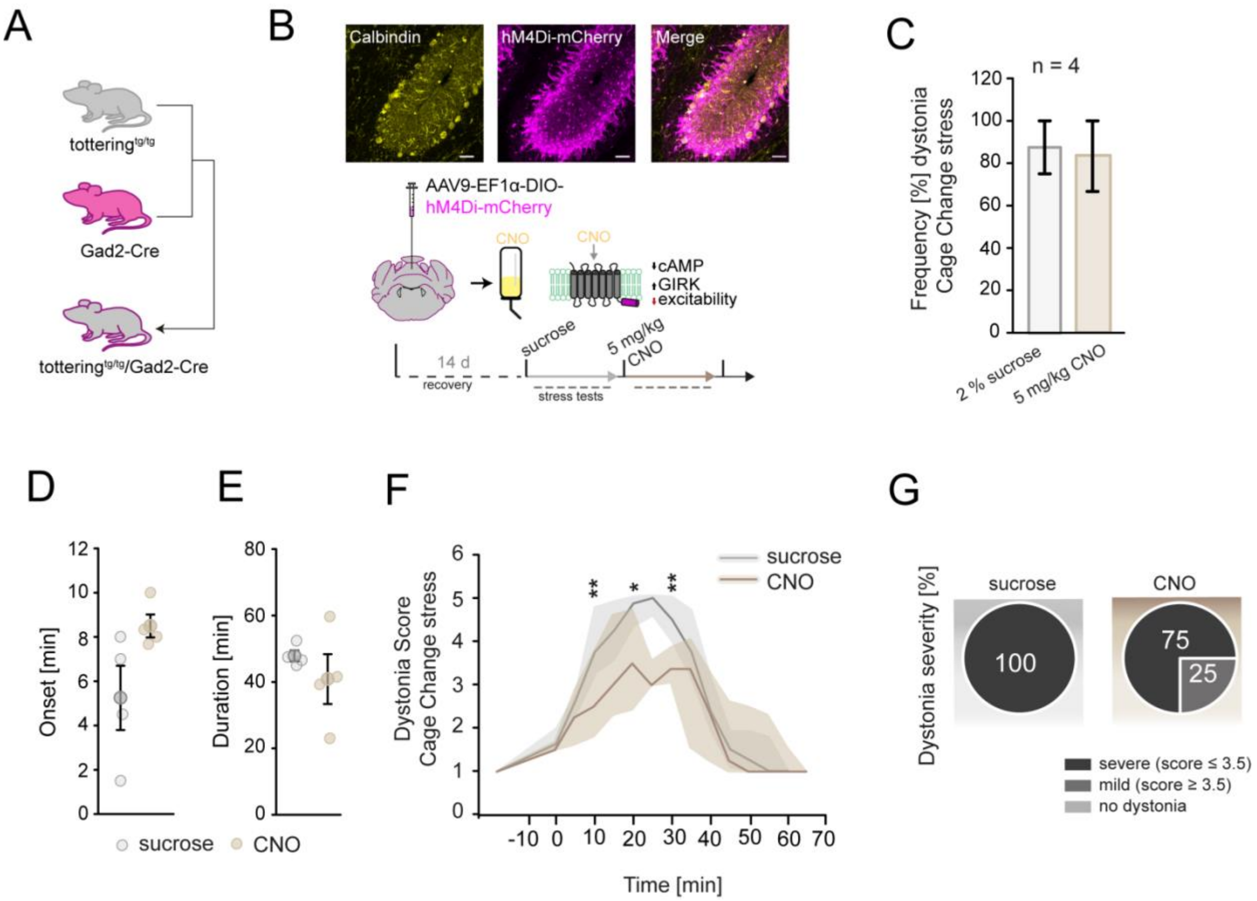


**Fig. S8. Chemogenetic silencing of cerebellar interneurons does not alleviate stress-induced paroxysmal dystonia in tottering$^{tg/tg}$/Gad2-Cre mice.**

**(A)** tottering$^{tg/tg}$ were bred with Gad2-Cre mice to obtain tottering$^{tg/tg}$/Gad2-Cre mice to allow interneuron (INs)specific Cre expression. **(B)** Top: Example images showing viral expression of AAV9-EF1α-DIO-hM4Di-mCherry in cerebellar MLIs. Scale bar: 50 µm. Bottom: Scheme of the testing paradigm. **(C)** Silencing of DIO-hM4Di-mCherry expressing vermal INs via CNO did not alleviate stress-induced dystonia, but showed a trend to delay its onset **(D)** and reduce its duration (**E**). **(F)** There was no significant difference for the dystonia score in DIO-hM4Di-mCherry injected mice for sucrose compared to CNO administration. **(G)** Comparison of dystonia severity showed no difference for the different drinks. Data are presented as mean±SEM, except **(F)** is presented as median+25%/75% percentiles. Statistical significance was evaluated with student's t-test or Two Way repeated measure ANOVA. (*p≤0.05, **p≤0.01, ***p≤0.001).

| Test | Sample Size | Statistics | mean±SEM | | p-value |
|---|---|---|---|---|---|
| | | | NaCl | Prazosin | |
| **Pole test** | n=10 | MW Rank Sum Test | 103.95±11.445 | 110.15±9.85 | p=0.584 |
| **Hang wire** | n=11 | Student's t-test | 15.212±4.787 | 26.364±5.129 | p=0.128 |
| **beam walk** | | | | | |
| *time (s)* | n=10 | MW Rank Sum test | 112.933±7.067 | 102.033±9.898 | p=0.280 |
| *idle (s)* | n=10 | MW Rank Sum test | 0 | 2.167±1.138 | **p=0.035** |
| *slips right HP (n)* | n=10 | MW Rank Sum test | 0.667±0.667 | 2.15±1.247 | p=0.180 |
| *slips left HP (n)* | n=10 | MW Rank Sum test | 0.267±0.267 | 1.45±0.914 | p=0.280 |
| **Footprint Analysis** | | | | | |
| *length right front paw (cm)* | n=11 | Student's t-test | 6.602±0.178 | 6.685±0.188 | p=0.751 |
| *length left front paw (cm)* | n=11 | MW Rank Sum test | 6.605±0.154 | 6.636±0.16 | p=0.921 |
| *length right hind paw (cm)* | n=11 | Student's t-test | 6.477±0.18 | 6.59±0.213 | p=0.883 |
| *length left hind paw (cm)* | n=11 | MW Rank Sum test | 6.155±0.284 | 6.319±0.287 | p=0.669 |
| *width front paws (cm)* | n=11 | MW Rank Sum test | 1.686±0.1223 | 1.685±0.135 | p=0.921 |
| *width hind paws (cm)* | n=11 | MW Rank Sum test | 3.126±0.179 | 3.165±0.167 | p=0.921 |

**Table S1. Adrenoreceptor blocker prazosin hydrochloride has no impact on ataxia in tottering $^{tg/tg}$ mice. Significant p-values are in bold.**

| Parameter | Statistics | Virus | mean±SEM 2 % sucrose vs. CNO | p-value |
|---|---|---|---|---|
| **total distance (cm)** | Student's t-test | DIO-hM4Di-mCh | 2729.364±361.65 vs. 2535.354±487.179 | p=0.752 |
| | Student's t-test | DIO-mCh | 2713.992±585.78 vs. 2628.739±328.404 | p=0.902 |
| **time spent in center (min)** | Student's t-test | DIO-hM4Di-mCh | 169.212±22.623 vs. 159.645±16.222 | p=0.367 |
| | Student's t-test | DIO-mCh | 152.438±26.835 vs. 173.148±30.795 | p=0.630 |
| **time at border (min)** | Student's t-test | DIO-hM4Di-mCh | 430.799±22.623 vs. 440.5±16.184 | p=0.732 |
| | Student's t-test | DIO-mCh | 447.563±26.835 vs. 426.852±30.795 | p=0.630 |

**Table S2. A high dose of 10 mg/kg CNO has no impact on locomotor activity in tottering$^{tg/tg}$ mice in the Open Field Test.**

| Analysis | Sample Size | Statistics | mean±SEM (Ref vs 2 mM BMY) | p-values |
|---|---|---|---|---|
| Simple Spikes/sec | N = 4; n = 16 | Wilcoxon Signed Rank Test | 34.525±2.926 vs 31.013±3.256 | p=0.002 |
| CV1 SS | | Wilcoxon Signed Rank Test | 0.618±0.0791 vs 0.702±0.0883 | p=0.159 |
| CV2 SS | | Wilcoxon Signed Rank Test | 0.504±0.0448 vs 0.541±0.0386 | p=0.175 |
| Complex Spikes (count) | N = 4; n = 16 | Paired t-test | 64.125±9.66 vs 56.438±7.674 | p=0.058 |
| CV1 CS | | Wilcoxon Signed Rank Test | 0.825±0.0389 vs 0.814±0.052 | p=0.464 |
| CV2 CS | | Paired t-test | 0.91±0.0441 vs 0838±0.0324 | p=0.129 |

**Table S3. Pressure microinjection of 2mM BMY-7378 does impair PC SS firing**

| **Primary Antibody** | **Dilution** | **Diluted in** | **supplier** |
|---|---|---|---|
| rb-α-α1A | 1:400 | blocking buffer | Invitrogen™, #PA1-047 |
| rb-α-α1B | 1:200 | blocking buffer | alomone labs, #AAR-018 |
| rb-α-α1D | 1:200 | blocking buffer | alomone labs, #AAR-019 |
| rb-α-α2A | 1:200 | blocking buffer | Invitrogen™, #PA1-048 |
| rb-α-α2B | 1:200 | blocking buffer | alomone labs, #AAR-021 |
| rb-α-α2C | 1:200 | blocking buffer | Invitrogen™, #PA5-114828 |
| ms-α-Actin Ab-5 | 1:10000 | blocking buffer | BD Biosciences, #612656 |
| ms-α-Calbindin | 1:500 | blocking buffer | Sigma-Aldrich, C9848 |
| **Secondary Antibodies** | | | |
| goat α-rb HRP-linked | 1:50000 | 1% milk powder, 0.5% BSA in 0.1% TBS-T | GE healthcare, NA934V5 |
| goat α-ms HRP-linked | 1:50000 | 1% milk powder, 0.5% BSA in 0.1% TBS-T | GE healthcare, NA931V5 |

**Table S4. Primary and secondary antibodies used for Western Blot analysis.**

| Primer name | Sequence |
|---|---|
| **ADR α1$_A$ fw** | 5’- TGGCTGCCATTCTTCCTCGTGA -3’ |
| **ADR α1$_A$ rev** | 5’- TTCTTGAACTCCTGGCTGGAGC -3’ |
| **ADR α1$_B$ fw** | 5’- CCTTGGGCATTGTAGTCGGA -3’ |
| **ADR α1$_B$ rev** | 5’- AAGTAGCCCAGCCAGAACAC -3’ |
| **ADR α1$_D$ fw** | 5’- CTGCCAAGACCCTAGCCATC -3’ |
| **ADR α1$_D$ rev** | 5’- GATGACCTTGAAGACGCCCT -3’ |
| **ADR α2$_A$ fw** | 5’- CAGGTGACACTGACGCTGGTTT -3’ |
| **ADR α2$_A$ rev** | 5’- GACACCAGGAAGAGGTTTTGGG -3’ |
| **ADR α2$_B$ fw** | 5’- GTTCCAGCCTCGGCTAAAGT -3’ |
| **ADR α2$_B$ rev** | 5’- GACCAATTGGGTGGCAAAGC -3’ |
| **ADR α2$_C$ fw** | 5’- CTTGATCTGGGCCTCGACTG -3’ |
| **ADR α2$_C$ rev** | 5’- GGTGCGCATCATTCCTTTGG -3’ |
| **HPRT fw** | 5’- AGTTCTTTGCTGACCTGCTG -3’ |
| **HRPT rev** | 5’- CCACCAATAACTTTTATGTCCCC -3’ |
| **ACVRL1 fw** | 5’- CCTAGTGCTATGGGAGATCGC -3’ |
| **ACVRL1 rev** | 5’- TGGGTGTCTGCTGGTCAAC -3’ |
| **GAPDH fw** | 5’- GCATTGTGGAAGGGCTCATG -3’ |
| **GAPDH rev** | 5’- TGCAGGGATGATGTTCTGGG -3’ |
| **PGK1 fw** | 5’- CCTGTTGACTTTGTCACTGC -3’ |
| **PGK1 rev** | 5’- CCCACAGCCTCGGCATATTT-3’ |
| **CALB1 fw** | 5’- ATTTCGACGCTGACGGAAGT -3’ |
| **CALB1 rev** | 5’- CCAATCCAGCCTTCTTTCGC -3’ |

**Table S5. Used quantitative Real-Time PCR primers.**

ACVRL1 = activin A receptor like type 1; ADR = adrenergic receptor; CALB1 = Calbindin 1, GDPDH = glyceraldehyde-3-phosphate dehydrogenase; HRPT = hypoxanthine guanine phosphoribosyl transferase; PGK1 = phosphoglycerate kinase 1.

| Figure | Sample Size | Statistics | | Comparison | %±SEM | p-value |
|---|---|---|---|---|---|---|
| 1A | n=11 | MW Rank Sum Test | frequency | NaCl vs Praz | 72.0±0 vs 9.0±0 | **p=0.003** |
| | | | | NaCl vs Yoh | 72.0±0 vs 63.6±0 | p=0.684 |
| | | | | Praz vs Yoh | 9.0±0 vs 63.6±0 | **p=0.011** |
| | | | | **comparison** | **mean±SEM** | **p-value** |
| | | | onset (min) | NaCl vs Praz | 8.75±2.418 vs 10.0 | not tested |
| | | Students t-test | | NaCl vs Yoh | 8.75±2.418 vs 5.286±1.426 | p=0.256 |
| | | | | Praz vs Yoh | 10.0 vs 5.286±1.426 | - |
| | | | | **comparison** | **mean±SEM** | **p-value** |
| | | | duration (min) | NaCl vs Praz | 37.875±7.11 vs 27.0 | - |
| | | Students t-test | | NaCl vs Yoh | 37.875±7.11 vs 39.857±7.049 | p=0.847 |
| | | | | Praz vs Yoh | 27.0 vs 39.857±7.049 | - |
| | | | | **comparison** | **%±SEM** | **p-value** |
| 1B | n=5 | MW Rank Sum Test | frequency | NaCl vs RS-17053 | 80.0 vs 80.0 | p=1.0 |
| | | Students t-test | onset (min) | NaCl vs RS-17053 | 7.5±3.304 vs 19.25±3.351 | **p=0.047** |
| | | Students t-test | duration (min) | NaCl vs RS-17053 | 74.75±12.195 vs 74.25±17.24 | p=0.982 |
| | | | | **comparison** | **mean±SEM** | **p-value** |
| 1C | n=15 | MW Rank Sum Test | frequency | NaCl vs BMY-7378 | 66.67 vs 13.3 | **p=0.004** |
| | | Students t-test | onset (min) | NaCl vs BMY-7378 | 48.8±8.612 vs 33.0±7.0 | p=0.451 |
| | | Students t-test | duration (min) | NaCl vs BMY-7378 | 6.1±1.516 vs 7.5±6.5 | p=0.745 |
| | | | **Comparison** | | **time** | **p-value** |
| 1D | n=5 | Two Way RM ANOVA post-hoc Holm-Sidak | NaCl vs Praz | | -15 | p=1.0 |
| | | | | | 0 | p=0.711 |
| | | | | | 5 | p=0.276 |
| | | | | | 10 | p=0.079 |
| | | | | | 15 | **p=0.031** |
| | | | | | 20 | **p=0.015** |
| | | | | | 25 | **p=0.019** |
| | | | | | 30 | **p=0.031** |
| | | | | | 35 | **p=0.04** |
| | | | | | 40 | **p=0.05** |
| | | | | | 45 | p=0.099 |
| | | | | | 50 | p=0.099 |
| | | | | | 55 | **p=0.05** |
| | | | | | 60 | p=0.079 |
| | | | | | 65 | p=0.099 |
| | | | | | 70 | p=0.187 |
| | | | | | 75 | p=0.539 |
| | | | | | 80 | p=0.462 |
| | | | | | 85 | p=0.539 |

| | | | | | |
|---|---|---|---|---|---|
| | | | | 90 | p=0.805 |
| | | | | 95 | p=0.901 |
| | | | | 100 | p=1.0 |
| | | | **Comparison** | **time** | **p-value** |
| 1D | N=5 | Two Way RM ANOVA post-hoc Holm-Sidak | NaCl vs BMY-7378 | -15 | p=1.0 |
| | | | | 0 | p=0.516 |
| | | | | 5 | p=0.079 |
| | | | | 10 | p=0.366 |
| | | | | 15 | **p=0.021** |
| | | | | 20 | **p=0.012** |
| | | | | 25 | **p=0.019** |
| | | | | 30 | **p=0.021** |
| | | | | 35 | **p=0.036** |
| | | | | 40 | **p=0.047** |
| | | | | 45 | **p=0.028** |
| | | | | 50 | **p=0.047** |
| | | | | 55 | **p=0.036** |
| | | | | 60 | p=0.061 |
| | | | | 65 | p=0.079 |
| | | | | 70 | p=0.162 |
| | | | | 75 | p=0.516 |
| | | | | 80 | p=0.437 |
| | | | | 85 | p=0.516 |
| | | | | 90 | p=0.794 |
| | | | | 95 | p=0.896 |
| | | | | 100 | p=1.0 |
| | **Sample size** | **Statistics** | **Comparison** | **%±SEM** | **p-value** |
| 1F | 10 vs 10 | Students t-test | $\alpha1_A$ tottering$^{tg/tg}$ vs tottering$^{-/-}$ | 5.031±0.254 vs.4.906±0.288 | p=0.564 |
| | | MW Rank Sum Test | $\alpha1_B$ tottering$^{tg/tg}$ vs tottering$^{-/-}$ | 7.805±0.866 vs.5.609±0.391 | **p=0.038** |
| | | MW Rank Sum Test | $\alpha1_D$ tottering$^{tg/tg}$ vs tottering$^{-/-}$ | 6.665±0.888 vs.4.273±0.434 | **p=0.045** |
| | | Students t-test | $\alpha2_A$ tottering$^{tg/tg}$ vs tottering$^{-/-}$ | 6.121±0.291 vs.5.894±0.298 | p=0.592 |
| | | Students t-test | $\alpha2_B$ tottering$^{tg/tg}$ vs tottering$^{-/-}$ | 8.841±0.225 vs.9.875±0.391 | **p=0.040** |
| | | Students t-test | $\alpha2_C$ tottering$^{tg/tg}$ vs tottering$^{-/-}$ | 6.923±0.288 vs.6.538±0.353 | p=0.410 |
| **Figure** | **Sample size** | **statistics** | **comparison** | **%±SEM** | **p-value** |
| 2G | n=22, N=5 | signed rank sum test | Ref vs NE | 25.014±2.318 vs 14.882±1.657 | **p≤0.001** |
| | | signed rank sum test | NE vs Yoh | 14.882±1.657 vs 13.464±1.642 | p=0.055 |
| | | paired t-test | Ref vs Yoh | 25.014±2.318 vs 13.464±1.642 | **p≤0.001** |
| 2H | n=22, N=5 | signed rank sum test | Ref vs NE | 4.623±1.33 vs 3.229±2.516 | p=0.284 |
| | | signed rank sum test | NE vs Yoh | 3.229±2.516 vs 3.241±0.622 | p=0.194 |
| | | signed rank sum test | Ref vs Yoh | 4.623±1.33 vs 3.241±0.622 | **p=0.044** |

| | | | | | |
|---|---|---|---|---|---|
| 2I | n=22, N=5 | paired t-test | Ref vs NE | 0.788±0.0468 vs 0.828±0.0304 | p=0.195 |
| | | signed rank sum test | NE vs Yoh | 0.828±0.0304 vs 0.868±0.0322 | p=0.098 |
| | | paired t-test | Ref vs Yoh | 0.788±0.0468 vs 0.868±0.0322 | **p=0.006** |
| 2K | n=12, N=4 | paired t-test | Ref vs NE | 23.325±4.281 vs 7.65±2.51 | **p≤0.001** |
| | | paired t-test | NE vs Praz | 7.65±2.51 vs 18.942±3.209 | **p≤0.001** |
| | | paired t-test | Ref vs Praz | 23.325±4.281 vs 18.942±3.209 | p=0.113 |
| 2L | n=12, N=4 | signed rank sum test | Ref vs NE | 2.123±0.22 vs 2.574±0.476 | p=0.85 |
| | | paired t-test | NE vs Praz | 2.574±0.476 vs 2.388±0.288 | p=0.741 |
| | | paired t-test | Ref vs Praz | 2.123±0.22 vs 2.388±0.288 | p=0.291 |
| 2M | n=12, N=4 | paired t-test | Ref vs NE | 0.866±0.0681 vs 0.977±0.0524 | p=0.114 |
| | | paired t-test | NE vs Praz | 0.977±0.0524 vs 0.860±0.0657 | p=0.034 |
| | | paired t-test | Ref vs Praz | 0.866±0.0681 vs 0.860±0.0657 | p=0.860 |
| 2O | n=29, N=5 | paired t-test | Ref vs NE | 30.203±1.94 vs 17.486±1.711 | **p≤0.001** |
| | | paired t-test | NE vs BMY-7378 | 17.486±1.711 vs 22.031±1.77 | **p=0.013** |
| | | signed rank sum test | Ref vs BMY-7378 | 30.203±1.94 vs 22.031±1.77 | **p≤0.001** |
| 2P | n=29, N=5 | signed rank sum test | Ref vs NE | 1.892±0.367 vs 2.449±0.512 | p=0.210 |
| | | signed rank sum test | NE vs BMY-7378 | 2.449±0.512 vs 1.786±0.363 | p=0.125 |
| | | signed rank sum test | Ref vs BMY-7378 | 1.892±0.367 vs 1.786±0.363 | p=1 |
| 2Q | n=29, N=5 | paired t-test | Ref vs NE | 0.7±0.0249 vs 0.772±0.0268 | **p=0.002** |
| | | signed rank sum test | NE vs BMY-7378 | 0.772±0.0268 vs 0.693±0.0362 | **p=0.015** |
| | | signed rank sum test | Ref vs BMY-7378 | 0.7±0.0249 vs 0.693±0.0362 | p=0.067 |
| **Figure** | **Sample size** | **statistics** | **comparison** | **%±SEM** | **p-value** |
| 3B | 5 vs 5 | Two Way RM ANOVA p≤0.001 post-hoc Holm-Sidak | frequency within ACSF pre vs. 5d | 93.33±6.667 vs 80.0 | p=0.749 |
| | | | frequency within ACSF pre vs. 7d | 93.33±6.667 vs 80.0 | p=0.684 |
| | | | frequency within ACSF pre vs. 10d | 93.33±6.667 vs 80.0 | p=0.842 |
| | | | frequency within ACSF pre vs. 14d | 93.33±6.667 vs 100.0 | p=1.0 |
| | | | frequency within ACSF 5d vs. 7d | 80.0 vs 80.0 | p=1.0 |
| | | | frequency within ACSF | 80.0 vs 80.0 | p=1.0 |

| | | | | | |
|---|---|---|---|---|---|
| | | | 5d vs. 10d | | |
| | | | frequency within ACSF<br>5d vs 14d | 80.0 vs 100.0 | p=0.9 |
| | | | frequency within ACSF<br>7d vs 10d | 80.0 vs 80.0 | p=1.0 |
| | | | frequency within ACSF<br>7d vs 14d | 80.0 vs 100.0 | p=0.875 |
| | | | frequency within ACSF<br>10d vs 14d | 80.0 vs 100.0 | p=0.801 |
| | | | frequency within BMY-7378<br>pre vs. 5d | 86.667±6.667 vs 40.0 | **p=0.003** |
| | | | frequency within BMY-7378<br>pre vs. 7d | 86.667±6.667 vs 0.0 | **p≤0.001** |
| | | | frequency within BMY-7378<br>pre vs. 10d | 86.667±6.667 vs 0.0 | **p≤0.001** |
| | | | frequency within BMY-7378<br>pre vs. 14d | 86.667±6.667 vs. 0.0 | **p≤0.001** |
| | | | frequency within BMY-7378<br>5d vs. 7d | 40.0 vs 0.0 | p=0.084 |
| | | | frequency within BMY-7378<br>5d vs. 10d | 40.0 vs 0.0 | p=0.057 |
| | | | frequency within BMY-7378<br>5d vs 14d | 40.0 vs 0.0 | p=0.071 |
| | | | frequency within BMY-7378<br>7d vs 10d | 0.0 vs 0.0 | p=1.0 |
| | | | frequency within BMY-7378<br>7d vs 14d | 0.0 vs 0.0 | p=1.0 |
| | | | frequency within BMY-7378<br>10d vs 14d | 0.0 vs 0.0 | p=1.0 |
| | | | frequency within pre<br>ACSF vs BMY-7378 | 93.33±6.667 vs 86.667±6.667 | p=1.0 |
| | | | frequency within 5d<br>ACSF vs BMY-7378 | 80.0 vs 40.0 | **p=0.044** |
| | | | frequency within 7d<br>ACSF vs BMY-7378 | 80.0 vs 0.0 | **p≤0.001** |
| | | | frequency within 10d<br>ACSF vs BMY-7378 | 80.0 vs 0.0 | **p≤0.001** |
| | | | frequency within 14d<br>ACSF vs BMY-7378 | 100.0 vs 0.0 | **p≤0.001** |
| | **Sample size** | **statistics** | **comparison** | **mean±SEM** | **p-value** |
| 3C | 5 | Two Way RM ANOVA p≤0.001 post-hoc Holm-Sidak | onset (min) within ACSF<br>pre vs. 5d | 11.4±2.775 vs. 6.25±2.175 | p=0.987 |
| | | | onset (min) within ACSF<br>pre vs. 7d | 11.4±2.775 vs. 7.25±2.323 | p=0.989 |
| | | | onset (min) within ACSF<br>pre vs. 10d | 11.4±2.775 vs. 4.75±2.25 | p=0.884 |
| | | | onset (min) within ACSF<br>pre vs. 14d | 11.4±2.775 vs. 4.0±1.095 | p=0.435 |
| | | | onset (min) within ACSF<br>5d vs. 7d | 6.25±2.175 vs. 7.25±2.323 | p=0.976 |
| | | | onset (min) within ACSF<br>5d vs. 10d | 6.25±2.175 vs. 4.75±2.25 | p=0.988 |
| | | | onset (min) within ACSF<br>5d vs 14d | 6.25±2.175 vs. 4.0±1.095 | p=0.245 |

| | | | | | |
|---|---|---|---|---|---|
| | | | onset (min) within ACSF 7d vs 10d | 7.25±2.323 vs. 4.75±2.25 | p=0.993 |
| | | | onset (min) within ACSF 7d vs 14d | 7.25±2.323 vs. 4.0±1.095 | p=0.180 |
| | | | 1 onset (min) within ACSF 0d vs 14d | 4.75±2.25 vs. 4.0±1.095 | p=0.380 |
| | 5 | Two Way RM ANOVA p≤0.001 post-hoc Holm-Sidak | onset (min) within BMY-7378 pre vs 5d | 8.766±1.268 vs. 20.0±5.0 | **p≤0.001** |
| | | | onset (min) within pre ACSF vs BMY-7378 | 11.4±2.775 vs. 8.766±1.268 | p=0.678 |
| 3D | 5 | Two Way RM ANOVA p≤0.001 post-hoc Holm-Sidak | duration (min) within ACSF pre vs. 5d | 57.365±4.809 vs. 45.25±5.851 | **p=0.003** |
| | | | duration (min) within ACSF pre vs. 7d | 57.365±4.809 vs. 48.5±6.198 | **p=0.012** |
| | | | duration (min) within ACSF pre vs. 10d | 57.365±4.809 vs. 44.75±5.558 | **p=0.003** |
| | | | duration (min) within ACSF pre vs. 14d | 57.365±4.809 vs. 49.0±5.933 | p=0.417 |
| | | | duration (min) within ACSF 5d vs. 7d | 45.25±5.851 vs. 48.5±6.198 | p=0.862 |
| | | | duration (min) within ACSF 5d vs. 10d | 45.25±5.851 vs. 44.75±5.558 | p=0.941 |
| | | | duration (min) within ACSF 5d vs. 14d | 45.25±5.851 vs. 49.0±5.933 | p=0.126 |
| | | | duration (min) within ACSF 7d vs. 10d | 48.5±6.198 vs. 44.75±5.558 | p=0.924 |
| | | | duration (min) within ACSF 7d vs. 14d | 48.5±6.198 vs. 49.0±5.933 | p=0.284 |
| | | | duration (min) within ACSF 10d vs.14d | 44.75±5.558 vs. 49.0±5.933 | p=0.124 |
| | 5 | Two Way RM ANOVA p≤0.001 post-hoc Holm-Sidak | duration (min) within BMY-7378 pre vs. 5d | 55.632±5.593 vs. 12.5±6.5 | **p≤0.001** |
| | | | duration (min) within BMY-7378 pre vs. 7d | 55.632±5.593 vs. 0.0 | **p≤0.001** |
| | | | duration (min) within BMY-7378 pre vs. 10d | 55.632±5.593 vs. 0.0 | **p≤0.001** |
| | | | duration (min) within BMY-7378 pre vs. 14d | 55.632±5.593 vs. 0.0 | **p≤0.001** |
| | | | duration (min) within BMY-7378 5d vs. 7d | 12.5±6.5 vs. 0.0 | p=0.928 |
| | | | duration (min) within BMY-7378 5d vs. 10d | 12.5±6.5 vs. 0.0 | p=0.827 |
| | | | duration (min) within BMY-7378 5d vs. 14d | 12.5±6.5 vs. 0.0 | p=0.888 |
| | | | duration (min) within BMY-7378 7d vs. 10d | 0.0 vs. 0.0 | p=1.0 |
| | | | duration (min) within BMY-7378 7d vs. 14d | 0.0 vs. 0.0 | p=1.0 |
| | | | duration (min) within BMY-7378 10d vs.14d | 0.0 vs. 0.0 | p=1.0 |
| | | | duration (min) within BMY-7378 17d (post 3d) | 42.0±1.871 | - |
| | | | duration (min) within BMY-7378 | 37.2±1.655 | - |

| | | | | | |
|---|---|---|---|---|---|
| | | | 20d (post 7d) | | |
| | | | duration (min) within pre ACSF vs BMY-7378 | 57.365±4.809 vs. 55.632±5.593 | p=0.853 |
| | | | duration (min) within 5d ACSF vs BMY-7378 | 45.25±5.851 vs. 12.5±6.5 | **p=0.004** |
| | | | duration (min) within 7d ACSF vs BMY-7378 | 48.5±6.198 vs. 0.0 | **p≤0.001** |
| | | | duration (min) within 10d ACSF vs BMY-7378 | 44.75±5.558 vs. 0.0 | **p=0.002** |
| | | | duration (min) within 14d ACSF vs BMY-7378 | 49.0±5.933 vs. 0.0 | **p≤0.001** |
| | **Sample size** | **Statistics** | **Comparison** | **time** | **p-value** |
| 3E | 5 vs 5 | Two Way RM ANOVA p≤0.001 post-hoc Holm-Sidack | 5d ACSF vs BMY-7378 | -15 | p=1 |
| | | | | 0 | p=1 |
| | | | | 5 | p=0.121 |
| | | | | 10 | **p=0.015** |
| | | | | 15 | **p≤0.001** |
| | | | | 20 | **p≤0.001** |
| | | | | 25 | **p≤0.001** |
| | | | | 30 | **p≤0.001** |
| | | | | 35 | **p=0.004** |
| | | | | 40 | **p=0.002** |
| | | | | 45 | **p=0.002** |
| | | | | 50 | p=0.340 |
| | | | | 55 | p=1 |
| | | | | 60 | p=1 |
| 3F | 5 vs 5 | Two Way RM ANOVA p≤0.001 post-hoc Holm-Sidack | 7d ACSF vs BMY-7378 | -15 | p=1 |
| | | | | 0 | p=0.218 |
| | | | | 5 | **p=0.017** |
| | | | | 10 | **p≤0.001** |
| | | | | 15 | **p≤0.001** |
| | | | | 20 | **p≤0.001** |
| | | | | 25 | **p≤0.001** |
| | | | | 30 | **p≤0.001** |
| | | | | 35 | **p≤0.001** |
| | | | | 40 | **p=0.002** |
| | | | | 45 | p=0.069 |
| | | | | 50 | p=0.535 |
| | | | | 55 | p=0.535 |
| | | | | 60 | p=0.535 |
| 3G | 5 vs 5 | Two Way RM ANOVA p≤0.001 post-hoc Holm-Sidack | 10d ACSF vs BMY-7378 | -15 | p=1 |
| | | | | 0 | p=0.124 |
| | | | | 5 | **p≤0.001** |
| | | | | 10 | **p≤0.001** |
| | | | | 15 | **p≤0.001** |
| | | | | 20 | **p≤0.001** |
| | | | | 25 | **p≤0.001** |
| | | | | 30 | **p≤0.001** |
| | | | | 35 | **p≤0.001** |
| | | | | 40 | **p=0.024** |
| | | | | 45 | p=0.057 |
| | | | | 50 | p=0.245 |
| | | | | 55 | p=0.436 |

| | | | | | |
|---|---|---|---|---|---|
| | | | | 60 | p=1 |
| 3H | 5 vs 5 | Two Way RM ANOVA p≤0.001 post-hoc Holm-Sidack | 14d ACSF vs BMY-7378 | -15 | p=1 |
| | | | | 0 | p=0.094 |
| | | | | 5 | **p≤0.001** |
| | | | | 10 | **p≤0.001** |
| | | | | 15 | **p≤0.001** |
| | | | | 20 | **p≤0.001** |
| | | | | 25 | **p≤0.001** |
| | | | | 30 | **p≤0.001** |
| | | | | 35 | **p≤0.001** |
| | | | | 40 | **p=0.001** |
| | | | | 45 | **p=0.018** |
| | | | | 50 | p=0.094 |
| | | | | 55 | p=0.467 |
| | | | | 60 | p=0.467 |
| | | | | 65 | p=0.808 |
| **Figure** | **Sample size** | **statistics** | **Comparison** | **%±SEM** | **p-value** |
| 4C | 10 | Two Way RM ANOVA p≤0.001 post-hoc Holm-Sidack | frequency within shRNA pre vs 3 weeks | 80.0±10.0 vs 20.0±0.0 | **p≤0.001** |
| | 10 | | frequency within shRNA pre vs 4 weeks | 80.0±10.vs 0.0 | **p≤0.001** |
| | 10 | | frequency within shRNA pre vs 5 weeks | 80.0±10.vs 0.0 | **p≤0.001** |
| | 10 | | frequency within shRNA pre vs 6 weeks | 80.0±10.vs 0.0 | **p≤0.001** |
| | 10 | | frequency within shRNA 3weeks vs 4 weeks | 20.0±0.0 vs 0.0 | **p=0.037** |
| | 10 | | frequency within shRNA 3weeks vs 5 weeks | 20.0±0.0 vs 0.0 | **p=0.031** |
| | 10 | | frequency within shRNA 3weeks vs 6 weeks | 20.0±0.0 vs 0.0 | **p=0.025** |
| | 10 | | frequency within shRNA 4 weeks vs 5 weeks | 0.0 vs 0.0 | p=1 |
| | 10 | | frequency within shRNA 4 weeks vs 6 weeks | 0.0 vs 0.0 | p=1 |
| | 10 | | frequency within shRNA 5 weeks vs 6 weeks | 0.0 vs 0.0 | p=1 |
| | 5 | | frequency within scrambleRNA pre vs 3 weeks | 86.667±6.667 vs 100 | p=1 |
| | 5 | | frequency within scrambleRNA pre vs 4 weeks | 86.667±6.667 vs 100 | p=1 |
| | 5 | | frequency within scrambleRNA pre vs 5 weeks | 86.667±6.667 vs 100 | p=1 |
| | 5 | | frequency within scrambleRNA pre vs 6 weeks | 86.667±6.667 vs 100 | p=1 |
| | 5 | | frequency within scrambleRNA 3 weeks vs 4 weeks | 100 vs 100 | p=1 |
| | 5 | | frequency within scrambleRNA 3 weeks vs 5 weeks | 100 vs 100 | p=1 |
| | 5 | | frequency within scrambleRNA 3 weeks vs 6 weeks | 100 vs 100 | p=1 |
| | 5 | | frequency within scrambleRNA 4 weeks vs 5 weeks | 100 vs 100 | p=1 |

| | | | | | |
|---|---|---|---|---|---|
| | 5 | | frequency within scrambleRNA 4 weeks vs 6 weeks | 100 vs 100 | p=1 |
| | 5 | | frequency within scrambleRNA 5 weeks vs 6 weeks | 100 vs 100 | p=1 |
| | 10 vs 5 | | frequency within pre | 80.0±10.0 vs 86.667±6.667 | p=1 |
| | 10 vs 5 | | frequency within 3 weeks | 20.0±0.0 vs 100 | **p≤0.001** |
| | 10 vs 5 | | frequency within 4 weeks | 0 vs 100 | **p≤0.001** |
| | 10 vs 5 | | frequency within 5 weeks | 0 vs 100 | **p≤0.001** |
| | 10 vs 5 | | frequency within 6 weeks | 0 vs 100 | **p≤0.001** |
| | **Sample size** | **statistics** | **Comparison** | **mean±SEM** | **p-value** |
| 4D | 10 vs 5 | Two Way RM ANOVA p=0.838 post-hoc Holm-Sidack | not tested | | |
| 4E | 10 | Two Way RM ANOVA p=0.019 post-hoc Holm-Sidack | duration (min) within shRNA pre vs 3 weeks | 67.816±7.782 vs 40.0±17 | **p≤0.001** |
| | 10 | | duration (min) within shRNA pre vs 4 weeks | 67.816±7.782 vs 0.0 | **p≤0.001** |
| | 10 | | duration (min) within shRNA pre vs 5 weeks | 67.816±7.782 vs 0.0 | **p≤0.001** |
| | 10 | | duration (min) within shRNA pre vs 6 weeks | 67.816±7.782 vs 0.0 | **p≤0.001** |
| | 10 | | duration (min) within shRNA 3weeks vs 4 weeks | 40.0±17 vs 0.0 | p=0.907 |
| | 10 | | duration (min) within shRNA 3weeks vs 5 weeks | 40.0±17 vs 0.0 | p=0.942 |
| | 10 | | duration (min) within shRNA 3weeks vs 6 weeks | 40.0±17 vs 0.0 | p=0.851 |
| | 10 | | duration (min) within shRNA 4 weeks vs 5 weeks | 0.0 vs 0.0 | p=1 |
| | 10 | | duration (min) within shRNA 4 weeks vs 6 weeks | 0.0 vs 0.0 | p=1 |
| | 10 | | duration (min) within shRNA 5 weeks vs 6 weeks | 0.0 vs 0.0 | p=1 |
| | 5 | | duration (min) within scrambleRNA pre vs 3 weeks | 86.032±17.131 vs 79.4±20.825 | p=0.844 |
| | 5 | | duration (min) within scrambleRNA pre vs 4 weeks | 86.032±17.131 vs 49.6±5.316 | p=0.059 |
| | 5 | | duration (min) within scrambleRNA pre vs 5 weeks | 86.032±17.131 vs 52.0±6.465 | p=0.087 |
| | 5 | | duration (min) within scrambleRNA pre vs 6 weeks | 86.032±17.131 vs 15.255±6.822 | p=0.254 |
| | 5 | | duration (min) within scrambleRNA 3 weeks vs 4 weeks | 79.4±20.825 vs 49.6±5.316 | p=0.171 |
| | 5 | | duration (min) within scrambleRNA 3 weeks vs 5 weeks | 79.4±20.825 vs 52.0±6.465 | p=0.227 |
| | 5 | | duration (min) within scrambleRNA 3 weeks vs 6 | 79.4±20.825 vs 15.255±6.822 | p=0.524 |

| | | | | | |
|---|---|---|---|---|---|
| | | | weeks | | |
| | 5 | | duration (min) within scrambleRNA 4 weeks vs 5 weeks | 49.6±5.316 vs 52.0±6.465 | p=0.851 |
| | 5 | | duration (min) within scrambleRNA 4 weeks vs 6 weeks | 49.6±5.316 vs 15.255±6.822 | p=0.878 |
| | 5 | | duration (min) within scrambleRNA 5 weeks vs 6 weeks | 52.0±6.465 vs 15.255±6.822 | p=0.891 |
| | 10 vs 5 | | duration (min) within pre | 67.816±7.782 vs 86.032±17.131 | p=0.98 |
| | 10 vs 5 | | duration (min) within 3 weeks | 40.0±17 vs 79.4±20.825 | **p≤0.001** |
| | 10 vs 5 | | duration (min) within 4 weeks | 0.0 vs 49.6±5.316 | **p≤0.001** |
| | 10 vs 5 | | duration (min) within 5 weeks | 0.0 vs 52.0±6.465 | **p≤0.001** |
| | 10 vs 5 | | duration (min) within 6 weeks | 0.0 vs 15.255±6.822 | **p≤0.001** |
| | **Sample size** | **statistics** | **Comparison** | **time** | **p-value** |
| 4F | 10 vs 5 | Two Way RM ANOVA p≤0.001 post-hoc Holm-Sidack | week 3 post injection shRNA vs scrambleRNA | -15 | p=1 |
| | | | | 0 | **p≤0.001** |
| | | | | 5 | **p≤0.001** |
| | | | | 10 | **p≤0.001** |
| | | | | 15 | **p≤0.001** |
| | | | | 20 | **p≤0.001** |
| | | | | 25 | **p≤0.001** |
| | | | | 30 | **p≤0.001** |
| | | | | 35 | **p≤0.001** |
| | | | | 40 | **p≤0.001** |
| | | | | 45 | **p≤0.001** |
| | | | | 50 | **p≤0.001** |
| | | | | 55 | **p=0.002** |
| | | | | 60 | **p=0.009** |
| | | | | 65 | **p=0.009** |
| | | | | 70 | **p=0.009** |
| | | | | 75 | **p=0.009** |
| | | | | 80 | **p=0.009** |
| | | | | 85 | p=0.097 |
| | | | | 90 | p=0.315 |
| | | | | 95 | p=0.182 |
| | | | | 100 | **p=0.047** |
| | | | | 105 | p=0.097 |
| | | | | 110 | p=0.182 |
| | | | | 115 | p=0.182 |
| | | | | 120 | p=1 |
| | | | | 125 | p=0.182 |
| | **Sample** | **statistics** | **Comparison** | **mean±SEM** | **p-value** |

| | | | | | | |
|---|---|---|---|---|---|---|
| | **size** | | | | | |
| 4G | 9 vs 5 | Student’s t-test | ADRA1D shRNA vs scrambleRNA | | 0±0.0903 vs 0.343±0.0572 | p=0.006 |
| 4I | 9 vs 5 | Student’s t-test | ADRA1D shRNA vs scrambleRNA | | 0.521±0.137 vs 0.916±0.110 | p=0.007 |
| | 2 | - | tottering$^{tg/tg}$ | | 0.734±.0682 | - |
| **Figure** | **Sample size** | **statistics** | **Comparison** | | **mean±SEM** | **p-value** |
| 5E | 204 NaCl vs 205 BMY | Wilcoxon Signed Rank Test | ROI firing | NaCl pre vs during | 3.843±0.0639 vs 3.901±0.0716 | p=0.317 |
| | | Paired t-test | | NaCl pre vs post | 3.843±0.0639 vs 3.877±0.0616 | p=0.587 |
| | | Wilcoxon Signed Rank Test | | NaCl during vs post | 3.901±0.0716 vs 3.877±0.0616 | p=0.706 |
| | | Wilcoxon Signed Rank Test | | BMY pre vs during | 3.222±0.0653 vs 3.039±0.0666 | **p≤0.001** |
| | | Wilcoxon Signed Rank Test | | BMY pre vs post | 3.222±0.0653 vs 3.097±0.668 | p=0.211 |
| | | Wilcoxon Signed Rank Test | | BMY during vs post | 3.097±0.668vs 3.097±0.668 | p=0. 191 |
| | | MW Rank Sum test | | pre NaCl vs BMY | 3.843±0.0639 vs 3.222±0.0653 | **p≤0.001** |
| | | Student’s t-test | | during NaCl vs BMY | 3.901±0.0716 vs 3.097±0.668 | **p≤0.001** |
| | | Student’s t-test | | post NaCl vs BMY | 3.877±0.0616 vs 3.097±0.668 | **p≤0.001** |
| 5G | | Wilcoxon Signed Rank Test | ΔF/F | NaCl pre vs during | 0.489±0.0315 vs 0.586±0.0453 | **p≤0.001** |
| | | Wilcoxon Signed Rank Test | | NaCl pre vs post | 0.489±0.0315 vs 0.515±0.0465 | p=0.085 |
| | | Wilcoxon Signed Rank Test | | NaCl during vs post | 0.586±0.0453 vs 0.515±0.0465 | **p≤0.001** |
| | | Wilcoxon Signed Rank Test | | BMY pre vs during | 0.336±0.0231 vs 0.261±0.0159 | **p≤0.001** |
| | | Wilcoxon Signed Rank Test | | BMY pre vs post | 0.336±0.0231 vs 0.228±0.0147 | **p≤0.001** |
| | | Wilcoxon Signed Rank Test | | BMY during vs post | 0.261±0.0159 vs 0.228±0.0147 | p=0.091 |
| | | MW Rank Sum test | | pre NaCl vs BMY | 0.489±0.0315 vs 0.336±0.0231 | **p≤0.001** |
| | | MW Rank Sum test | | during NaCl vs BMY | 0.586±0.0453 vs 0.261±0.0159 | **p≤0.001** |
| | | MW Rank Sum test | | post NaCl vs BMY | 0.515±0.0465 vs 0.228±0.0147 | **p≤0.001** |

| | | | | | | |
|---|---|---|---|---|---|---|
| 5I | 204 NaCl vs 205 BMY | Wilcoxon Signed Rank Test | Peak Height | NaCl pre vs during | 0.0133±0.000698 vs 0.0156±0.000994 | **p=0.004** |
| | | Wilcoxon Signed Rank Test | | NaCl pre vs post | 0.0133±0.000698 vs 0.013±0.000912 | **p=0.003** |
| | | Wilcoxon Signed Rank Test | | NaCl during vs post | 0.0156±0.000994 vs 0.013±0.000912 | **p≤0.001** |
| | | Wilcoxon Signed Rank Test | | BMY pre vs during | 0.011±0.000497 vs 0.00957±0.000369 | **p≤0.001** |
| | | Wilcoxon Signed Rank Test | | BMY pre vs post | 0.011±0.000497 vs 0.00888±0.000359 | **p≤0.001** |
| | | Wilcoxon Signed Rank Test | | BMY during vs post | 0.00957±0.000369 vs 0.00888±0.000359 | **p=0.012** |
| | | MW Rank Sum test | | pre NaCl vs BMY | 0.0133±0.000698 vs 0.011±0.000497 | **p≤0.001** |
| | | MW Rank Sum test | | during NaCl vs BMY | 0.0156±0.000994 vs 0.00957±0.000369 | **p≤0.001** |
| | | MW Rank Sum test | | post NaCl vs BMY | 0.013±0.000912 vs 0.00888±0.000359 | p=0.173 |
| **Figure** | **Sample size** | **statistics** | **Comparison** | | **%±SEM** | **p-value** |
| 6C | 11 | Two Way RM ANOVA p=0.002 post-hoc Holm-Sidack | frequency within DIO-hM4Di-mCh | Sucrose vs CNO | 90.9±0 vs 27.24±9.12 | **p≤0.001** |
| | | | | Water vs CNO | 89.873±6.063 vs 27.24±9.12 | **p≤0.001** |
| | | | | Sucrose vs water | 90.9±0 vs 89.873±6.063 | p=0.57 |
| | 5 | | frequency within DIO-mCh | Sucrose vs CNO | 93.33±6.667 vs 100±0 | p=0.965 |
| | | | | Water vs CNO | 100±0 vs 100±0 | p=1.0 |
| | | | | Sucrose vs water | 93.33±6.667 vs 100±0 | p=0.893 |
| | 11 vs 5 | | frequency within CNO | DIO-hM4Di-mCh vs DIO-mCh | 27.24±9.12 vs 100±0 | **p≤0.001** |
| | 11 vs 5 | | frequency within sucrose | DIO-hM4Di-mCh vs DIO-mCh | 90.9±0 vs 93.33±6.667 | p=0.863 |
| | 11 vs 5 | | frequency within water | DIO-hM4Di-mCh vs DIO-mCh | 87.873±6.063 vs 100±0 | p=0.284 |
| | | | | | **mean±SEM** | **p-value** |
| 6D | 11 | Two Way RM ANOVA p=0.008 | onset (min) within DIO-hM4Di-mCh | Sucrose vs CNO | 14.03±1.941 vs 23.962±3.01 | **p=0.023** |
| | | | | Water vs CNO | 9.029±1.941 vs 23.962±3.01 | **p=0.001** |

| | | | | | | |
|---|---|---|---|---|---|---|
| | | post-hoc Holm-Sidack | | Sucrose vs water | 14.03±1.941 vs 9.029±3.01 | p=0.083 |
| | 5 | | onset (min) within DIO-mCh | Sucrose vs CNO | 7.193±2.626 vs 6.733±2.626 | p=0.903 |
| | | | | Water vs CNO | 4.133±2.626 vs 6.733±2.626 | p=0.742 |
| | | | | Sucrose vs water | 7.193±2.626 vs 4.133±2.626 | p=0.805 |
| | | | | | **Diff. of mean** | **p-value** |
| | 11 vs 5 | | onset (min) within CNO | DIO-hM4Di-mCh vs DIO-mCh | 17.230 | **p=0.002** |
| | 11 vs 5 | | onset (min) within sucrose | DIO-hM4Di-mCh vs DIO-mCh | 6.837 | p=0.103 |
| | 11 vs 5 | | onset (min) within water | DIO-hM4Di-mCh vs DIO-mCh | 4.895 | p=0.238 |
| | | | | | **mean±SEM** | **p-value** |
| 6E | 11 | Two Way RM ANOVA p=0.021 post-hoc Holm-Sidack | duration (min) within DIO-hM4Di-mCh | Sucrose vs CNO | 65.525±6.873 vs 27.436±10.658 | **p=0.021** |
| | | | | Water vs CNO | 50.746±6.874 vs 27.436±10.658 | p=0.155 |
| | | | | Sucrose vs water | 65.525±6.873 vs 50.746±6.873 | p=0.144 |
| | 5 | | duration (min) within DIO-mCh | Sucrose vs CNO | 54.733±9.298 vs 72.133±9.298 | p=0.489 |
| | | | | Water vs CNO | 64.913±9.298 vs 72.133±9.298 | p=0.695 |
| | | | | Sucrose vs water | 54.733±9.298 vs 64.913±9.298 | p=0.589 |
| | | | | | **Diff. of mean** | **p-value** |
| | 11 vs 5 | | duration (min) within CNO | DIO-hM4Di-mCh vs DIO-mCh | 44.697 | **p=0.025** |
| | 11 vs 5 | | duration (min) within sucrose | DIO-hM4Di-mCh vs DIO-mCh | 10.793 | p=0.488 |
| | 11 vs 5 | | duration (min) within water | DIO-hM4Di-mCh vs DIO-mCh | 14.167 | p=0.364 |
| | **Sample size** | **Statistics** | **Comparison** | | **time** | **p-value** |
| 6F | 11 vs 5 | Two Way RM ANOVA p≤0.001 post-hoc Holm-Sidack | DIO-hM4Di-mCh vs DIO-mCh | | -15 | p=1.0 |
| | | | | | 0 | p=0.922 |
| | | | | | 5 | p=0.175 |
| | | | | | 10 | **p=0.008** |
| | | | | | 15 | **p=0.013** |
| | | | | | 20 | p=0.244 |
| | | | | | 25 | p=0.244 |
| | | | | | 30 | p=0.082 |
| | | | | | 35 | p=0.207 |

| | | | | | |
|---|---|---|---|---|---|
| | | | | 40 | p=0.101 |
| | | | | 45 | **p=0.021** |
| | | | | 50 | **p=0.022** |
| | | | | 55 | p=0.244 |
| | | | | 60 | p=0.770 |
| | | | | 65 | p=0.495 |
| | | | | 70 | p=0.626 |
| | | | | 75 | p=0.559 |
| | | | | 80 | p=0.922 |
| | | | | 85 | p=0.696 |
| | | | | 90 | p=1.0 |
| | | | | 95 | p=0.845 |
| | | | | 100 | p=0.626 |
| | | | | 105 | p=0.696 |
| | | | | 110 | p=0.770 |
| | | | | 115 | p=0.770 |
| | | | | 120 | p=0.770 |
| | | | | 125 | p=0.770 |
| | | | | 130 | p=0.845 |
| | | | | 135 | p=0.845 |
| | **Sample size** | | **Analysis** | **mean±SEM** | |
| 6H | 11 | | Dopamine-β-hydroxylase$^+$ neurons | 3084.818±93 | |
| | | | DIO-hM4Di-mCh$^+$ neurons | 2456.909±97.617 | |
| | | | Dopamine-β-hydroxylase$^+$/ DIO-hM4Di-mCh$^+$ neurons | 1899.909±86.377 | |
| | **Sample Size** | **Statistics** | **Comparison** | **percentage±SEM** | **p-value** |
| 6I | 11 | | % of Dopamine-β-hydroxylase$^+$/ DIO-hM4Di-mCh$^+$ neurons | 62.078±3.136 | |
| | **Sample Size** | **Statistics** | **Comparison** | **mean±SEM** | **p-value** |
| 6J | 11 vs 5 | MW Rank Sum test | DIO-hM4Di-mCh vs DIO-mCh | 53.091±16.47 vs 1059.8±107.213 | p=0.002 |

**Table S6. List of statistical tests, p-values and data.** Significant p-values are highlighted in bold.

| | Sample Size | Statistics | Comparison | mean±SEM | p-values |
|---|---|---|---|---|---|
| S3B | n=44, N=6 tottering$^{-/-}$ n=33, N=4 tottering$^{tg/tg}$ | MW Rank Sum test | Simple Spikes | 30.289±1.657 vs 26.43±2.778 | p=0.142 |
| | | MW Rank Sum test | CV1 | 0.71±0.0769 vs 1.765±0.216 | **p≤0.001** |
| | | MW Rank Sum test | CV2 | 0.470±0.0179 vs 0.728±0.0285 | **p≤0.001** |
| S3C | n=44, N=6 tottering$^{-/-}$ n=33, N=4 tottering$^{tg/tg}$ | MW Rank Sum test | Complex Spikes | 0.232±0.0245 vs 0.145±0.0169 | **p=0.012** |
| | | MW Rank Sum test | CV1 | 0.855±0.0349 vs 0.895±0.0406 | p=0.53 |
| | | Student's t-test | CV2 | 0.857±0.015 vs 0.901±0.0233 | p=0.108 |
| S4C | N = 3; n = 34 | Paired t-test | Simple Spikes Reference vs NE | 30.146±1.697 vs 16.340±1.996 | **p≤0.001** |
| | N = 3; n = 34 | Wilcoxon Signed Rank Test | CV1 Reference vs NE | 1.356±0.218 vs 3.261±1.085 | p=0.242 |
| | N = 3; n = 34 | Wilcoxon Signed Rank Test | CV2 Reference vs NE | 0.632±0.0242 vs 0.732±0.0303 | **p≤0.001** |
| | N = 3; n = 34 | Wilcoxon Signed Rank Test | Complex Spikes Reference vs NE | 0.179±0.0183 vs 0.103±0.0171 | **p=0.003** |
| | N = 3; n = 34 | Paired t-test | CV1 Reference vs NE | 0.832±0.0304 vs 0.881±0.0681 | p=0.335 |
| | N = 3; n = 34 | Wilcoxon Signed Rank Test | CV2 Reference vs NE | 0.914±0.0305 vs 0.927±0.0418 | p=0.459 |
| S5A | n=22, N=5 | Wilcoxon Signed Rank Test | Ref vs NE | 0.162±0.0253 vs 0.14±0.0184 | p=0.542 |
| | | Wilcoxon Signed Rank Test | NE vs Yoh | 0.14±0.0184 vs 0.121±0.0196 | p=0.175 |
| | | Wilcoxon Signed Rank Test | Ref vs Yoh | 0.162±0.0253 vs 0.121±0.0196 | p=0.083 |
| S5B | n=22, N=5 | Paired t-test | Ref vs NE | 0.878±0.0565 vs 0.785±0.0483 | p=0.223 |
| | | Paired t-test | NE vs Yoh | 0.785±0.0483 vs 0.76±0.0564 | p=0.52 |
| | | Paired t-test | Ref vs Yoh | 0.878±0.0565 vs 0.76±0.0564 | p=0.082 |
| S5C | n=22, N=5 | Paired t-test | Ref vs NE | 0.942±0.0428 vs 0.902±0.0529 | p=0.724 |
| | | Paired t-test | NE vs Yoh | 0.902±0.0529 vs 0.878±0.0493 | p=0.680 |
| | | Paired t-test | Ref vs Yoh | 0.942±0.0428 vs 0.878±0.0493 | p=0.301 |
| S5E | n=12, N=4 | Paired t-test | Ref vs NE | 0.3±0.127 vs 0.338±0.128 | p=1 |
| | | Wilcoxon | NE vs Praz | 0.338±0.128 vs | p=0.688 |

| | | | | | | |
|---|---|---|---|---|---|---|
| | | Signed Rank Test | | | | |
| | | Paired t-test | Ref vs Praz | | 0.3±0.127 vs 0.35±0.117 | p=0.594 |
| S5F | n=12, N=4 | Paired t-test | Ref vs NE | | 0.564±0.0815 vs 0.753±0.0574 | p=0.059 |
| | | Paired t-test | NE vs Praz | | 0.753±0.0574 vs 0.882±0.0619 | p=0.114 |
| | | Paired t-test | Ref vs Praz | | 0.564±0.0815 vs 0.882±0.0619 | p=0.012 |
| S5G | n=12, N=4 | Paired t-test | Ref vs NE | | 0.736±0.0731 vs 0.9±0.0679 | p=0.086 |
| | | Paired t-test | NE vs Praz | | 0.9±0.0679 vs 0.931±0.0672 | p=0.956 |
| | | Paired t-test | Ref vs Praz | | 0.736±0.0731 vs 0.931±0.0672 | p=0.201 |
| S5I | n=29, N=5 | Wilcoxon Signed Rank Test | Ref vs NE | | 0.248±0.037 vs 0.189±0.033 | **p=0.007** |
| | | Wilcoxon Signed Rank Test | NE vs BMY-7378 | | 0.189±0.033 vs 0.192±0.0312 | p=0.104 |
| | | Wilcoxon Signed Rank Test | Ref vs BMY-7378 | | 0.248±0.037 vs 0.192±0.0312 | **p=0.003** |
| S5J | n=29, N=5 | Wilcoxon Signed Rank Test | Ref vs NE | | 0.803±0.047 vs 0.951±0.0726 | p=0.333 |
| | | Paired t-test | NE vs BMY-7378 | | 0.951±0.0726 vs 1.061±0.120 | p=0.192 |
| | | Wilcoxon Signed Rank Test | Ref vs BMY-7378 | | 0.803±0.047 vs 1.061±0.120 | p=0.494 |
| S5K | n=29, N=5 | Paired t-test | Ref vs NE | | 0.930±0.03 vs 0.938±0.0426 | p=0.877 |
| | | Paired t-test | NE vs BMY-7378 | | 0.938±0.0426 vs 0.848±0.0423 | p=0.482 |
| | | Paired t-test | Ref vs BMY-7378 | | 0.930±0.03 vs ± 0.848±0.0423 | **p=0.005** |
| | **Sample Size** | **Statistics** | **Comparison** | | **%±SEM** | **p-value** |
| S6A | 11 | Two Way RM ANOVA p=0.004 post-hoc Holm-Sidack | frequency within DIO-hM4Di-mCh | Sucrose vs CNO | 84.84±8.02 vs 27.27±5.248 | **p≤0.001** |
| | | | | Water vs CNO | 90.903±5.251 vs 27.27±5.248 | **p≤0.001** |
| | | | | Sucrose vs water | 84.84±8.02 vs 90.903±5.251 | p=0.730 |
| | 5 | | frequency within DIO-mCh | Sucrose vs CNO | 100±0 vs 86.667±6.667 | p=0.673 |
| | | | | Water vs CNO | 93.33±6.667 vs 86.667±6.667 | p=0.610 |
| | | | | Sucrose vs water | 100±0 vs 93.33±6.667 | p=0.848 |
| | 11 vs 5 | | frequency within CNO | DIO-hM4Di-mCh vs DIO-mCh | 27.27±5.248 vs 86.667±6.667 | **p≤0.001** |

| | | | | | | |
|---|---|---|---|---|---|---|
| | 11 vs 5 | | frequency within sucrose | DIO-hM4Di-mCh vs DIO-mCh | 84.84±8.02 vs 100±0 | p=0.240 |
| | 11 vs 5 | | frequency within water | DIO-hM4Di-mCh vs DIO-mCh | 90.903±5.251 vs 93.333±6.667 | p=0.670 |
| | | | | | **mean±SEM** | **p-value** |
| S6B | 11 | Two Way RM ANOVA p=0.034 post-hoc Holm-Sidack | onset (min) within DIO-hM4Di-mCh | Sucrose vs CNO | 20.909±1.605 vs 17.106±2.686 | p=0.237 |
| | | | | Water vs CNO | 11.969±1.605 vs 17.106±2.686 | p=0.217 |
| | | | | Sucrose vs water | 20.909±1.605 vs 11.969±1.605 | **p=0.002** |
| | 5 | | onset (min) within DIO-mCh | Sucrose vs CNO | 10.327±2.381 vs 9.937±2.381 | p=0.908 |
| | | | | Water vs CNO | 8.933±2.381 vs 9.937±2.381 | p=0.947 |
| | | | | Sucrose vs water | 10.327±2.381 vs 8.933±2.381 | p=0.968 |
| | | | | | **Diff. of mean** | **p-value** |
| | 11 vs 5 | | onset (min) within CNO | DIO-hM4Di-mCh vs DIO-mCh | 7.172 | p=0.133 |
| | 11 vs 5 | | onset (min) within sucrose | DIO-hM4Di-mCh vs DIO-mCh | 10.582 | p=0.008 |
| | 11 vs 5 | | onset (min) within water | DIO-hM4Di-mCh vs DIO-mCh | 3.036 | p=0.420 |
| | | | | | **mean±SEM** | **p-value** |
| S6C | 11 | Two Way RM ANOVA p≤0.001 post-hoc Holm-Sidack | duration (min) within DIO-hM4Di-mCh | Sucrose vs CNO | 53.455±2.984 21.05±4.993 | **p≤0.001** |
| | | | | Water vs CNO | 58.014±2.984 vs 21.05±4.993 | **p≤0.001** |
| | | | | Sucrose vs water | 53.455±2.984 vs 58.014±2.984 | p=0.292 |
| | 5 | | duration (min) within DIO-mCh | Sucrose vs CNO | 58.199±4.426 vs 65.6±4.426 | p=0.578 |
| | | | | Water vs CNO | 63.2±4.426 vs 65.6±4.426 | p=0.705 |
| | | | | Sucrose vs water | 58.199±4.426 vs 63.2±4.426 | p=0.678 |
| | | | | | **Diff. of mean** | **p-value** |
| | 11 vs 5 | | duration (min) within CNO | DIO-hM4Di-mCh vs DIO-mCh | 44.55 | **p=0.016** |
| | 11 vs 5 | | duration (min) within sucrose | DIO-hM4Di-mCh vs DIO-mCh | 4.745 | p=0.725 |

| | | | | | | |
|---|---|---|---|---|---|---|
| | 11 vs 5 | | duration (min) within water | DIO-hM4Di-mCh vs DIO-mCh | 5.186 | p=0.701 |
| | | | | | **%±SEM** | **p-value** |
| S6E | 11 | Two Way RM ANOVA p≤0.001 post-hoc Holm-Sidack | frequency within DIO-hM4Di-mCh | Sucrose vs CNO | 81.547±5.019 vs 24.24±3.03 | **p≤0.001** |
| | | | | Water vs CNO | 90.9±0 vs 24.24±3.03 | **p≤0.001** |
| | | | | Sucrose vs water | 81.547±5.019 vs 90.9±0 | p=0.291 |
| | 5 | | frequency within DIO-mCh | Sucrose vs CNO | 100±0 vs 100±0 | p=1.0 |
| | | | | Water vs CNO | 100±0 vs 100±0 | p=1.0 |
| | | | | Sucrose vs water | 100±0 vs100±0 | p=1.0 |
| | 11 vs 5 | | frequency within CNO | DIO-hM4Di-mCh vs DIO-mCh | ± vs 100±0 | **p≤0.001** |
| | 11 vs 5 | | frequency within sucrose | DIO-hM4Di-mCh vs DIO-mCh | ± vs 100±0 | p=0.090 |
| | 11 vs 5 | | frequency within water | DIO-hM4Di-mCh vs DIO-mCh | ± vs 100±0 | p=0.326 |
| | | | | | **mean±SEM** | **p-value** |
| S6F | 11 | Two Way RM ANOVA p=0.013 post-hoc Holm-Sidack | onset (min) within DIO-hM4Di-mCh | Sucrose vs CNO | 22.333±1.065 vs 21.557±1.757 | p=0.710 |
| | | | | Water vs CNO | 16.151±0.903 vs 21.557±1.757 | **p=0.026** |
| | | | | Sucrose vs water | 22.333±1.065 vs 16.151±0.903 | **p≤0.001** |
| | 5 | | onset (min) within DIO-mCh | Sucrose vs CNO | 13.66±1.34 vs 12.926±1.340 | p=0.703 |
| | | | | Water vs CNO | 12.134±1.340 vs 12.926±1.340 | p=0.898 |
| | | | | Sucrose vs water | 13.66±1.34 vs 12.134±1.34 | p=0.815 |
| | | | | | **Diff. of mean** | **p-value** |
| | 11 vs 5 | | onset (min) within CNO | DIO-hM4Di-mCh vs DIO-mCh | 8.631 | **p=0.034** |
| | 11 vs 5 | | onset (min) within sucrose | DIO-hM4Di-mCh vs DIO-mCh | 8.673 | **p=0.008** |
| | 11 vs 5 | | onset (min) within water | DIO-hM4Di-mCh vs DIO-mCh | 4.017 | p=0.164 |
| | | | | | **mean±SEM** | **p-value** |
| | | Two Way RM | duration | Sucrose vs CNO | 59.169±6.186 vs | n.s. |

| | | | | | | |
|---|---|---|---|---|---|---|
| S6G | 11 | ANOVA p=0.343 post-hoc Holm-Sidack | (min) within DIO-hM4Di-mCh | | 31.679±10.2 | |
| | | | | Water vs CNO | 63.985±5.244 vs 31.679±10.2 | n.s. |
| | | | | Sucrose vs water | 59.169±6.189 vs 63.985±5.244 | n.s. |
| | 5 | | duration (min) within DIO-mCh | Sucrose vs CNO | 60.8±7.779 vs 63.799±7.779 | n.s. |
| | | | | Water vs CNO | 69.599±7.779 vs 63.799±7.779 | n.s. |
| | | | | Sucrose vs water | 60.8±7.779 vs 69.599±7.779 | n.s. |

| | **Sample size** | **statistics** | **Comparison** | **time** | **p-value** |
|---|---|---|---|---|---|
| S6D | 11 vs 5 | Two Way RM ANOVA p = 0.002 post-hoc Holm-Sidack | DIO-hM4Di-mCh vs DIO-mCh | -15 | p=1.0 |
| | | | | 0 | p=0.910 |
| | | | | 5 | p=0.784 |
| | | | | 10 | p=0.336 |
| | | | | 15 | **p=0.016** |
| | | | | 20 | **p=0.005** |
| | | | | 25 | **p≤0.001** |
| | | | | 30 | **p≤0.001** |
| | | | | 35 | **p=0.019** |
| | | | | 40 | p=0.092 |
| | | | | 45 | p=0.16 |
| | | | | 50 | p=0.066 |
| | | | | 55 | p=0.262 |
| | | | | 60 | p=0.126 |
| | | | | 65 | p=0.061 |
| | | | | 70 | p=0.070 |
| | | | | 75 | p=0.249 |
| | | | | 80 | p=0.290 |
| | | | | 85 | p=0.440 |
| | | | | 90 | p=0.336 |
| | | | | 95 | p=0.086 |
| | | | | 100 | p=0.562 |
| | | | | 105 | p=1.0 |
| S6H | 11 vs 5 | Two Way RM ANOVA p≤0.001 post-hoc Holm-Sidack | DIO-hM4Di-mCh vs DIO-mCh | -15 | p=1.0 |
| | | | | 0 | p=0.963 |
| | | | | 5 | p=0.664 |
| | | | | 10 | p=0.451 |
| | | | | 15 | p=0.246 |
| | | | | 20 | p=0.086 |
| | | | | 25 | **p=0.006** |
| | | | | 30 | **p=0.001** |
| | | | | 35 | **p=0.004** |
| | | | | 40 | p=0.091 |
| | | | | 45 | **p=0.032** |
| | | | | 50 | p=0.552 |
| | | | | 55 | p=0.362 |
| | | | | 60 | p=0.104 |
| | | | | 65 | p=0.147 |
| | | | | 70 | p=0.072 |
| | | | | 75 | p=0.072 |
| | | | | 80 | p=0.204 |
| | | | | 85 | p=0.204 |

| | | | | | |
|---|---|---|---|---|---|
| | | | | 90 | p=0.308 |
| | | | | 95 | p=0.204 |
| | | | | 100 | p=0.032 |
| | | | | 105 | p=0.048 |
| | | | | 110 | p=0.465 |
| | | | | 115 | p=0.714 |
| | | | | 120 | p=0.714 |
| | | | | 125 | p=1.0 |
| | **Sample Size** | **Statistics** | **Comparison** | **mean±SEM** | **p-values** |
| S7C | 13 | Student's t-test | NaCl vs CNO | 66.66±2.564 vs 40.717±2.717 | p=0.119 |
| S7D | 13 | Student's t-test | NaCl vs CNO | 10.365±1.366 vs 11.561±2.868 | p=0.672 |
| S7E | 13 | MW Rank Sum Test | NaCl vs CNO | 52.257±7.437 vs 38.354±12.243 | p=0.107 |
| | **Sample size** | **statistics** | **Comparison** | **time** | **p-value** |
| S7F | 13 | Two Way RM ANOVA (p=0.354) post-hoc Holm-Sidak | NaCl vs CNO | -15 | p=1 |
| | | | | 0 | p=0.719 |
| | | | | 5 | p=1 |
| | | | | 10 | p=0.351 |
| | | | | 15 | **p=0.003** |
| | | | | 20 | p=0.199 |
| | | | | 25 | p=0.077 |
| | | | | 30 | p=0.253 |
| | | | | 35 | p=0.565 |
| | | | | 40 | p=0.430 |
| | | | | 45 | p=0.719 |
| | | | | 50 | p=0.283 |
| | | | | 55 | p=0.119 |
| | | | | 60 | p=0.199 |
| | | | | 65 | p=0.472 |
| | | | | 70 | p=0.518 |
| | | | | 75 | p=0.565 |
| | | | | 80 | p=0.565 |
| | | | | 85 | p=0.719 |
| | | | | 90 | p=0.666 |
| | **Sample Size** | **Statistics** | **Comparison** | **mean±SEM** | **p-values** |
| S7G | 13 | Student's t-test | NaCl vs CNO | 69.227±4.443 vs 33.333±6.782 | p=0.024 |
| S7H | 13 | Student's t-test | NaCl vs CNO | 18.788±1.283 vs 19.833±2.264 | p=0.670 |
| S7I | 13 | Student's t-test | NaCl vs CNO | 35.287±4.179 vs 36.309±5.691 | p=0.885 |
| | **Sample size** | **statistics** | **Comparison** | **time** | **p-value** |
| | | Two Way RM ANOVA (p=0.154) post-hoc Holm-Sidak | NaCl vs CNO | -15 | p=1 |
| | | | | 0 | p=0.750 |
| | | | | 5 | p=0.382 |
| | | | | 10 | **p=0.007** |
| | | | | 15 | p=0.004 |
| | | | | 20 | p=0.059 |
| | | | | 25 | p=0.234 |
| | | | | 30 | p=0.178 |

| | | | | | |
|---|---|---|---|---|---|
| S7J | 13 | | | 35 | p=0.811 |
| | | | | 40 | p=0.577 |
| | | | | 45 | p=0.426 |
| | | | | 50 | p=0.873 |
| | | | | 55 | p=0.936 |
| | | | | 60 | p=0.873 |
| | | | | 65 | p=0.524 |
| | | | | 70 | p=0.340 |
| | | | | 75 | p=0.205 |
| | | | | 80 | p=0.524 |
| | | | | 85 | p=0.524 |
| | | | | 90 | p=1 |
| | **Sample Size** | **Statistics** | **Comparison** | **mean±SEM** | **p-values** |
| S7L | 8 vs 5 | Student's t-test | no dystonia vs dystonia | 50.451±4.694 vs 50.424±2.4 | p=0.997 |
| S7M | 8 vs 5 | MW Rank Sum test | no dystonia vs dystonia | 82±28.293 vs 429.8±169.208 | **p=0.03** |
| S8C | 4 | Student's t-test | sucrose vs CNO | 87.5±12.5 vs 83.3±16.667 | p=0.537 |
| S8D | 4 | Student's t-test | sucrose vs CNO | 5.25±1.451 vs 8.5±0.518 | p=0.079 |
| S8E | 4 | Student's t-test | sucrose vs CNO | 47.875±1.625 vs 40.875±7.503 | p=0.397 |
| | **Sample size** | **statistics** | **Comparison** | **time** | **p-value** |
| S8F | 4 | Two Way RM ANOVA (p=0.029) post-hoc Holm Sidak | sucrose vs CNO | -15 | p=1 |
| | | | | 0 | p=0.813 |
| | | | | 5 | p=0.555 |
| | | | | 10 | p=0.009 |
| | | | | 15 | p=0.085 |
| | | | | 20 | p=0.032 |
| | | | | 25 | p=0.004 |
| | | | | 30 | p=0.053 |
| | | | | 35 | p=0.107 |
| | | | | 40 | p=0.723 |
| | | | | 45 | p=0.813 |
| | | | | 50 | p=0.813 |
| | | | | 55 | p=0.723 |
| | | | | 60 | p=0.723 |
| | | | | 65 | p=1 |

**Table S7. List of statistical tests, p-values and data.** Significant p-values are highlighted in bold.

**Movie S1.**
Tottering with sh-ADRA1D expression does not exhibit stress-induced dystonia.

**Movie S2.**
Tottering with sh-scramble expression at the onset of stress-induced dystonia displays motor deficiencies.

**Movie S3.**
Example in-vivo calcium recording of a homozygous tottering mouse experiencing dystonia after NaCl injection.

**Movie S4.**
Example in-vivo calcium recording of a homozygous tottering mouse without dystonia after BMY-7378 injection.
